# Two-photon microscopy of brain endothelial glycocalyx uncovers spatial heterogeneity, vesicular transport, and lectin-binding kinetics in the living brain

**DOI:** 10.64898/2025.12.15.694376

**Authors:** Oleg Zhukov, Nikolay P. Kutuzov, Teddy Groves, Mette Mathiesen Janiurek, Sally Dabelsteen, Anders Hay-Schmidt, Hans H. Wandall, Christoffer Goth, Brian DellaValle, Martin Lauritzen, Krzysztof Kucharz

## Abstract

The endothelial glycocalyx is a key regulator of cerebrovascular function and remains one of the most difficult structures to study *in vivo*. Here we uncover new structural and dynamical features of the brain endothelial glycocalyx using *in vivo* two-photon microscopy. We identified glycocalyx enrichment at endothelial junctions and arteriolar branch points, visualized its Vesicular transport in real-time, and found evidence for its compositional Variations along the arteriovenous axis. Fluorescence recovery after photobleaching revealed two distinct kinetics of wheat germ agglutinin binding, including a previously undescribed one. Finally, super-localization of the glycocalyx estimated glycocalyx thickness as 775±17 nm and 622±34 nm before and after enzymatic shedding, reconciling discrepancies between past optical and electron microscopy estimates. Together these findings establish the first miltiscale framework of glycocalyx distribution and heterogeneity, transport, and molecular interaction kinetics in the living brain.

## Introduction

Glycans, along with nucleic acids, proteins, and lipids are fundamental building blocks of cells. Glycans’ abundance, chemical diversity, and their wide range of physiological functions have motivated researchers to decipher the “glycocode” in order to better understand the physiology and pathophysiology of single cells, organs, and tissues^1^. Among these, the endothelial glycocalyx – a glycan-rich layer on the luminal surface of the endothelium – acts as a component of the blood-brain barrier^2–4^, regulates the trafficking of blood-borne immune cells into the brain parenchyma^5^, senses shear stress, and transmits molecular signals to the endothelium^6–8^.

Despite its importance, the glycocalyx remains poorly characterized due to several challenges. Glycans are among the most variable, complex, and difficult macromolecules to measure^9,10^, and profiling them in situ is challenged by a lack of selective probes^10^. *In vivo* imaging of microvascular glycocalyx is further constrained by its thickness (∼500 nm)^11,12^, close to the resolution limit of two-photon microscopy (TPM, ∼300–400 nm), the gold standard for *in vivo* brain imaging^13^. In addition, motion artifacts, such as vessel wall and center pulsations^14^, displace the glycocalyx by ∼500 nm during each cardiac cycle^15^, causing severe motion blur, which further confounds the measurements. As a result of these challenges, the reported glycocalyx thickness of microvessels in the brain varies from hundreds of nanometers to several microns^16–18^. Notably, *in vivo* estimates are often several-fold greater than those obtained by electron microscopy (EM)^11,19,20^, highlighting a persistent discrepancy between the methods^21^.

Because of these challenges, the glycocalyx is mostly studied *in vitro* or *ex vivo*. These approaches showed glycocalyx enrichment at endothelial cells (ECs) junctions^7,22,23^ and Vessel branching points^18,24^; Variation in chemical composition across the peripheral circulation^25^ and within single cultured ECs^22^; and responses to flow changes^7^ and ageing^11,25^. More recently, surface plasmon resonance-based methods have enabled *in situ* profiling of glycocalyx components, revealing cell-surface compositional differences^10,26^. In addition, kinetic modelling of lectin-glycocalyx interactions has provided binding/unbinding constants for lectin-binding motifs, helping to identify specific chemical groups on single-cell surfaces^26^. Metabolic labeling has enabled real-time monitoring of glycans in zebrafish embryos^27^—specifically sialoglycans in the brain and other organs *in vivo*^28^—though without the submicrometer resolution required to study the brain glycocalyx. *In vivo* data on the brain glycocalyx remain scarce^4,16,18,29–31^ and often inconsistent, particularly regarding glycocalyx distribution along the arteriovenous axis^16,19^.. The lack of high-resolution *in vivo* data and limited analytical tools continue to impede our understanding of the brain endothelial glycocalyx.

Here, we developed and applied three complementary approaches to characterize the glycocalyx in the living brains of anesthetized mice using TPM. First, dual-lectin labeling enabled profiling of single endothelial cells, revealing compositional variability along the arteriovenous axis, enrichment at vascular branch points, and dynamics within endothelial transport pathways. Second, kinetic modeling combined with fluorescence recovery after photobleaching and enzymatic treatment identified slow and fast WGA–glycocalyx interaction kinetics. Third, super-localization of the glycocalyx allowed measurement of its thickness with nanoscale precision *in vivo*. Together, these approaches provide the first *in vivo* insights into the spatial organization, transport pathways, and previously unrecognized binding interactions of the brain endothelial glycocalyx, redefining its structural and functional landscape in the living brain.

## Results

### Three-dimensional localization of the endothelial glycocalyx with two-photon microscopy

To establish the three-dimensional geometry of the vessel walls and the glycocalyx distribution, we analyzed the distribution of the glycocalyx, labeled with i.a.-injected Alexa Fluor 594-conjugated wheat germ agglutinin (WGA-AF), relative to the GFP-expressing endothelium in Tie2GFP mice^32^, using TPM (Fig. 1a–c). Single arteriolar ECs were elongated in the direction of the flow (Extended Data Fig. 1a,d,f), measuring 30–50 µm in length and 10–15 µm in width, consistent with previous reports^32^ and with the contours of ECs junctions observed *in vivo*^33^. The luminal surface of ECs deviated from circular shape due to their nuclei protruding into the lumen (Fig. 1d). As expected, the highest WGA-AF fluorescence co-localized with the luminal surface of the endothelium, following its shape (I_gcx_; Fig. 1d)^34^. Moving from the glycocalyx towards the vessel’s center, WGA-AF fluorescence first reached a local minimum, corresponding to WGA-AF in blood plasma (I_plsm_), in a 1–2-µm-wide region (Fig. 1e) void of red blood cells (RBCs)^35^, neglecting other blood cells constituting < 0.1% of all blood cells^36^. Fluorescence in the central part of blood vessels was, on average, higher than I_plsm_ due to additional fluorescence from WGA-AF bound to the glycocalyx of RBCs (I_RBC_) ^4,37^. In contrast to WGA-AF, when blood plasma was labeled with fluorescent dextrans, the vessel’s center exhibited the lowest fluorescence since dextrans do not label RBCs (Extended data Fig. 1bc).

**Figure 1.**
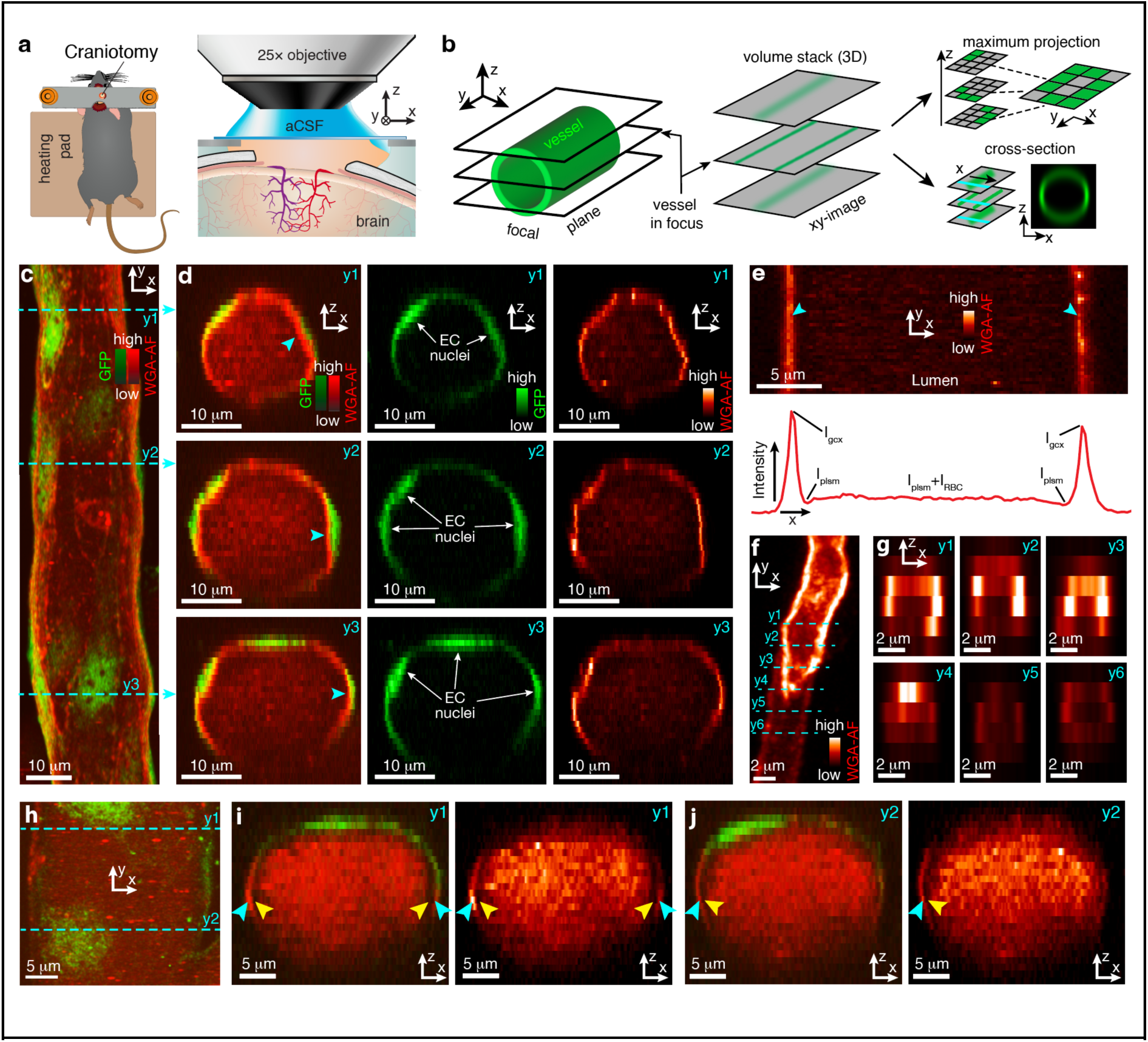
Two-photon imaging of the brain endothelial glycocalyx *in vivo*. **(a)** Experimental setup for two-photon imaging. **(b)** xy-images acquired at successive depths (z-axis; *left*) formed a (xyz)-volume stack (*middle*), from which we extracted (xz)*-*cross-sections (along cyan lines; *lower-right*) and maximum intensity projections (xy) to visualize brightest fluorescence in the volume on a single image (*upper-right*). **(c)** Maximum projection, showing a pial arteriole with WGA-AF-labeled glycocalyx and GFP-labeled endothelium. **(d)** Cross-sections (xz) of the arteriole at three y-locations, extracted along three cyan lines in (c), showing WGA-AF endothelial GFP (*left*), only GFP (*middle*), and only WGA-AF (*right*). Blue arrowheads indicate glycocalyx labeling on the luminal surface of endothelium. **(e)** *Top:* Image of an arteriole in focus, where two vertical bands of fluorescence represent the intersections of the focal plane with WGA-AF-labeled glycocalyx. *Bottom:* WGA-AF fluorescence intensity profile, obtained by averaging the image above along the y*-*axis. The profile highlights the glycocalyx (I_gcx_), RBC-free zone of lumen near the vessel wall (I_plsm_), and the central part of the lumen, where the fluorescence comes from the plasma and WGA-labeled RBCs (I_RBC_). **(f)** Maximum projection showing a capillary labeled with WGA-AF. **(g)** Cross-sections of the capillary at six locations (cyan lines in *left*), showing heterogeneous distribution of WGA-AF. **(h)** Maximum projection, showing a pial venule with less intense WGA-AF labeling of the glycocalyx, compared to the arteriole in (c). **(i,j)** Two cross-section (xz) of the venule (GFP + WGA-AF and only WGA-AF on the left and right subpanels, respectively), obtained by averaging the volume stack (xyz) over two 1-µm-tall y-ranges centered at y=y1 and y2, depicted as cyan lines in (h). Cyan and yellow arrowheads: I_gcx_ and I_plsm_, respectively.

### Glycocalyx heterogeneity on single endothelial cells

Having established the influence of vascular ECs geometry on the 3D glycocalyx distribution, we asked how the glycocalyx is distributed on individual ECs. Arterioles, venules, and capillaries demonstrated a heterogeneous distribution of WGA-AF fluorescence intensity (Fig. 1d-j), as contrasted by a uniformly-labeled model cylindrical surface (Extended Data Fig. 2gh). To investigate this heterogeneity, we imaged the top surfaces of pial arterioles, where we probed the glycocalyx fluorescence distribution with the xy-lateral resolution of 351±4 nm (3.7 times better than the axial resolution along the *z*-axis; Extended Data Fig. 2a-f). We identified increased WGA-AF fluorescence at the periphery of ECs, away from their nuclei (Fig. 2a-c), suggesting that glycocalyxes of contacting ECs coalesce to form a high-density glycocalyx layer at contact sites between ECs^7,22,23,38^.

**Figure 2.**
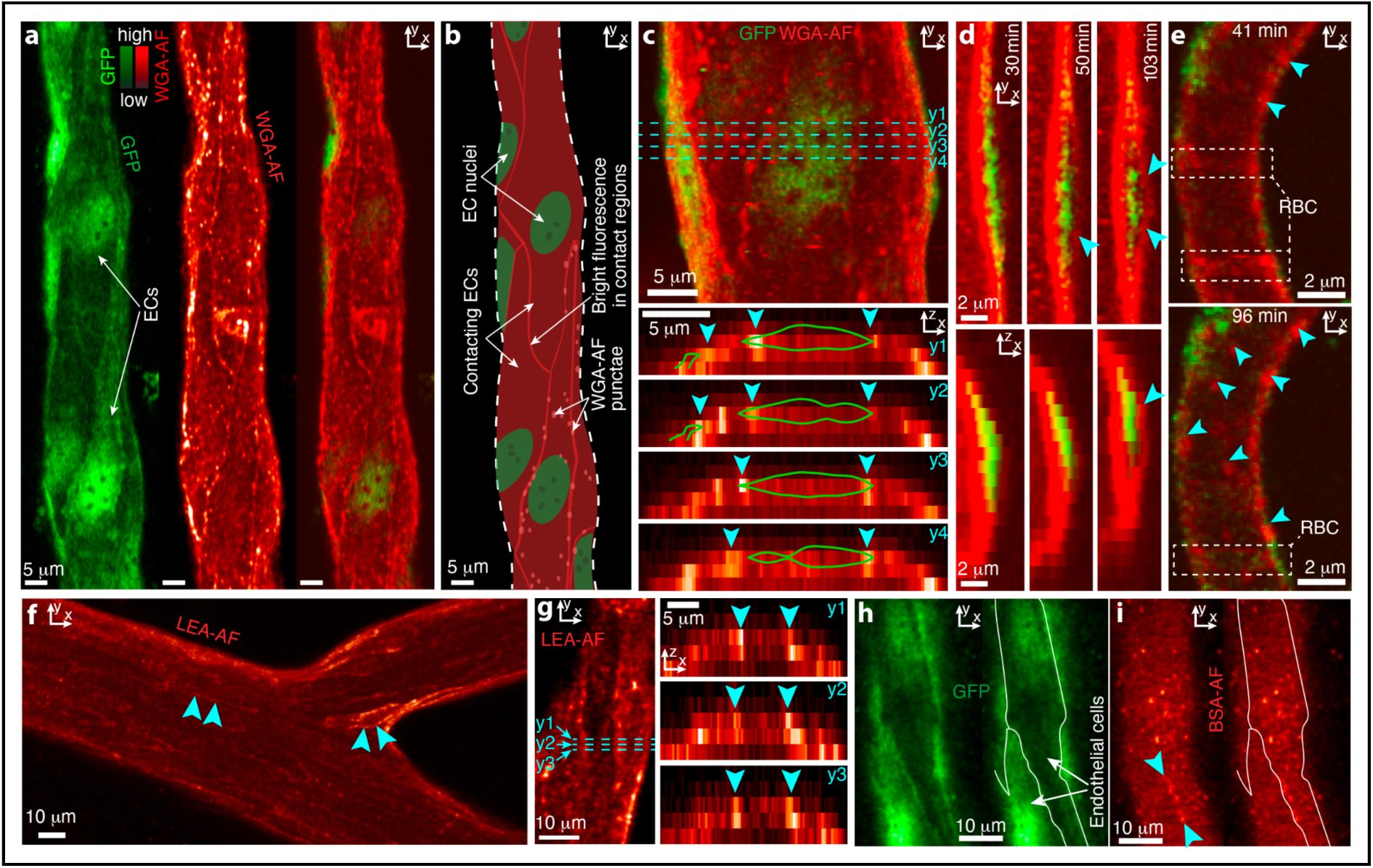
Micro-heterogeneity of glycocalyx on single ECs. **(a)** Maximum projection showing a pial arteriole (same vessel as in Fig.1c) with endothelial GFP (*left*), WGA-AF (*middle*), and their overlay (*right*). **(b)** A drawing of main features of images in (a). **(c)** *Top*: Magnified region from (a). *Bottom*: Cross-sections (*xz*) of the arteriole averaged over a 1-µm-wide y-region centered at dashed cyan lines in *top*. Arrowheads: local maxima of WGA-AF fluorescence intensity on the sides of the EC, visualized as isocontours of the half-maximal intensity of GFP fluorescence (green). **(d)** Arteriolar EC (vessel lumen on the left), shown in xy-(top) and xz-plane (bottom) 30, 50 and 103 min after WGA-AF injection. Arrowheads: Abluminal WGA-AF. **(e)** Maximum projections showing a capillary at 41 (top) and 96 (bottom) minutes after WGA-AF injection. Rectangles show WGA-AF-labeled red blood cells (RBC). Arrowheads: WGA-AF punctae. **(f)** Maximum projection showing a pial arteriole with LEA-AF. Arrowheads: WGA-AF-like labeling bands. **(g)** *Left*: Maximum projection of a pial arteriole’s top with LEA-AF-labeled glycocalyx. *Right*: same as in (c; bottom), for LEA-AF. **(h)** GFP fluorescence on the surface of a pial arteriole (left) and the same image with marked ECs perimeters (right). **(i)** BSA-AF fluorescence on the surface of arteriole from (g), showing luminal fluorescence, punctae, and bands of fluorescence near ECs perimeters (arrowheads).

At the surfaces of pial venules, we did not observe the same labeling pattern (Fig. 1h), which may be due to our limited ability to distinguish the weak venous glycocalyx fluorescence from the intraluminal fluorescence (Fig. 1i–j). In capillaries, the size of individual endothelial cells (ECs; ∼5–10 µm wide, Fig. 2a–c) is comparable to the capillary diameter. As a result, a single EC does not lie flat—as it does at the top of the cylindrical surface of a larger arteriole—but curves along the capillary wall. Therefore, resolving two contacting ECs in the same focal plane (2D) —which is optimal for resolving glycocalyx enrichment at contact sites—was impossible in capillaries. Alternatively, doing the same in 3D is further complicated by the capillary wall curvature and low spatial resolution along the z-axis (Extended Data Fig. 2i). Furthermore, glycocalyx fluorescence intensity in capillaries is confounded by fluorescently labeled RBCs, and we were unable to obtain RBC-free images due to the slow image acquisition inherent for 3D imaging with a scanning microscope.

Next, we detected accumulation of WGA-AF fluorescence on the abluminal side of the endothelium within ∼1 hour after WGA-AF injection (Fig. 2d). Furthermore, WGA-AF exhibited punctate labeling of arterioles (Fig. 2c), providing evidence of vesicular transport^39,40^. In venules and capillaries, low WGA-AF and GFP fluorescence intensities limited their 3D localization, which is necessary to monitor transendothelial WGA-AF transport in real time. Nevertheless, we noted punctate WGA-AF labeling in capillaries appearing at similar time post-injection (Fig.2e) as in arterioles, suggesting that vesicular transport also occurred in capillaries.

To rule out that the enrichment of WGA-AF fluorescence at the periphery of ECs and its punctate-like distribution were specific to WGA-AF, which binds a variety of residues with N-acetylglucosamine (GlcNAc) and sialic acid being the primary binding targets^26,34,41^, we next examined glycocalyx labeling with two additional probes. First, we tested Alexa Fluor 594–conjugated Lycopersicon esculentum agglutinin (LEA-AF), which preferentially binds polyLacNAc structures and similar to WGA also binds GlcNAc oligomers^34^, and as expected produced labeling patterns comparable to WGA-AF, including elevated fluorescence at EC peripheries and punctate foci (Fig. 2fg, Extended Data Fig. 3a). Second, we tested Alexa-Fluor 594-conjugated bovine serum albumin (BSA-AF), an analog of endogenous serum albumin, which does not interact selectively with glycans, but is known to interact with and stabilize the glycocalyx^42^. Similarly to WGA-AF and LEA-AF, BSA-AF formed punctae, reflecting vesicular transport^33,43–46^, and showed slightly elevated fluorescence at the peripheries of ECs (Fig. 2hi, Extended Data Fig. 3bc). In contrast, a Texas Red–conjugated 40-kDa dextran, a commonly used plasma tracer that does not bind to the glycocalyx^47^, produced neither a detectable glycocalyx labeling nor patterns of puncta uptake into endothelium (Extended Data Fig.1e). Thus, based on the used probes, we showed that the *in vivo* glycocalyx is heterogeneous at the leVel of single ECs and exhibits punctae, suggesting vesicular transport by ECs.

### Glycocalyx heterogeneity along the arteriovenous axis

Having established that glycocalyx fluorescent labeling is heterogeneous at the level of single ECs, we next quantified glycocalyx heterogeneity along the vascular tree. Although glycocalyx labeling was present in all types of the brain microvasculature, the intensity of WGA-AF fluorescence varied between the vessel segments (Fig. 1d–j). The strongest glycocalyx labeling was detected in arterioles, then capillaries, while venules displayed the lowest WGA-AF intensities. Despite earlier reports of absent venous labeling^16^, we reliably detected the venous glycocalyx by resolving the adjacent RBC-free plasma zone (I_plsm_ in Fig. 1e,i,j).

To quantify the arteriovenous glycocalyx heterogeneity, we introduced the *glycocalyx maps* (Fig. 3a-d, Extended Data Fig. 4), which quantified the spatial distribution of the relative glycocalyx intensities, 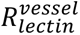 (*Methods: Glycocalyx maps*), of WGA-AF and LEA-AF in different vessel types along the arteriovenous axis (vessel segment classification follows Ref. ^43^). Capillaries were excluded because their glycocalyx fluorescence was contaminated by RBC-bound lectins (Fig. 2e, Fig. 7a). The glycocalyx maps in Fig. 3b,d demonstrated that the glycocalyx intensities of WGA-AF and LEA-AF (Fig. 3e,f) varied several-fold on the scales of (i) tens of micrometers, corresponding to intra-endothelial cell variations (Fig. 2a-c), and (ii) several micrometers, corresponding to the size of individual WGA-AF puncta (Fig. 2e).

**Figure 3.**
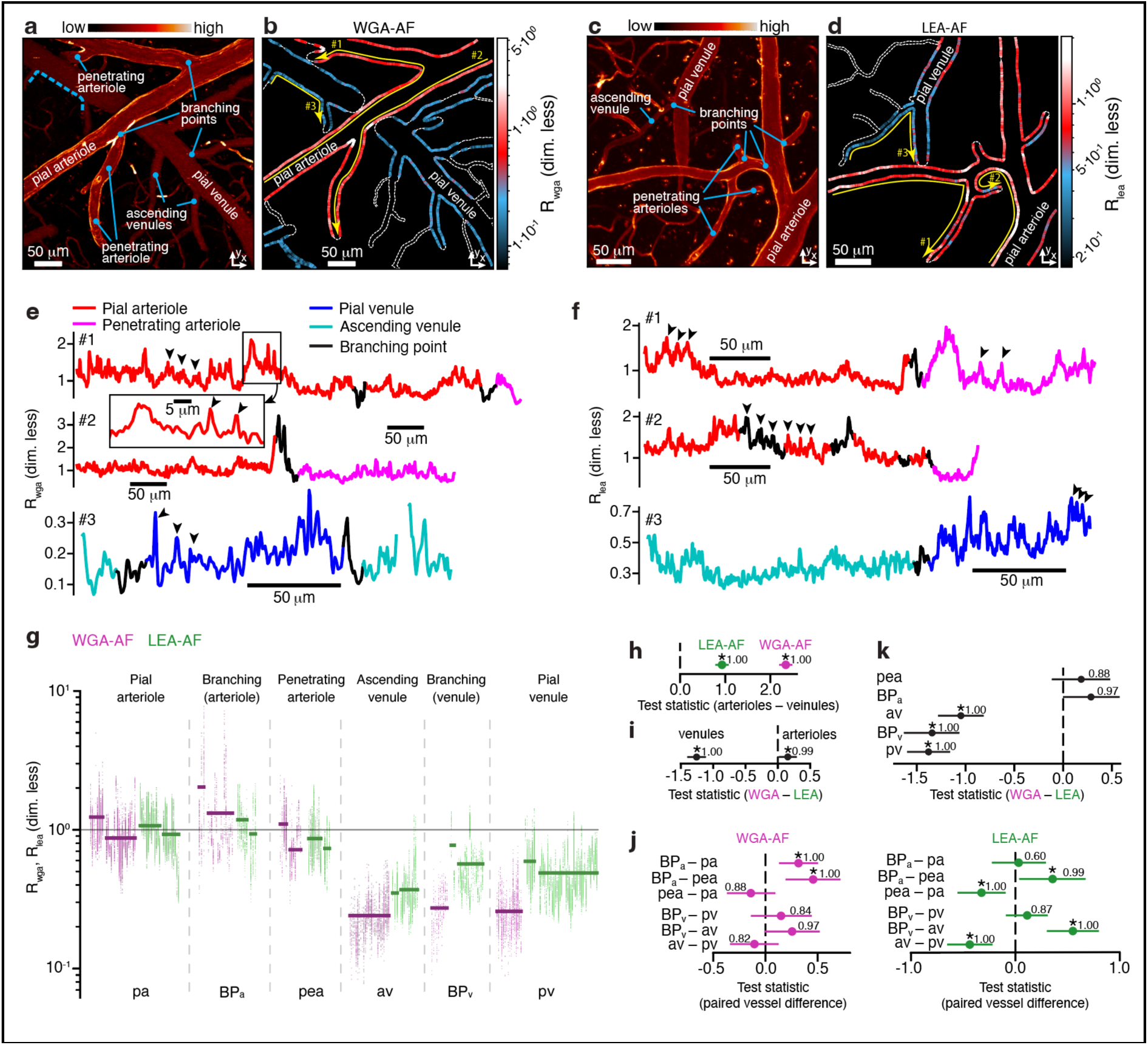
Glycocalyx heterogeneity along the arteriovenous axis. **(a)** Blood vessels labeled with WGA-AF (maximum projection; Fig.1c). **(b)** Map of the relative glycocalyx intensities for WGA-AF (R_wga_), estimated along vessel contours from (a), e.g., contour #3 corresponds to the dashed blue contour in (a). Dashed lines: vessels where glycocalyx could not be localized. **(c,d)** Same as in (a,b), but for LEA-AF (R_lea_) in a different mouse. **(e)** R_wga_ in arterioles (#1,#2) and venules (#3), along yellow contours with corresponding numbers in (b). *Inset*: a region of trace #1 (same *y*-scale) illustrating heterogeneous fluorescence with local maxima (arrowheads). **(f)** Same as in (e), but for R_lea_ in arterioles (#1,#2) and venules (#3). Arrowheads: local maxima of R_lea_. **(g)** Clouds of dots: R_wga_ (magenta) and R_lea_ (green) for different vessel types. One cloud contains R_wga_ obtained from multiple contours from the same vessel. Horizontal bars: Average R_wga_ and R_lea_ for a given vessel. **(h)** Mean of test statistic (TS), representing the difference in R_wga_ (magenta) and in R_lea_ (green) between arteriolar (pea, BPa) and venous segments (pv, av, BPv). **(i)** Means of TS, representing differences between R_wga_ and R_lea_ for arteriolar and venous segments. **(j)** Means of TS, representing differences in R_wga_ (left) and in R_lea_ (right) between six pairs of vessel types. For example, TS>0 for the *pa*–*pea* pair (left) indicates higher glycocalyx fluorescence in *pa* than in *pea*. **(k)** Same as in (i) but for five vessel segments. **(h-k)**: vessel-segment abbreviations are explained along the x-axis in (g); lines denote [2.5 %, 97.5 %] quartile ranges of TS; * denotes statistical difference. Data are from four mice: two labeled with WGA-AF and two labeled with LEA-AF. See Statistical models (Model A) in Methods for details.

Furthermore, for both WGA-AF and LEA-AF, venous segments exhibited lower glycocalyx intensities than arteriolar segments (Fig. 3e-h), which suggests either an overall denser (or thicker) glycocalyx in arterioles or a constant density (thickness) combined with a relative enrichment of WGA-AF- and LEA-AF-binding motifs in arterioles compared to venules. Notably, the arteriovenous difference was greater for WGA-AF than for LEA-AF (Fig. 3g,h). Glycocalyx intensities in penetrating arterioles were lower than in pial arterioles for LEA-AF but not for WGA-AF (Fig. 3g,i). Glycocalyx intensities in ascending venules were lower than in pial venules for LEA-AF but not for WGA-AF (Fig. 3j). Thus, *in vivo* glycocalyx mapping revealed higher overall lectin labeling in arterioles than in venules with lectin-specific labeling differences along the arteriovenous axis (Fig. 3h-k), highlighting vessel-type– dependent glycocalyx organization.

### Glycocalyx of arteriolar branching points shows enrichment of WGA-, but not LEA-binding motifs

In addition to varying lectin fluorescence along the arteriovenous axis, we observed elevated fluorescence intensities near arteriolar branching points, visible both on glycocalyx maps (Fig. 3e,f) and on three-dimensional reconstructions of pial arterioles (Fig. 4, Extended Data Fig. 5). Statistical analysis confirmed the increased glycocalyx intensities at arteriolar branching points when compared to adjacent pial arterioles for WGA-AF, but not for LEA (Fig. 3i,j). In comparison, we found no differences between glycocalyx intensities in venous branching points compared to pial venules neither for WGA-AF nor for LEA (Fig. 3i,j).

**Figure 4.**
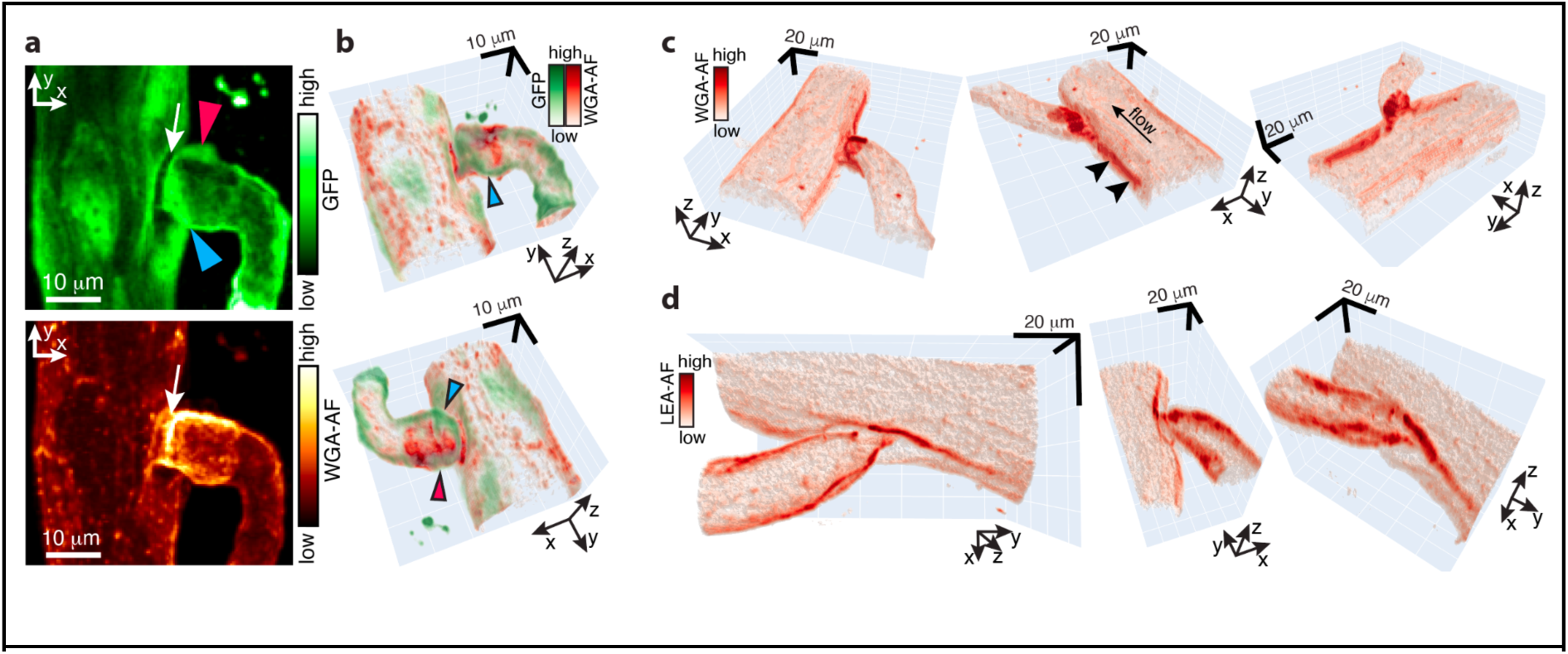
Arterial branching points are enriched in WGA-binding motifs. **(a)** Maximum projections of a volume stack, capturing a branching point of a pial arteriole (white arrow) with GFP fluorescence in the endothelium (*top*) and WGA-AF fluorescence in the glycocalyx (*bottom*). Two bright bulges of GFP fluorescence on two sides of the branching point (arrowheads) correspond to two ECs. **(b)** The same branching point shown in 3D. The blue and red arrowheads point to the same locations as in the *top* subpanel in (a). **(c)** Arterial branching point with WGA-AF labeled glycocalyx from another mouse, shown from three angles. *Middle:* High WGA-AF fluorescence (arrowheads) extends upstream the blood flow (arrow). *Right:* The side of the parent vessel opposite to the branching point shows no increase of the glycocalyx labeling. **(d)** Arterial branching point with LEA-AF labeled glycocalyx from another mouse, shown from three angles.

Three-dimensional imaging of WGA-AF-labeled glycocalyx in Tie2GFP mice showed that the elevated WGA-AF fluorescence near arteriolar branching points was adjacent to ECs nuclei at the base of daughter branches (Fig. 4a,b). Some of the branching points exhibited notably more extended and complex labeling patterns at the start of the daughter branches (Extended Data Fig. 5), while at some branching points WGA-AF fluorescence extended upstream in the direction opposite to the blood flow (Fig. 4c,d). Thus, our TPM *in vivo* imaging revealed an enrichment of WGA-labeled glycocalyx at arteriolar branching points suggesting localized structural specialization absent in venous branching points.

### Kinetics analysis suggests a novel transient interaction of WGA with the glycocalyx

Whilst dual-lectin glycocalyx maps revealed spatial heterogeneity of lectin-binding motifs along the brain vasculature, they did not specify the chemical nature of the motifs for a given lectin, which typically bind two or more different glycocalyx motifs^34^. Kinetic modelling determines reaction constants of glycocalyx-lectin interactions, enabling a more specific chemical profiling of glycans^26^. Therefore, we next performed kinetic analysis of fluorescence recovery after photobleaching (FRAP)^48–50^ of WGA-AF in pial arterioles (Fig.5), where the glycocalyx fluorescence intensity, I_gcx_ (Fig.1e, Fig.5a), was highest, but not in venules, where the fluorescence was too weak, and not in capillaries, where WGA-AF-labeled RBCs contaminated the glycocalyx fluorescence (Fig. 2e, Fig.7a). Defined that way I_gcx_ includes the fluorescence from both bound and free WGA-AF within the glycocalyx.

**Figure 5.**
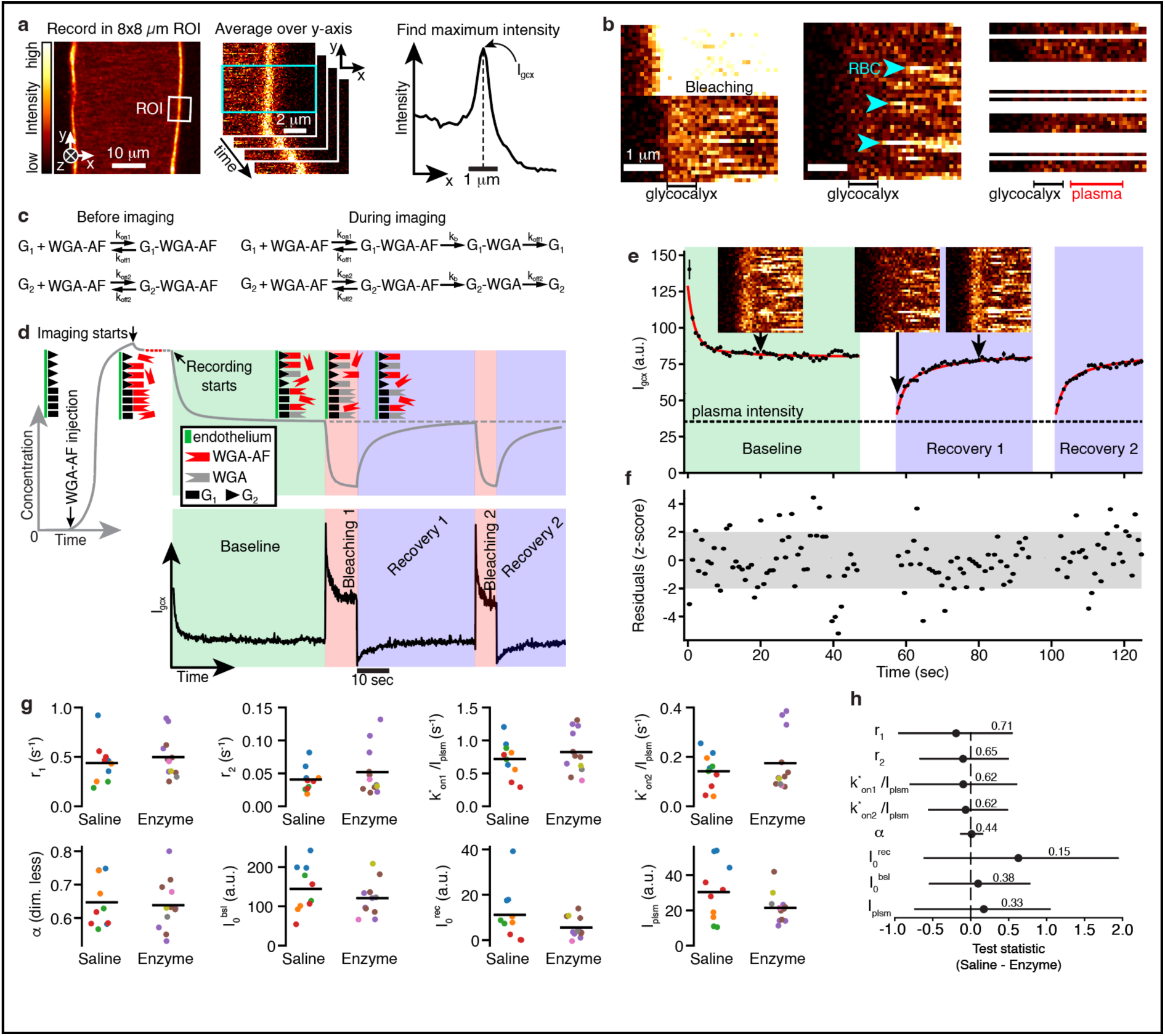
WGA-AF-glycocalyx interaction studied with fluorescence recovery after photobleaching (FRAP). **(a)** We estimated WGA-AF intensity in glycocalyx by recording images in a region of interest (ROI) (*left*), averaging images along y-axis (*middle*), and extracting the maximum value, I_gcx_, of the line-profile of WGA-AF fluorescence (*right*). **(b)** *Left*: Step-like increase in fluorescence intensity at the start of the bleaching phase, initiated by increasing laser intensity. *Middle*: Bright horizontal line-segments corresponding to WGA-AF-labeled red blood cells (RBCs). *Right*: Image from the middle without rows containing the RBCs. Plasma fluorescence is estimated by averaging pixels in RBC-free image lines over *plasma* region **(c)** Reaction models before and during imaging. G_1_ and G_2_ are two glycocalyx motifs binding WGA-AF. **(d)** Grey curve: Expected time-course of the concentration of bound and fluorescent WGA-AF in the glycocalyx following WGA-AF injection. Cartoons: distribution of fluorescent and bleached WGA-AF. Black curve: Measured I_gcx_(t). Note that I_gcx_ depends on the concentration of fluorescent WGA-AF and on laser intensity; Hence, at the onset of bleaching, I_gcx_ increases sharply, while concentration of fluorescent WGA-AF does not. **(e)** Dots with error bars: time-averaged I_gcx_(t) and the corresponding s.e.m.. Red curves: Fit of Eq.3 to the points with error bars(Online Methods). Dashed line: WGA-AF intensity in the plasma. The three images illustrate the reversible WGA-AF bleaching. **(f)** Standardized residuals. In ideal fit, they have a standard normal distribution with 5% of data outside the grey area. **(g)** Fitted parameters of Eq.3 in four saline- and five enzyme-treated mice. Dots of the same color are from the same mouse. **(h)** Means of test statistic (TS) representing the differences between saline (four mice) and enzyme groups (five mice) for all fitted parameters. TS>0 indicates saline higher than enzyme. The line denotes [2.5 %, 97.5 %] quartile range of TS; * denotes statistical difference. See Statistical models (Model C) in Methods for details.

At the beginning of a FRAP experiment (*Baseline*), I_gcx_(t) decreased to a steady-state value due to undesired but unavoidable photobleaching (black trace in Fig.5d). Next, increasing the laser intensity (*Bleaching*) caused a higher photobleaching rate (steeper decay of the black trace in *Bleaching* compared to *Baseline* in Fig.5d) and a step-like increase of fluorescence intensity (Fig.5b, left), as it is proportional to the squared laser intensity. Note that despite this artificial intensity increase, the expected number of fluorescent glycocalyx-bound WGA-AF was decreasing during the *Bleaching* (grey curve in Fig. 5d). Next, we decreased the laser intensity back to the initial value and recorded the recovery of I_gcx_ (*Recovery*, Fig. 5d). After that, we repeated the *Bleaching* and *Recovery* phases one more time (Fig. 5d).

We derived our model (Eq.3; Supplementary Note: Modeling B) assuming (i) independent, reversible binding of WGA-AF to two motifs, G_1_ and G_2_ (Fig. 5c,d), which have different binding/unbinding rate constants, motivated by an improved fit of the data with the double-motif model^26^ (Fig. 5e vs. Extended Data Fig. 7ab); (ii) irreversible photobleaching of glycocalyx-bound WGA-AF; (iii) unbinding of bleached complexes into free G_1_ and G_2_ motifs and free WGA, rapidly cleared by the blood flow (*Supplementary Note: Modeling B*).

Fitting Eq.3 to the data (Fig. 5ef) revealed two recovery rates r_1_=0.44 ± 0.14 and r_2_=0.04 ± 0.01 s^-1^ (Fig. 5g), which are sums of unbinding rate constants for each motif (𝑘_𝑜𝑓𝑓1_, 𝑘_𝑜𝑓𝑓2_) and the photobleaching rate constant, 𝑘_𝑏_ (Fig. 5c). Because 𝑟_1_ ≫ 𝑟_2_ > 𝑘_𝑏_ we could neglect 𝑘_𝑏_ in the expression 𝑟_1_ = 𝑘_𝑜𝑓𝑓1_ + 𝑘_𝑏_ and estimate the unbinding rate constant 𝑘_𝑜𝑓𝑓1_ ≈ 𝑟_1_ = 0.44 ± 0.14 s^-1^. The respective motif G_1_ accounted for 65±20% of all binding motifs of WGA-AF (Fig. 5g), and its binding pseudo-rate constant, 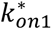, was higher than that of G_2_ motif, 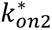 (Supplementary Table. 1, Fig. 5g). Similarly, Liu et al. identified two *in vitro* WGA-glycocalyx unbinding rate constants of 5.3×10^-4^ s^-1^ and 2.4×10^-2^ s^-1^, and attributed them to GlcNAc and N-Acetylneuraminic acid in the glycocalyx, respectively^26^. Our lowest fluorescence recovery rate, r_2_ = 0.04 ± 0.01 s^-1^ agrees with the highest of the two unbinding constants of Liu et al. We have not identified a recovery rate constant, corresponding to the unbinding constant of 5.3×10^-4^ s^-1^ found by Liu et al.^26^, as this would require a ∼1/(5.3×10^-4^ s^-1^) = 31-min-long recording, unfeasible *in vivo* because of the vascular out-of-focus motion.

The two motifs included in our model, G_1_ and G_2_, do not necessarily correspond to two distinct chemical motifs on the glycocalyx. Considering the complexity of the glycocalyx chemical composition that can affect kinetics of WGA-AF binding^41^ (e.g., sialic acid, number of GlcNAc’s, modifications of glycosaminoglycans), G_1_ and G_2_ likely correspond to two groups of glycocalyx motifs with similar kinetics of WGA-glycocalyx interaction within each group.

To test whether altering composition of the glycocalyx has an effect on kinetics of WGA-AF and glycocalyx interactions, we performed FRAP experiment after enzymatic shedding of glycocalyx by a mix of heparinase I and hyaluronidase. Hyaluronidase cleaved the hyaluronan, as shown by the increased plasma hyaluronan concentration^51,52^ (from 50 ± 5 ng/ml to 109 ± 21 ng/ml; Fig. 6a). Heparinase I cleaves heparan sulfate, which we confirmed with the reduced fluorescence intensity of 10E4 antibody-immunostained heparan sulfate in the brain vessels (Fig. 6b,c; Extended Data Fig. 6). Thus, the enzymes cleaved heparan sulfate and hyaluronan—two major WGA-binding GAGs of the glycocalyx—but there are likely other types of GAGs that remained after the shedding, including keratan sulfates, chondroitin sulfates, dermatan sulfates, and glycosphingolipids with abundant sialic acid.

**Figure 6.**
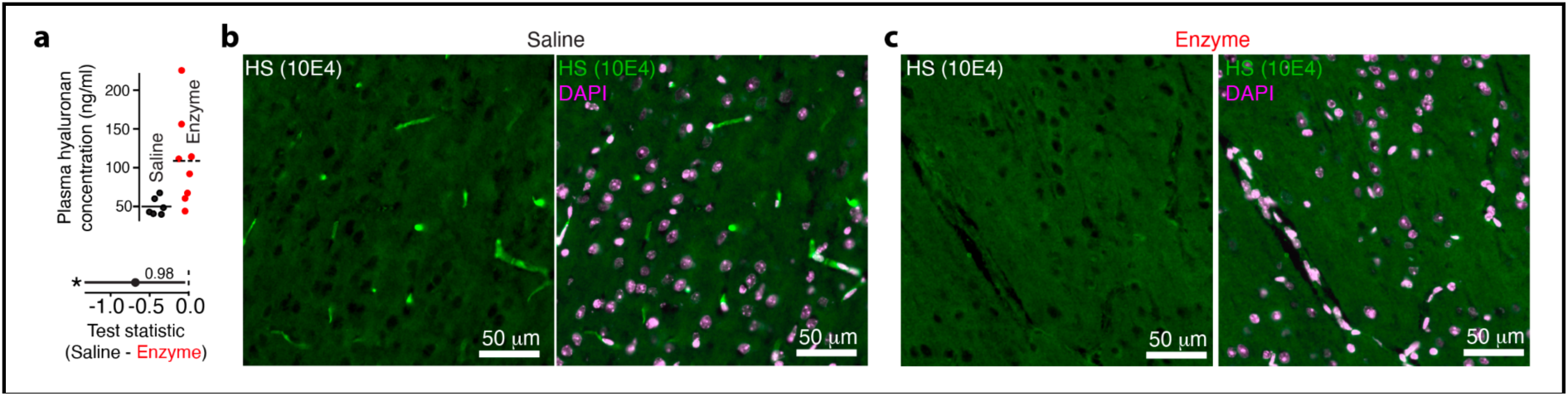
Intravenously injected Heparinase I and Hyaluronidase shed glycocalyx globally and locally, in the brain. **(a)** *Top*: Plasma concentration of hyaluronan fragments in saline- and enzyme-treated mice. *Bottom*: Mean of test statistic (TS) representing the difference between plasma hyaluronan concentration between three control (saline) mice and four enzyme-treated mice. For both groups, there are two repeated measurements per mouse. The line denotes [2.5 %, 97.5 %] quartile range of TS; * denotes statistical difference. See Statistical models (Model B) in Methods for details. **(b)** Immunohistochemical staining of heparan sulfate (10E4 antibody, green) in cerebral cortex of saline-treated mouse. *Left*: 10E4 staining. *Right*: 10E4 (green) and DAPI (magenta). **(c)** Same as (b), for enzyme-treated mouse.

Enzymatic degradation of heparan sulfate and hyaluronan had no effect on the fitted parameters (Fig. 5g,h). The unchanged 𝑟_1_ and 𝑟_2_ – hence unchanged 𝑘_𝑜𝑓𝑓1_ and 𝑘_𝑜𝑓𝑓2_ – were expected as the shedding of the glycocalyx motifs should not affect the kinetics properties of the remaining motifs. The unchanged relative proportion, 𝛼, between binding motifs G_1_ and G_2_, may indicate comparable shedding for both. Finally, 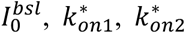 did not change during enzymatic treatment, despite that these parameters depend on the concentration of glycocalyx-binding motifs. This lack of change can be explained by a broad specificity of WGA-AF^34^, i.e., not limited to binding to the shedded heparan sulfate and hyaluronan. In summary, FRAP analysis uncovered for the first time two kinetically distinct interactions of WGA-AF with the glycocalyx *in vivo*, introducing a new methodological framework to interrogate the kinetics of molecular interactions with the glycocalyx in the living brain.

### The enzymatic shedding of heparan sulfate and hyaluronan thinned the glycocalyx by 153 nm but did not change its fluorescence intensity

Glycocalyx is commonly characterized *in vivo* by labeling it with a fluorescent lectin and estimating either its labeling intensity or its thickness. However, the lack of a standard method to estimate either of the two quantities has led to conflicting results across studies^16–18^. To resolve this discrepancy, we next asked: what is the most optimal way to quantify glycocalyx *in vivo*?

We first estimated glycocalyx intensity in arterioles, veins and in the capillaries using the same method as in the FRAP analysis (Fig. 5a and inset in Fig. 7b). Capillaries showed glycocalyx fluorescence intensity similar to that in venules and lower than in pial arterioles and penetrating arterioles (Fig. 7b). Similarly to the results of FRAP analysis, *in vivo* glycocalyx fluorescence intensities of WGA-AF remained unchanged across vessel types after the enzymatic treatment (Fig. 7b,c). We hypothesized that the maximal fluorescence intensity in the glycocalyx is a sub-optimal estimate as it relies on a single value—the maximum of the fluorescence intensity profile (Fig. 7b, inset)—rather than optimally using the entire profile.

To achieve that, we have developed an approach to estimate both the glycocalyx thickness and its fluorescence intensity by fitting a model (Eq. 4; Supplementary Note: Modeling C) to entire WGA-AF fluorescence line profiles (Fig. 7d,e). This approach is equivalent to super-localization microscopy^53^, where we localized a complex structure, consisting of several fluorescent compartments, including the lumen, glycocalyx, and extravascular space (Fig. 7e,f). We estimated glycocalyx thickness of 775 ± 17 nm (mean ± s.e.m.; Fig. 7g,h; Supplementary Table 2), consistent with 726 ± 148-nm-thick (mean ± s.e.m.) glycocalyx in brain pial arterioles estimated with electron microscopy^19^. Similarly, we resolved the endothelial layer as 123 ± 8 nm thick, six times thinner than the glycocalyx, also in agreement with electron microscopy^43,54^. Our model considers the accumulation of the dye on the abluminal surface of the ECs (Fig. 2d), allowing us to isolate only the fluorescence from the luminal surface (Fig. 7d). In contrast, estimating glycocalyx thickness with full width at half maximum (FWHM)^37^ or similar metrics^16,18^, applied to the entire WGA-AF fluorescence distribution – which includes unwanted contributions from plasma and the abluminal side of ECs – can lead to substantial overestimation, by as much as twofold (Fig. 7e, inset). Moreover, rapid image acquisition was essential to minimize motion blur and exclude artificial fluorescence from RBCs (Supplementary Table 3).

In agreement with the results in Fig. 7b,c, glycocalyx intensity remained unchanged after enzymatic shedding of heparan sulfate and hyaluronan (Fig. 7g,i; Supplementary Table 2), which may indicate that most of the WGA-AF is bound to e.g. sialic acid and other non-shed glycocalyx components. In addition, although the same amount of WGA-AF was injected into each mouse, plasma concentrations varied markedly across mice (I_plsm_ in Fig. 7g), which is a systematic, not stochastic, variability since the error bars on I_plsm_ are much smaller than the differences between mice (Extended Data Fig. 10). These systematic differences – arising from factors such as differences in body weight, blood volume, plasma clearance – propagated to the glycocalyx fluorescence intensity, producing noisy estimates.

In contrast, glycocalyx thickness, derived from the width of the fluorescence profile and not from its peak value, is not expected to vary with the plasma WGA-AF concentration. Therefore, a more precisely estimated glycocalyx thickness allowed us to detect that shedding of heparan sulfate and hyaluronan reduced the glycocalyx thickness by 153 ± 38 nm (20% decrease; Fig. 7g,h) – 2.3-fold smaller than the diffraction-limited resolution of our TPM (351 ± 4 nm, Extended Data Fig. 2e). The sub-diffractional resolution was achieved through optimal analysis of fluorescence line-profiles. Thus, the glycocalyx thickness, combined with super-localization analysis, becomes a metric-of-choice for studying enzymatic shedding of the glycocalyx and other processes leading to nanoscopic changes in the glycocalyx structure.

**Figure 7.**
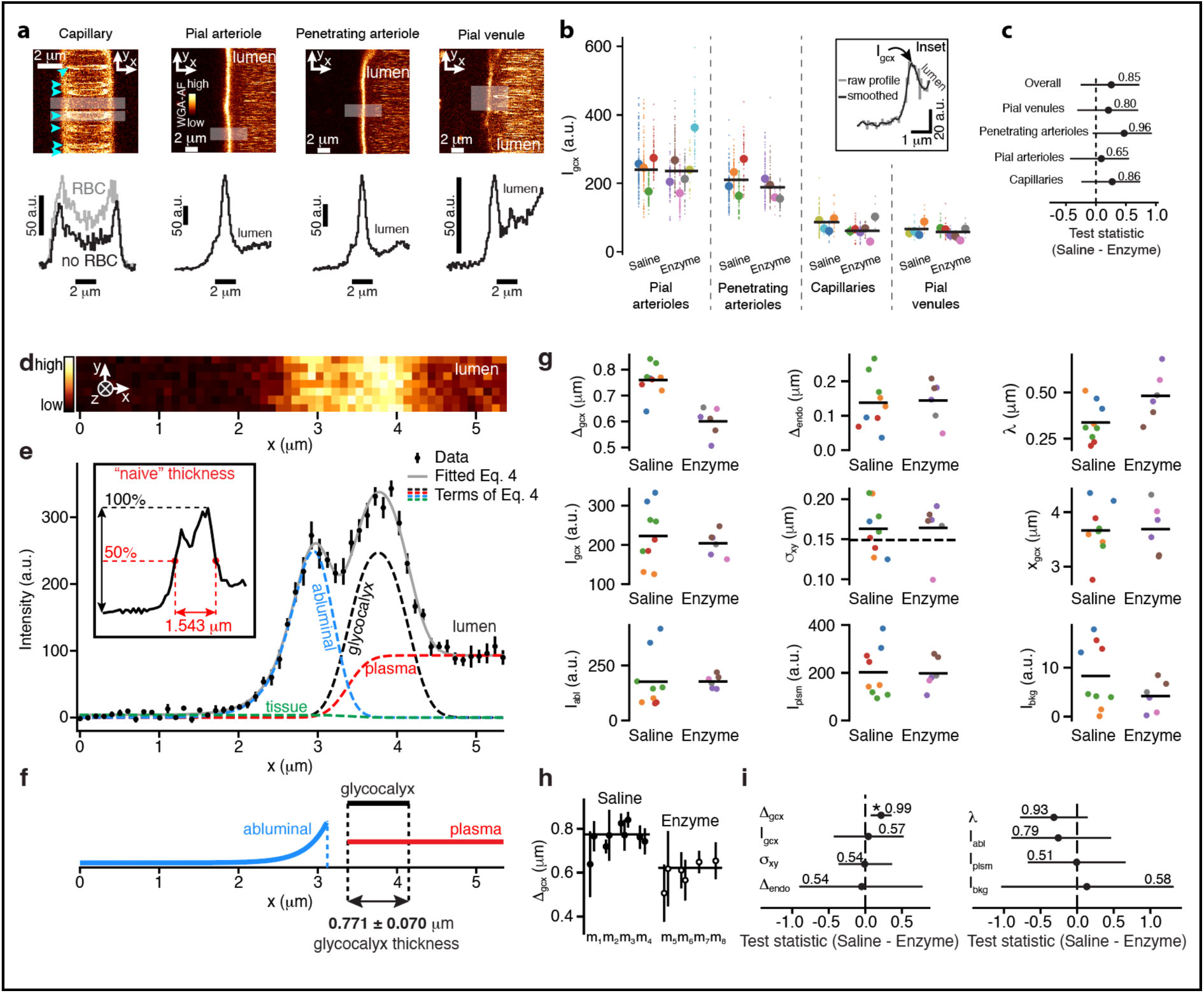
*In vivo* estimation of WGA-AF-labeled glycocalyx thickness and fluorescence intensity. **(a)** *Top*: WGA-AF fluorescence in four Vessel types. Cyan arrowheads: WGA-AF-labeled red blood cells. *Bottom*: fluorescence intensity profiles (two for capillary) estimated by averaging fluorescence along y-axis in white rectangles in top panels. **(b)** Small dots: Glycocalyx fluorescence intensity, I_gcx_, from individual profiles. Large dots: mean I_gcx_ for each mouse. Lines: means for saline- and enzyme-treated mice groups. *Inset:* Estimation of I_gcx_. **(c)** Means of test statistic (TS) representing the differences in I_gcx_ between saline (four mice) and enzyme groups (six mice). The line denotes [2.5 %, 97.5 %] quartile range of TS; * denotes statistical difference. See Statistical models (Model D) in Methods for details **(d)** WGA-AF-labeled glycocalyx (*x*∼4µm) of a pial arteriole. **(e)** Dots with error bars: mean ± s.e.m. of fluorescence intensity profile, calculated by averaging image in (a) along *y-*axis. Gray curve: Fit of Eq.4. Colored curves: Terms of fitted Eq.4; They sum to gray curve. Inset: Naive estimation of glycocalyx thickness. **(f)** Model of WGA-AF distribution in abluminal, glycocalyx, and plasma compartments (Eq.4; Supplementary Note Modeling C) and estimation of glycocalyx thickness based on fitting Eq.4 in (b). **(g)** Dots: Fitted parameters of Eq. 4: 𝛥_*g*𝑐𝑥_, glycocalyx thickness; 𝛥*_endo_*), endothelium thickness; λ, length constant of abluminal fluorescence decay; *I*_*g*𝑐𝑥_, glycocalyx fluorescence intensity; σ*_xy_*, standard deviation of the PSF in xy plane; *x*_*g*𝑐𝑥_, glycocalyx location; *I_abl_*, abluminal fluorescence intensity; *I_plsm_*, plasma fluorescence intensity; *I_bkg_*, background fluorescence intensity. Horizontal lines: within-group means. Dashed line: σ_xy_ measured from nanobeads (Extended Data Fig.2e). **(h)** Same as top-left (g), with error bars (s.e.m.) from fitting; m1, m2,..: mice. Horizontal lines: weighted averages, 775±17 nm and 622±34 nm, before and after enzymatic shedding, respectively. **(i)** Means of test statistic (TS) representing the differences between saline (four mice) and enzyme groups (four mice) for all fitted parameters of Eq.4. The line denotes [2.5 %, 97.5 %] quartile range of TS; * denotes statistical difference. See Statistical models (Model E) in Methods for details.

## Discussion

The critical physiological role of the endothelial glycocalyx is widely recognized^11,19,30,38^, yet studying it — especially in the dynamic environment of the living brain — remains challenging due to nanoscopic size of the glycocalyx, combined with its motion with the pulsating vessel walls, and due to complex glycan chemistry, resulting in a partial (e.g. if a single fluorescent lectin is used) and heterogeneous glycocalyx labeling. Here, we addressed these challenges by combining *in vivo* TPM with dual-lectin labeling, FRAP analysis, enzymatic degradation, and super-localization-based quantification of glycocalyx thickness with nanoscale resolution. Our work provides a significant step towards a multi-scale characterization of glycocalyx in the living brain (Fig. 8).

**Figure 8.**
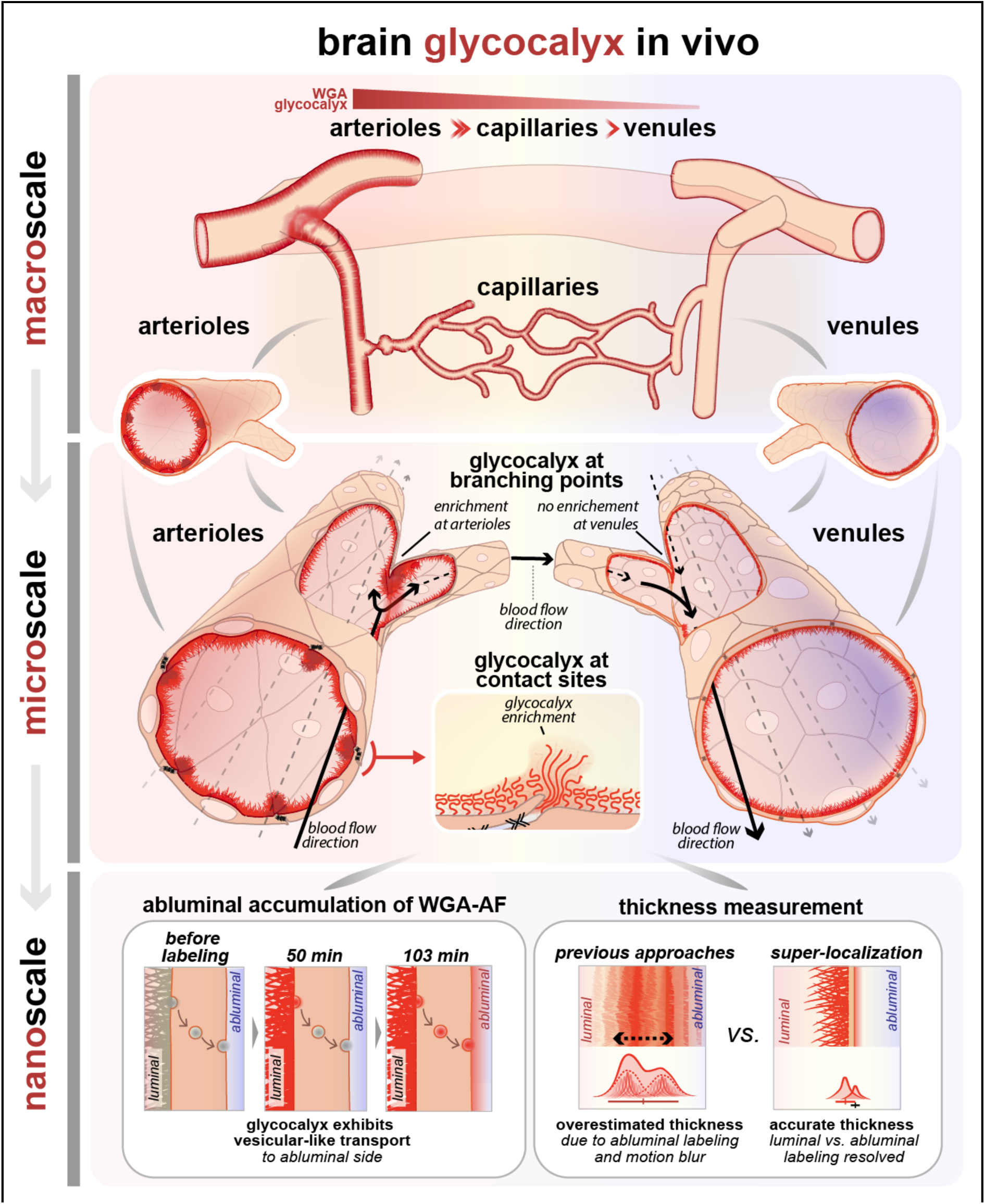
In vivo glycocalyx heterogeneity and topology on the macro-, micro-, and nano-scale. On the macro-scale, arterioles showed the highest glycocalyx labeling intensity (WGA-AF), followed by capillaries and venules. On the microscale, arterioles exhibit glycocalyx enrichment at ECs contact sites and in the branching points, where the blood flow shear stress is the highest. On the nanoscale, *glycocalyx* undergoes transcytosis-like transport to the abluminal side. With super-localization, we estimated 775 ± 17 nm-thick in pial arterioles, consistent with TEM ultrastructure analyses.

At the subcellular level, we found that glycocalyx was enriched at ECs contact sites, observed here as bright bands of both WGA-AF and LEA-AF fluorescence along the perimeters of individual ECs. This local enrichment of fluorescence can stem from an increased local thickness of the glycocalyx, from increased local density, or from both combined (Fig. 8, inset). The latter can occur, e.g., if two ECs with uniformly distributed glycocalyx meet at their contact sites. There, the overlapping glycocalyxes from the ECs can force the glycocalyx glycans and proteins to pack more densely and stretch (Fig. 8, inset), as polymer brushes do^55,56^. Both increased glycocalyx density and thickness will increase WGA-AF fluorescence due to an increased number of bound WGA-AF molecules within the volume from which we record the fluorescence (Fig. 8, inset). In agreement with our TPM data, lanthanum-labeled glycocalyx appeared more dense near the paracellular cleft in the brain vessels (Fig. 1a in Ref.^19^), and heparan sulfate concentration was elevated in peripheral regions of ECs in the lung^22^, rat adipose tissue, and umbilical vein exposed to shear stress^7,23^. The enriched glycocalyx between neighboring ECs may enhance the paracellular barrier by limiting the influx of molecules to the tight junctions and possibly shielding them from the shear stress.

Notably, we observed frequent WGA-AF and LEA-AF puncta coinciding with WGA-AF accumulation on the abluminal side of ECs within one hour post-injection, similar to BSA shuttled via adsorptive-mediated transcytosis^43^. Since both lectins are unlikely to cross the BBB via the paracellular route due to their sizes (36 kDa for WGA, 71 kDa for LEA)^57^, the punctate labeling suggest vesicular transport of the lectins, which is supported by earlier observations of WGA transcytosis^39,40,58^. Electron microscopy also showed vesicular uptake of the glycocalyx into peripheral endothelia^19^. Glycocalyx endocytosis is a key mechanism of glycocalyx turnover, and it may have occurred along with the lectin uptake observed here — either passively or via glycocalyx-specific endocytosis receptors^38^.

Lectins have been widely used in decoding the chemical composition of glycocalyx, yet no such data is available for the brain glycocalyx *in vivo*^38^. At the level of single blood vessels, our glycocalyx maps revealed a zonation of WGA-AF and LEA-AF labeling. Fluorescence intensity of both WGA-AF and LEA-AF in the glycocalyx decreased from arterioles to venules, consistent with reports of reduced glycocalyx density in the venular end of the brain microcirculation^16,59^. The decrease was more pronounced for WGA-AF than for LEA-AF (Fig. 3h), suggesting a compositional difference between arteriolar and venous glycocalyces. That is, the relative density of oligomeric GlcNAc, terminal GlcNAc, and sialic acid—major binding motifs of WGA^26,34,41^—decreases from arterioles to venules more than that of oligomeric GlcNAc, polyLacNAc, or LacdiNAc—major binding motifs of LEA^34,41^. Note that the glycocalyx exhibits a multilayered structure, with the innermost layer comprising the bases of glycoproteins, proteoglycans, and glycosphingolipids anchored to the EC’s plasma membrane, and the outermost layer forming a meshwork of glycans and entrapped plasma components^21^. Therefore, when discussing compositional differences here, we refer specifically to the composition of the glycocalyx region (“layer”) accessible to WGA-AF and LEA-AF. The smaller lectin, WGA-AF (36 kDa), can potentially penetrate deeper into the glycocalyx compared to the larger LEA-AF (71 kDa). Considering this, the observed differences in the glycocalyx maps obtained with WGA-AF and LEA-AF may reflect a functional specialization of microvascular segments—for example, supporting blood flow regulation in arterioles and capillaries, while facilitating immune cell–endothelial interactions in venules along the arteriovenous axis.

We further showed that arteriolar branching points — i.e., areas exposed to elevated shear stress^60^ — have the highest glycocalyx fluorescence intensity of WGA-AF, but not LEA-AF. Since shear stress stimulates glycocalyx synthesis^61^, particularly via incorporation of hyaluronan^62^, and since the glycocalyx facilitates efficient RBC flow^63^, the glycocalyx enrichment at arteriolar branches may be an adaptation supporting optimal perfusion to downstream capillaries. Our data suggest that WGA-binding motifs in glycocalyx may play a more prominent role in such adaptation than LEA-binding motifs.

*In vivo* kinetic analysis of WGA-AF interaction with the glycocalyx identified an unbinding rate constant of 0.44 ± 0.14 s^-1^, which has not yet been identified neither *in vivo* nor in *in vitro* experiments^26,64–66^. We believe, however, that the small but noticeable deviations between the data and the fitted theory in Fig.1e and Fig.2d (e.g. at ∼200 sec) in Liu et al.^26^ may indicate an unbinding constant similar to one reported here. The observed interaction with a high unbinding constant may describe the formation of an unstable and quickly dissociating complex of WGA-AF with the glycocalyx (Fig.4A in Ref.^67^). Identification of the chemical nature of this interaction is important for understanding lectin-glycocalyx interactions and requires further studies. Our FRAP analysis represents, to the best of our knowledge, the first *in vivo* application of this technique to individual brain microvessels to probe chemical interactions between circulating molecules and the endothelial glycocalyx.

Finally, with *in vivo* super-localization of the glycocalyx, we addressed a long-standing discrepancy between *in vivo* and *ex vivo* EM measurements of the glycocalyx thickness. Until now, *in vivo* glycocalyx thickness measurements^18,37^ were on average markedly greater than those obtained by EM^11,19^, which we explain by a number of confounding factors, of *in vivo* microscopy, that inflate glycocalyx thickness (Supporting Table 3)^68^. Approaches using two fluorescent labels, e.g., one in glycocalyx, one in endothelium^16,17,29^, are less likely to overestimate glycocalyx thickness, but have lower interpretability due to estimation of the combined thickness of the ECs with that of glycocalyx. Here, we developed a novel, single-lectin-based approach capable of detecting enzyme-induced thinning of the glycocalyx by 153±38 nm. Our baseline thickness was severalfold lower than in previous *in vivo* studies^16,18^, and agrees with recent EM data^19^. Therefore, our novel approach bridges the gap between *in vivo* and EM measurements of glycocalyx, and we believe that it establishes a new standard for *in vivo* glycocalyx quantification, enabling a more complete understanding of glycocalyx structure and function.

In summary, the brain endothelial glycocalyx has hitherto eluded precise characterization *in vivo*. We introduce a TPM strategy integrating dual lectin labeling, FRAP kinetics, enzymatic perturbation, and quantitative modeling to achieve the first high-resolution and multi-scale mapping of glycocalyx structure and dynamics in the living brain. We uncover a striking glycocalyx heterogeneity, with enrichment near endothelial junctions and at arteriolar branching points, and a compositional shift along the arteriovenous axis. Our super-localization-based glycocalyx thickness measurement reconciled conflicting optical and EM thickness estimates. These insights establish a novel paradigm for precise quantitative glycocalyx organization, dynamics, and resilience in the living brain.

## Acknowledgements

We would like to acknowledge Micael Lønstrup for his assistance with animal surgery and Birgit Poulsen for technical support with immunostaining. This study was supported by Lundbeck Foundation (#R273-2017-1791) and (#R223-2016-563); the Independent Research Fund Denmark (#1030-00374B); Det Frie Forskningsråd (#0602-01965B) and (#2066-00043B); Research grants from the Danish Cardiovascular Academy, which is funded by the Novo Nordisk Foundation, grant number NNF20SA0067242; Novo Nordisk Foundation (#NNF19OC0058796); and Læge Sophus Carl Emil Friis og hustru Olga Doris Friis’ Legat; and the Danish National Research Foundation (DNRF196).

## Methods

### Mouse models

We used 15 C57CL/6J, and one Tg(TIE2GFP)287Sato/J transgenic reporter mice (Tie2-GFP; cat #003658, Jackson Laboratory) expressing GFP in endothelium^69^. All mice weighed between 22–28 g and were 14–20 weeks old and were housed in ventilated cages at room temperature, 12 h light/12 h dark light cycle, and with *ad libitum* access to food and water. All procedures involving animals were approved by The Danish National Committee on Health Research Ethics in accordance with the European Council’s Convention for the Protection of vertebrate Animals Used for Experimental and Other Scientific Purposes. All procedures complied with the ARRIVE guidelines.

### Microsurgery

Mice underwent standard surgical procedures^32^. In brief, anesthesia was induced by intraperitoneal (i.p.) injection of xylazine (10 mg/kg animal) and ketamine (60 mg/kg animal) and sustained by ketamine (30 mg/kg animal i.p., every 20-25 min). The trachea was intubated for mechanical ventilation (180-220 μl volume; 190-240 strokes/min; Minivent Type 845 ventilator, Harvard Apparatus). The left femoral vein was cannulated to allow infusion of anesthetics during imaging. The femoral artery was cannulated to monitor mean arterial blood pressure (MABP; Pressure Monitor BP-1, World Precision Instruments), to measure arterial pCO_2_ and pO_2_, and to inject fluorescent probes (Fig.1a). The temperature of the animals was maintained at 37°C via a heating pad with feedback from a rectal thermal probe coupled to a homeothermic controller unit (EZ-TC-1000 Mouse Proportional). End-tidal CO_2_ (ETCO_2_) was monitored in real-time (Capnograph Type 340, Harvard Apparatus), and the respiration volume and stroke frequency were adjusted when necessary. Adjusting the ventilation parameters according to the mouse weight and blood gas, we ensured the normal physiological state of the mice: MABP: 60 – 80 mmHg, ETCO_2_: 2.2 – 2.8% (Ref. ^44^).

To prepare cranial windows, the scalp was removed, and the periosteum of the exposed skull was gently peeled off the bone. Next, the skull was dried and glued to a metal plate stabilized for imaging. The application of cooled saline to the bone prevented overheating during the drilling of the bone over the somatosensory (barrel) cortex. The bone flap was lifted, and the dura was removed. These procedures were performed under artificial cerebrospinal fluid (aCSF; in mM: NaCl 120, KCl 2.8, Na_2_HPO_4_ 1, MgCl_2_ 0.876, NaHCO_3_ 22, CaCl_2_ 1.45, glucose 2.55, at 37°C, aerated with 95% air/5% CO_2_ to pH 7.4). A drop of low-melting agarose (0.75% agarose, type III-A, Sigma-Aldrich) was applied to cover the brain surface and was covered with a glass coverslip (Menzel-Gläser, 24x66mm, #1.5) (Fig.1b). After surgery, the anesthesia was changed to alpha-chloralose (Sigma-Aldrich) infused into the venous catheter (0.5 g/mL, 0.01 mL/h/10 g).

### Fluorescent labeling of the glycocalyx

The glycocalyx was labeled by an intra-arterial (i.a.) injection of Alexa Fluor 594-conjugated wheat germ agglutinin (WGA-AF, Invitrogen; 150 µl of 1 mg/ml in saline injected within a few seconds) or by Alexa Fluor 594-conjugated tomato lectin (LEA-AF, Invitrogen; 300 µl of 0.5 mg/ml in saline injected within 5 minutes)^18,29,37^. The dose of WGA-AF was selected as low as possible, such that it does not change ETCO_2_ and MABP, yet sufficiently large to make glycocalyx bright on recorded images.

### Enzymatic treatment of the glycocalyx

Mice received either 150 µl saline (saline-treated mice) or hyaluronidase Type IV-S from bovine testes (3.8 mg; 750–3000 units/mg; Sigma Aldrich) and Heparinase I from *Flavobacterium heparinum* (9.4 IU; Sigma Aldrich) in 150 µl saline (enzyme-treated mice) by infusion into the carotid artery at the rate of 30 µl/min. Forty-five minutes after the infusion, the mouse was either perfusion-fixated for immunohistochemistry or injected with WGA-AF for two-photon microscopy.

### Immunohistochemistry

Two C57BL/6J mice were anesthetized and operated as described above, but without craniotomy and venous catheterization. Saline (one mouse) and the enzymes (one mouse) were infused as described above. Forty-five minutes after the injection, the mice were transcardially perfused with ice-cold PBS. The brains were extracted and post-fixed in freshly mixed 3/4 of 99.8% ethanol and 1/4 of glacial acetic acid. After five days, the brains were cryoprotected in 2M sucrose in PBS and frozen-cut into 4–7-µm coronal sections.

Brain sections were incubated with primary antibody diluted in 2.5% BSA in PBS (anti-heparan sulfate, 1:100, Amsbio EU, clone 10E4) overnight at 4°C, washed in 1× PBS, and incubated with secondary antibody, Alexa Fluor 488-conjugated goat anti-mouse (1:500, Molecular Probes, Thermo Scientific, EU). After washing, the sections were mounted using ProLong Gold Antifade Reagent with DAPI (Molecular Probes, Thermo Scientific, EU).

For imaging of the immunostained brain sections, we used an Axio Observer Z1 Inverted Fluorescence Microscope (Carl Zeiss Microscopy GMBH) equipped with a 20× (NA=0.8) Plan-Apochromat objective. The Alexa Fluor 488 and DAPI were excited using 475 nm and 386 nm LED-Modules, respectively. The fluorescence light was registered by an Axiocam 705 camera (Carl Zeiss Microscopy GMBH) after 500-550 nm (Alexa Fluor 488) and 430-470 nm (DAPI) band-pass filters. From the brain sections, we recorded tiled volume (xyz)-stacks (dimension of scanned brain area ∼ 9 mm × 6 mm × 24 µm; voxel size 0.345 µm × 0.345 µm × 3.000 µm) that were automatically stitched in ZEN software (Carl Zeiss Microscopy GMBH). From the stitched images, we selected regions in the cerebral cortex, hippocampus, thalamus, and striatum.

### Quantification of hyaluronan in blood plasma

Venous blood was collected from three saline-treated and enzyme-treated mice after the TPM imaging. The blood was drawn from the retro-orbital sinus via a 20-μl glass capillary flushed with EDTA before use. The blood samples were collected into K2-EDTA tubes and centrifuged at 4°C, 2000×g, for 10 minutes. The plasma was aliquoted (two aliquots per mouse), snap-frozen, and stored at -80°C. On day 1 Maxisorp Elisa plates (Thermo Scientific Nunc Cat# 476635) were coated overnight with 50 μl of human recombinant aggrecan G1-IGD-G2 domains (RnD cat# 1220-PG) at a concentration of 1.5 µg/ml in PBS. The next day, plates were washed three times with PBST (0.01%) using a Tecan hydroflex washer. Every subsequent wash was performed this way. Each well was then blocked with 200 μl PBST (5%) and incubated at room temperature for 1 hour to minimize nonspecific binding. After blocking, the plate was washed again, and 50 μl of each appropriately diluted standard, sample, and control were added to their respective wells. Each sample was assayed in duplicate. The plate was covered and incubated at room temperature for 2 hours and then washed to remove unbound substances. 50 μl of biotinylated human aggrecan at a concentration of 0.75 µg/ml was added to each well. The plate was then incubated at room temperature for 2 hrs. After the incubation, the plate was washed again to remove unbound biotinylated aggrecan and 50 μl of Streptavidin-HRP (Thermo Scientific Cat#N100) was added to each well, and the plate was incubated for 30 minutes at room temperature. A final wash was performed, and 100 μl of TMB ultra substrate (Thermo Scientific cat# 34029) was added to each well. After 15 minutes, the reaction was stopped with 50 μl 0.25M sulfuric acid, and the absorbance was read on a Thermo multiscan Fc. Concentrations were calculated using Graphad Prism by a 4-parameter nonlinear fit of the standard curve and interpolating the individual values from the samples. Plasma hyaluronan was measured in 3 saline- and 4 enzyme-treated mice, two measurements per mouse.

### Two-photon microscopy (TPM)

We used a Fluoview FVMPE-RS microscope (Olympus) equipped with a 25× (NA=1.05) water immersion objective. The two-photon excitation was delivered by a tunable femtosecond laser (Mai-Tai DeepSee) at 830 nm. The fluorescence light was captured by two high-sensitivity GaAsP detectors with cut-off frequencies 601-657 nm (WGA-AF) and 489-531 nm (GFP from the endothelium). We’ve removed a small periodic bias (∼17 a.u. with 12-bit sampling) in the output of the PMTs by subtracting the output of a PMT with no photon input from the output of another PMT that recorded the fluorescence^70^. LEA-AF labeling was imaged using a SP5 upright laser scanning microscope (Leica Microsystems) coupled to a MaiTai Ti:Sapphire laser (Spectra-Physics) via a 20× 1.0 NA water-immersion objective and a 560–625 nm bandpass filter (Leica Microsystems). The excitation wavelength was 830 nm.

### Estimation of the point-spread function (PSF) and optical resolution

We characterised the point-spread function (PSF) and the optical resolution of our microscope (*Supplementary Note: Modeling A*) by estimating standard deviation of the Gaussian-like PSF in xy-plane (Extended Data Fig.2abe), 𝜎_𝑥𝑦_ = 149 ± 2 nm (mean ± s.e.m.), and along the objective’s z-axis (Extended Data Fig.2cdf), 𝜎_𝑧_ = 556 ± 5 nm (mean ± s.e.m.). The corresponding optical resolutions were estimated as full width at half maximum (FWHM) of the PSF: FWHM_xy_= 351 ± 4 nm (mean ± s.e.m.) and FWHM_z_= 1310 ± 11 nm (mean ± s.e.m.).

To estimate 𝜎_𝑥𝑦_, we imaged 100-nm-diameter beads (#T14792, ThermoFisher, USA) and fitted their fluorescence intensity distributions in the xy-plane, 𝐼_𝑏𝑒𝑎𝑑_(𝑥, 𝑦), with the following expression, derived in *Supplementary Note: Modeling A* (Eq.A7):

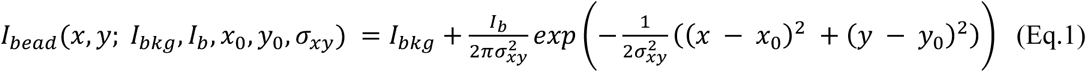

Here 𝐼_𝑏_ and 𝐼_𝑏𝑘g_are fluorescence intensities of the bead in a 2D image and fluorescent background, respectively; (𝑥_0_, 𝑦_0_) are the coordinates of the bead’s center; and 𝜎_𝑥𝑦_ is the standard deviation of the Gaussian PSF in xy-plane. See Extended Data Fig.1ab for the fit examples and *Supplementary Note: Modeling (A)* for preprocessing of the raw images.

To estimate 𝜎_𝑧_, we imaged 200-nm-diameter beads and summed their fluorescence intensity in the xy-planes, obtaining its distribution along z-axis, 𝐼_Z_(𝑧). We then fitted 𝐼_Z_(𝑧) with the following expression, derived in *Supplementary Note: Modeling* (Eq.A10):

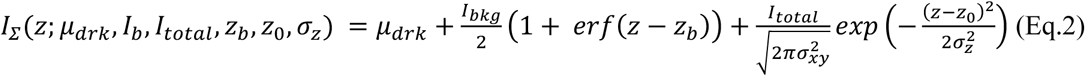

Here 𝜇_𝑑𝑟𝑘_ is the dark PMT output; 𝐼_𝑏𝑘g_ and 𝐼_𝑡𝑜𝑡𝑎𝑙_ are background and total bead fluorescence intensity, respectively (see *Supplementary Note: Modeling (A)* for the difference between 𝐼_𝑡𝑜𝑡𝑎𝑙_ in Eq. 2 and 𝐼_𝑏_ in Eq. 1); 𝑧_𝑏_ is the coordinate of the cover-glass/mounting medium (with beads) interface; 𝑧_0_ is the coordinate of the bead’s center; 𝜎_𝑧_ is the standard deviation of the Gaussian PSF along the z-axis. See Extended Data Fig.2bd for fits examples.

### Image processing for visualization

To facilitate visualization we applied *gaussian blur* and(or) *sigmoid correction* to some of the images shown in the figures. *Gaussian blur* is a low-pass filter, which reduces the noise, e.g. the shot noise. *Sigmoid correction* is a non-linear transformation, I = F(I_0_), of the normalized input pixel intensities, I_0_ (ranging from 0 to 1) using the logistic function:

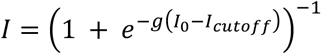

Here, *g* (gain) controls the steepness of the transition, and I_cutoff_ defines the midpoint of the intensity range where contrast is enhanced. The transformation pushes values I_0_ < I_cutoff_ towards 0 and I_0_ > I_cutoff_ towards 1, thereby stretching intensities near I_cutoff_ and improving contrast in that region.

*Gaussian blur* alone was applied to single-color images (either green or red fluorescence), like middle and right columns in Fig.1g with σ = 250 nm (standard deviation of the Gaussian kernel). The combination of *gaussian blur* and *sigmoid correction* was applied to all dual-colored images (green and red fluorescence shown in the same image), like those in Fig.1f-i. We typically used 𝐼_𝑐𝑢𝑡𝑜𝑓𝑓_ = 0.5 and *g* from 5 to 10. Both the gaussian blur and the sigmoid correction were used only for the visualization purposes and not for the data analysis/estimation, unless stated otherwise.

### Glycocalyx maps

#### Step 1: Image collection and pre-processing

We recorded a volume stack (Fig.1b) capturing pial arterioles, penetrating arterioles, pial venules, and ascending venules after injection of WG-AF (2 mice) and LEA-AF (2 mice). The volume stacks were 100–150-µm-deep with 1-2.5 tall (z-axis) and 0.19 µm (LEA-AF) or 0.5 µm (WGA-AF) wide (xy-plane) voxels. The volume stacks were Gaussian-filtered in 3D with the filter’s standard deviations σ_z_=5 µm, σ_xy_=0.5 µm (WGA-AF data), σ_xy_=0.2 µm (LEA-AF data). Next, we applied maximum intensity projection (MIP, Fig.1b) to the volume stack along the z-axis to obtain a 2D image of lectin labeling. Extended Data Fig.2gh explains why the highest fluorescence intensity in glycocalyx is captured in the focal plane bisecting the vessel (white dashed line shows position of the focal plane along z-axis). A MIP, along the z-axis, of the cross-section in Extended Data Fig.2h will, therefore, show two peaks of fluorescence where the focal plane intersected the vessel wall. In 3D, the two peaks will form two parallel bands of high fluorescence, which is exactly how each vessel looks after MIP (Fig. 3ac). Note elevated fluorescence inside each vessel, i.e. between the two bands, compared to the tissue outside of the vessel. This fluorescence originates from out-of-focus fluorescence on the vessel wall above and below the focal plane and from WGA-labeled RBCs.

#### Step 2: Localization of glycocalyx

We analysed the following vessel types: pial arteriole, penetrating arteriole, arteriolar branching point, ascending venule, pial venule, and venular branching point. For a segment of each vessel type, we first drew the initial contour by hand. We then optimized it using an active contour algorithm (*skimage.segmentation.active_contour* in Python’s Skimage library) and visually checked that the new contour traced the local maximum of the glycocalyx fluorescence intensity along a segment of a blood vessel. If it did not, we adjusted the optimization parameters (Supplementary Note: Fine-tuning active contour algorithms for generating glycocalyx maps) and repeated the optimization. We repeated this procedure for all vessels except (a) capillaries, whose glycocalyx fluorescence was corrupted by lectin-labeled RBCs, and (b) segments of venules, with low fluorescence intensities. The benefit of using the active contour algorithm was our ability to localize glycocalyx in regions with almost absent fluorescence (mostly in venules), using the localized glycocalyx in the neighboring regions.

#### Step 3: Generation of glycocalyx maps

We next extracted the fluorescence intensity, 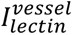, from the 2D image (step #1) at each point (spaced within ∼1 pixel size) along all optimized contours for a specific vessel type and a lectin (WGA or LEA). To facilitate comparison between the lectins, we divided 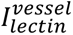 with the mean fluorescence intensity in pial arterioles (pa), 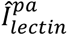, obtaining the relative glycocalyx intensities: 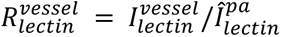 (by definition, the mean relative glycocalyx intensities in pial arterioles, 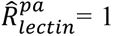). To generate the glycocalyx map, we color-coded 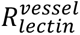 in a new 2D image, at each point of all optimized contours.

### Fluorescence of WGA-AF in capillaries

This protocol was developed specifically to estimate glycocalyx fluorescence intensity in capillaries, where glycocalyx maps (explained in the previous section) could not be estimated, but this protocol also applies to all other vessel types. For a selected vessel type, we recorded a single image (17×17-µm image; 0.067 µm/pixel, 2 µs per pixel) capturing the vessel lumen, the glycocalyx, and the brain tissue outside of the vessel (Fig. 7a), such that the glycocalyx fluorescence band is oriented along the *y*-axis. For pial and penetrating arterioles, we averaged pixel intensities in non overlapping 1-µm-tall blocks along the *y-*axis to obtain 1D profiles of WGA-AF fluorescence. For pial venules and capillaries where the glycocalyx fluorescence intensity was low, we averaged pixel intensities along up to 6-µm-tall *y-*blocks. We excluded 1D profile if (i) the vessel wall was not in focus as judged by its original image in corresponding *y*–block; or (ii) the vessel wall was not straight along the y-axis; or (iii) blood cells were visible (only for capillaries). We then smoothed all remaining fluorescence intensity profiles with a Savitzky-Golay filter (which preserves the peak intensities and shapes) and estimated the glycocalyx fluorescence intensity as the maximum value of the smoothed profile (Inset in Fig. 7b).

### Fluorescence recovery after photobleaching (FRAP)

#### Step 1: Image collection

The FRAP data acquisition pipeline consisted of six sequential phases: *Pre-imaging*, *Baseline*, *Bleaching #1*, *Recovery #1*, *Bleaching #2*, and *Recovery #2*.

The *Pre-imaging* phase consists of (i) imaging a large field of view containing multiple blood vessels; (ii) selecting a ∼8×8 µm region of interest (ROI) on a single pial arteriole (Fig.5a); and (iii) continuously acquiring images of the ROI while adjusting the imaging depth to ensure that the focal plane intersected the arteriole’s cylinder axis, thereby bringing it into focus.

The *Baseline* phase begins when we start recording images of the selected ROI and continue for 1– 2 minutes. From the moment imaging began—already during *Pre-imaging* described above—WGA-AF molecules, both free in the blood plasma and bound to the glycocalyx, began to undergo photobleaching. In a standard FRAP experiment, bleaching during baseline acquisition is typically negligible. However, *in vivo* experiments inherently involve measurable bleaching, driven by the need to collect sufficient photons for good image contrast. Achieving this requires higher laser intensities, which inevitably increase bleaching.

The *Bleaching #1* phase immediately followed the *Baseline* phase, with image acquisition continuing uninterrupted. At this stage, the laser intensity was increased, leading to a higher photobleaching rate, compared to the *Baseline,* and, thus, to a rapid decrease of fluorescence intensity in the glycocalyx. Data acquired during the bleaching phase were not used for further analysis.

The *Recovery #1* phase began immediately after the laser intensity was reduced from the high (*Bleaching*) level back to the *Baseline* level. Due to the preceding bleaching, the fluorescence intensity at the start of recovery was lower than its equilibrium value. During recovery, the fluorescence gradually increased toward equilibrium. The *Bleaching #2* and *Recovery #2* were analogous to *Bleaching #1* and *Recovery #1* but repeated later.

#### Step 2: Splitting recorded movies into five movies, corresponding to the five phases of FRAP

The entire recorded movie is 2-3 min long. We needed to isolate the Baseline, Recovery #1 and #2 phases for fitting. We did that by observing abrupt transitions in fluorescence intensity, like shown in Fig.5b (left). Knowing how long it takes to record a whole image and a single line of the image, we could estimate the time of changing laser intensity with ∼1-2 ms precision (it takes ∼1.1 ms to record one line of an image). With five imaging phases, we estimated four transition times: t_1_, t_2_, t_3_, t_4_. For example, t_1_ corresponded to *Baseline/Bleaching #1* transition (Fig.5b). Next we splitted the movie into five movies (one for each phase) based on the transition times. For simplicity, we did not use images within which the transition occurred, like one in Fig.5b (left). We only used movies corresponding to the *Baseline*, *Recovery #1* and *#2* for further analysis.

#### Step 3: Estimating fluorescence intensity in the plasma, 𝐼_𝑝𝑙𝑠𝑚_

To estimate fluorescence intensity of WGA-AF in the plasma, we (i) selected first images of the two *Recovery* phases, (ii) removed image rows contaminated by the RBCs fluorescence (Fig.5b, middle), and (iii) averaged all pixels in the remaining image rows within a region spanning about 1 µm along the x-axis and was 1-2 µm away from the glycocalyx layer (Fig.5b, right). We used images immediately after the *Bleaching* phases to minimize mixing of the fluorescence from the glycocalyx into the region where we estimated plasma fluorescence. From the two images we had two independent estimates (mean ± s.e.m.) of the 𝐼_𝑝𝑙𝑠𝑚_, which we compared and averaged if the both estimates agreed with a single value. We did not find a decrease in 𝐼_𝑝𝑙𝑠𝑚_ between the *Recovery #1* and *Recovery #2*, which indicated that the concentration of WGA-AF in the blood remained constant during a single, 2-3 min long. FRAP recording. If we could not consistently estimate 𝐼_𝑝𝑙𝑠𝑚_, e.g. because images had not enough contrast to remove rows contaminated by RBCs, we discarded the dataset.

#### Step 4: Estimating fluorescence intensity in the glycocalyx, 𝐼_*g*𝑐𝑥_

We next reduced the noise on individual images by (i) averaging image rows over the 4-µm-tall y-region (40 pixels tall) as shown in Fig.6a (middle), yielding a line-intensity profile along the x-axis, and by (ii) smoothing the line-profile with a moving average with 400-nm window (4-pixels long). The second smoothing ensured that the profile was smooth enough to robustly locate the maximum of fluorescence intensity. From each smoothed line-profile we extracted maximum fluorescence intensity, which we denoted as the glycocalyx fluorescence intensity, 𝐼_*g*𝑐𝑥_, as shown in Fig.6a (right).

#### Step 5: Fitting

First, we averaged estimated 𝐼_*g*𝑐𝑥_ in time, typically in non-overlapping blocks of six values (boxcar averaging with 0.66 s window) and estimated the error bars for the averages (s.e.m.). Second, we fitted the averaged 𝐼_*g*𝑐𝑥_ during the *Baseline*, *Recovery #1*, and *Recovery #2* simultaneously with the following expression:

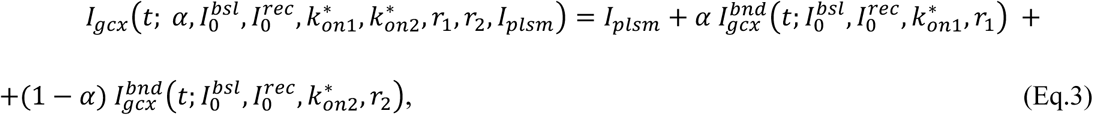

where 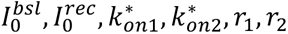 were the fitting parameters and 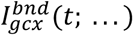 is a combination of single-exponential recovery functions. For the derivation see *Supplementary Note: Modelling B*. Note that we used the plasma fluorescence intensity 𝐼_𝑝𝑙𝑠𝑚_ as a constant estimated in Step #3. The fitting was done with weighted least squares optimization using estimated error bars on individual averaged 𝐼_*g*𝑐𝑥_ values. In some cases, the *Recovery #2* deviated from *Recovery #1*, possibly to a drift of the vessel out-of-focus, in which case we fitted only *Baseline* and *Recovery #1*. Finally we assessed the fit quality by checking for systematic deviations of the standardized residuals from the fitted theory (Fig. 5f). In an ideal fit, standardized residuals (z-score) should be distributed according to the standard normal distribution. Figure 5f demonstrates how some of the residuals deviated by more than four STDs from their mean. In Extended Data Fig.8 we show that these systematic errors were due to vasomotion – dilation/constriction of brain arterioles at ∼0.1 Hz frequency^71^. Constriction/dilation of a vessel, as well as the movement of its center^14^, relative to the focal plane will lead to a fluctuation of the measured fluorescence intensity, which we observed in our data. Fits that showed larger systematic deviations, beyond those caused by the vasomotion, were discarded. Finally, we normalized *k_on_*_1_*, *k_on_*_2_* by dividing them with 𝐼_𝑝𝑙𝑠𝑚_(Fig. 5g). Fig. 6e shows an example fit, where fitted parameters were α = 0.69 ± 0.04, I_0_^bsl^ = 92 ± 8 a.u., I_0_^rec^ = 4.3 ± 2.2 a.u., k*_1_ = 18.4 ± 2.7 s^-1^, k*_2_ = 5.8 ± 0.8 s^-1^, r_1_ = 0.55 ± 0.10 s^-1^, r_2_ = 0.085 ± 0.008 s^-1^.

#### Optional Step: A higher-dose WGA-AF FRAP experiment

As an additional control experiment, we injected a 2.7 times higher dose of WGA-AF (by weight) and performed the FRAP analysis as described above (Extended Data Fig.7cd). In this high-dose experiment we observed the following:

1. WGA-AF fluorescence intensity in the glycocalyx at the start of the FRAP measurement, I_0_^bsl^, measured 268 ± 8 a.u., which is 2.04 ± 0.18 times higher than the average I_0_^bsl^ = 132 ± 11 a.u. (Fig. 5g, averaged over saline- and enzyme-treated mice). This indicates that using our standard dose of WGA-AF does not saturate the free glycocalyx motifs, leaving free glycocalyx motifs available.
2. WGA-AF fluorescence intensity in the plasma increased by 2.67 ± 0.03 times (Extended Data Fig.7cd), in agreement with the elevated dose, validating our method for estimating WGA-AF plasma fluorescence (Fig. 5b, middle).
3. WGA-AF fluorescence intensity after the bleaching, I_0_^rec^ = 15 ± 5 a.u. agreed with the average I_0_^rec^ = 8 ± 2 a.u., estimated in experiments with the standard dose (Fig. 5g, averaged over saline- and enzyme-treated mice). This result indicates that our chosen laser intensity during the bleaching phase bleaches almost all of the WGA-AF bound to the glycocalyx.
4. Estimated parameters returned by fitting Eq.3 to the data, except for I_0_^bsl^ discussed in (1), fall into the range of values reported for the standard WGA-AF dose, indicating robustness of the measured parameters.

### Glycocalyx thickness

#### Step 1: Calculating line-profiles of fluorescence intensity

From the *Bleaching #1* movie we selected the first two frames, starting with the frame with a rapid low-to-high fluorescence intensity transition as in Fig.6b (left). From this two frames we selected ∼1-µm-tall y-regions (approximately 8-10 pixels tall) that (i) showed the lowest amount of WGA-AF-labeled RBCs (seen as bright one-line tall streaks of fluorescence) and (ii) agreed with the fitted theory (explained below). We kept the y-regions ≤ 10-pixel-tall to ensure minimal motion blur, which can artificially increase glycocalyx thickness if not accounted for. The time it takes to record 10 consecutive rows in an image is approximately 11 ms, which is 1/18 of the average heartbeat period, i.e. 200 ms (5 Hz heart rate), in mice we studied. This ensured negligible motion blur due to the pulsations of the vessel’s wall and center^14,68^. In some cases, the selected 10-pixel tall region contained traces of RBCs flowing near the glycocalyx. Then we removed the contaminated rows of the image as we did in Fig.6b (middle). Finally, we averaged all rows in the selected image (after RBC-rows removal) to obtain the profile of average fluorescence intensity and the corresponding error bars (s.e.m.). We always averaged over ≥ 7 rows of pixels to maximize accuracy of the estimated error bars (Fig.7a). The output of this step is shown in Fig.7b as the black dots with error bars.

### Step 2: Fitting the line-profiles of fluorescence intensity

We fitted the line profile obtained in the previous step, with weighted least squares, using the following expression for the expected fluorescence intensity profile:

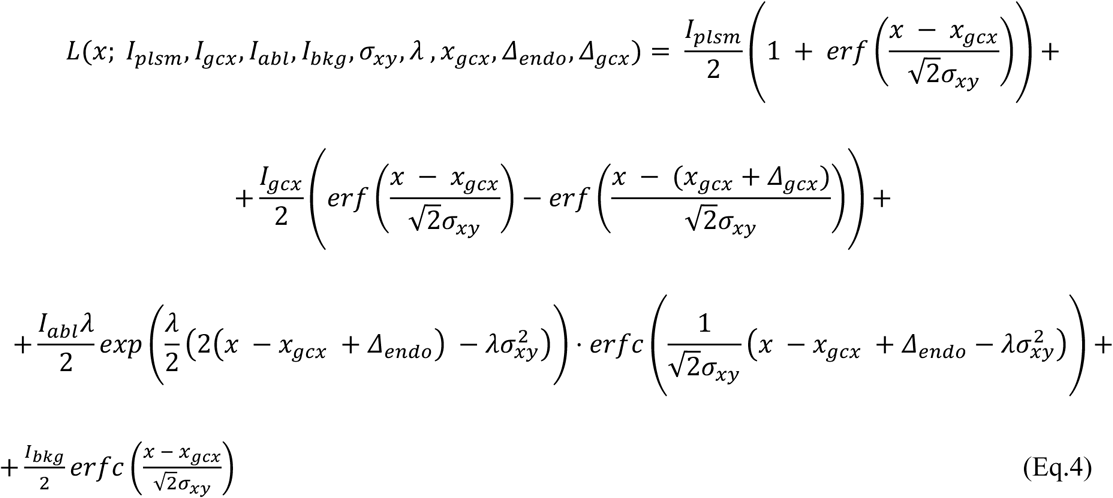

with 𝐼_𝑝𝑙𝑠𝑚_, 𝐼_*g*𝑐𝑥_, 𝐼_𝑎𝑏𝑙_, 𝐼_𝑏𝑘g_, 𝜎_𝑥𝑦_, 𝜆, 𝑥_*g*𝑐𝑥_, 𝛥_𝑒𝑛𝑑𝑜_, 𝛥_*g*𝑐𝑥_ as fitting parameters. For the derivation of Eq.4 see *Supplementary Note: Modeling C* (Eq.C9). Finally we assessed the fit quality by checking for systematic deviations of the standardized residuals from the fitted theory. Fits that showed such deviations were discarded. All fits are shown in Extended Data Fig.8. Note that the estimated STD of the microscope PSF was 175 ± 7 nm, exceeding the estimate of 149 ± 3 nm done with nanobeads (Extended Data Fig. 2e) by only 26 ± 8 nm, which is expected due to aberrations induced by the cranial window and the brain.

### Bayesian statistics

#### Bayesian multilevel models

Our recorded data had nested structure: For a given mouse, we took measurements in different blood vessels, and within each vessel we repeated measurements. If the expected values of the measurable vary from vessel to vessel, repeated measurements within a given vessel will form clusters of measurements. As a result, when pooling data from different mice, data recorded from individual mice can be correlated (not independent). Therefore here we could not apply standard statistical tests, e.g., t-test or ANOVA, which assume independence of the measurements. Here we used Bayesian multilevel generalized linear models (GLMs), which are efficient when analysing small datasets (< 10 measurements) with nested structure^72,73^. Here we followed the statistical analysis described in detail in Fjorbak et al. ^14^. Code implementing the statistical analysis, as well as reproduction instructions, are available at https://github.com/teddygroves/glycocalyx.

#### Choice of priors

We chose priors based to be weakly informative, that is, broadly consistent with the available information about the corresponding effects. We verified that our priors were also jointly consistent with the available information about the range of possible measurements using graphical prior predictive checks (Supplementary Fig. 1-5).

#### Markov chain Monte Carlo (MCMC)

We performed MCMC sampling using adaptive Hamiltonian Monte Carlo as provided by PyMC^74^ via bambi^75^ for models that could easily be expressed using formula notation, and Stan^76^ for the one remaining model. We used the default MCMC settings provided by these programs except for the target acceptance probability and maximum treedepth, which were set to maximise performance while achieving good diagnostics.

#### Test statistic, sign probability, and an example of a statistical test

After finalizing a model, we drew conclusions (analogous to statistical hypothesis testing in the frequentist approach) by defining a test statistic (TS) and examining its posterior distribution, obtained with a Monte Carlo simulation. Certainty in the conclusion (analogous to *p-value* in statistical hypothesis testing in the frequentist approach) was quantified as the sign probability (SP), i.e. the posterior mass on one side of zero, or 𝑚𝑎𝑥(𝑘, 1 − 𝑘) where 𝑘 is the proportion of samples where 𝑇𝑆 > 0. Statistical significance was assumed if SP exceeded the threshold of 0.975 (asterisk (*) in figures). This procedure was used for all statistical comparisons.

For example, in Fig. 3h we compared relative glycocalyx intensities of LEA-AF between arterioles 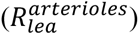 and venules 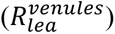. We fitted *model A* (see below) to the estimated 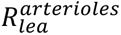 and 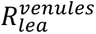, obtained the fitted model’s expected values 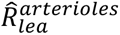 and 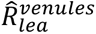, and defined TS = 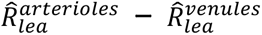. Monte Carlo simulation yielded the TS distribution, shown by its mean and 95% inter-quantile range as the green dot with an error bar in Fig. 3h. All of the Monte-Carlo generated samples were positive, which gave SP = 1.0, higher than the 0.975 threshold. Therefore, we concluded that 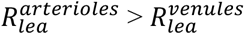.

### Statistical models

Briefly, we constructed five models, explained in detail in *Supplementary Note: Statistical models*. Model choice depended on measurement type, prior knowledge, and model performance. Following the Bayesian Workflow^77^, we began with simple models and iteratively modified components, priors, and hyperparameters to balance descriptive accuracy with computational stability. Each model was fitted by adaptive Hamiltonian Monte Carlo to obtain posterior samples and evaluated based on qualitative fit to the observed data under posterior predictive checking (Supplementary Fig. 1-5). See *Supplementary Note: Statistical models* for details about the models, computation, validation, interpretation of results, and supporting figures.

#### Model A

Model of relative glycocalyx fluorescence intensity (*glycocalyx maps*; Fig. 3). We used a linear regression on log scale, with random intercepts per vessel type and per lectin/vessel type interaction class. Supplementary Note: Statistical models (Model A) describes the implementation of the model and Supplementary Fig.1 shows posterior predictive check (model fit assessment). Code implementing the model, including the priors used, can be found at https://github.com/teddygroves/glycocalyx/blob/main/src/glycocalyx/model.stan.

#### Model B

Model of plasma hyaluronan concentration (Fig. 5e). We used a distributional model to account for treatment-dependent heteroskedasticity. For the expected measurement value we used a linear regression on logarithmic scale, with non-random effects for treatment (Enzyme or Saline) and random effects per mouse. For the expected measurement error we used a linear regression on logarithmic scale with a non-random treatment effect. Supplementary Note: Statistical models (Model B) describes the implementation of the model and Supplementary Fig.2 shows posterior predictive check (model fit assessment). Code implementing the model, including the priors used, can be found at https://github.com/teddygroves/glycocalyx/blob/main/scripts/plasma_hyaluronan.py.

#### Model C

Models of FRAP parameters (Fig. 6). We fit separate linear regression models to each set of parameters after transforming the parameter values to unconstrained space. Each model had non-random treatment effects and random effects for mouse and ROI. Supplementary Note: Statistical models (Model C) describes the implementation of the model and Supplementary Fig.3 shows posterior predictive check (model fit assessment). Code implementing the model, including the priors used, can be found at https://github.com/teddygroves/glycocalyx/blob/main/scripts/gcx_frap.py.

#### Model D

Model of absolute glycocalyx fluorescence intensity (fast imaging protocol; Fig.5d). Linear regression with non-random effects for treatment, vessel type and treatment:vessel type interaction class (i.e. the combination of treatment and vessel type), as well as random effects for mouse and vessel. Supplementary Note: Statistical models (Model D) describes the implementation of the model and Supplementary Fig.4 shows posterior predictive check (model fit assessment). Code implementing the model, including the priors used, can be found at https://github.com/teddygroves/glycocalyx/blob/main/scripts/abs_gcx_intensity.py.

#### Model E

Model of parameters obtained by fitting glycocalyx line-profiles of fluorescence (Fig.7). We used a distributional model due to availability of information about the likely accuracy of the measurements. For the expected measurement value we used a linear regression on log scale with non-random treatment effects and random effects per mouse. For the expected measurement error we used a linear regression on log scale with a non-random, positive-constrained effect for the measurement error predictor. Supplementary Note: Statistical models (Model E) describes the implementation of the model and Supplementary Fig.5 shows posterior predictive check (model fit assessment). Code implementing the model, including the priors used, can be found at https://github.com/teddygroves/glycocalyx/blob/main/scripts/gcx_thickness.py.

## Notes

https://github.com/teddygroves/glycocalyx/tree/main

### Competing Interest Statement

The authors have declared no competing interest.

