## Extended data and Supplementary information for "Two-photon microscopy of brain endothelial glycocalyx uncovers spatial heterogeneity, vesicular transport, and lectin-binding kinetics in the living brain"

[1]Department of Neuroscience, University of Copenhagen, Denmark.
[2]Department of Biomedical Sciences, University of Copenhagen, Denmark.
[3]The Novo Nordisk Foundation Center for Biosustainability, Technical University of Denmark, Denmark.
[4]Department of Odontology, University of Copenhagen, Denmark
[5]Department of Cellular and Molecular Medicine, University of Copenhagen, Denmark
[6]Copenhagen Center for Glycocalyx Research, Department of Cellular and Molecular Medicine, Faculty of Health, University of Copenhagen, Denmark
[7]GLX Analytix ApS, Copenhagen, Denmark
[8]Copenhagen Center for Translational Research, Copenhagen University Hospital, Bispebjerg and Frederiksberg Hospital, Copenhagen, Denmark
[†]These authors contributed equally: Oleg Zhukov, Nikolay P. Kutuzov

December 5, 2025

#### Contents

**Extended Data Figures** 2

**Supplementary Figures** 12

**Supplementary Tables** 17

**Supplementary Note: Modeling** 19

**Supplementary Note: Fine-tuning active contours for glycocalyx maps** 25

**Supplementary Note: Statistical models** 26

#### Extended Data Figures

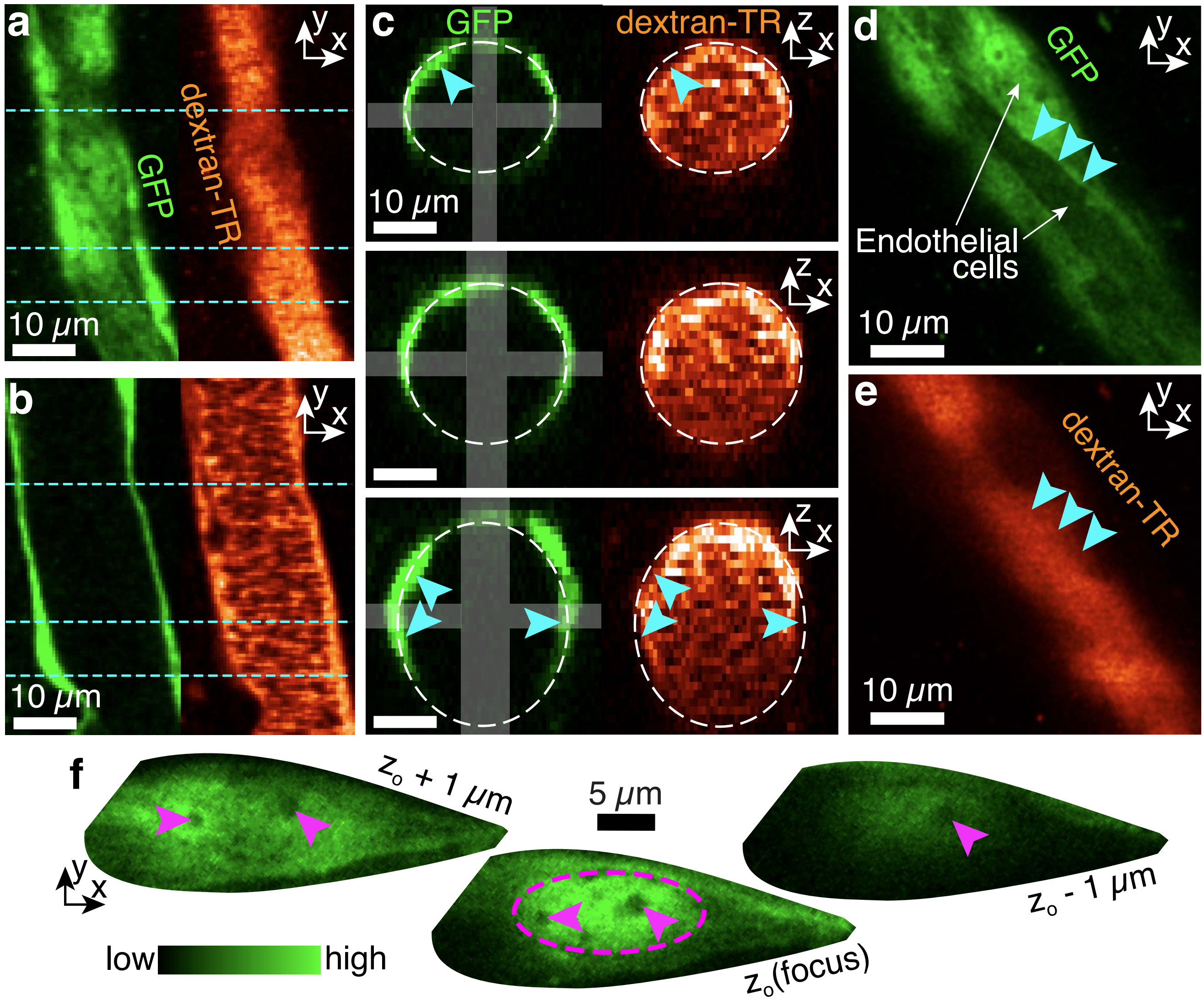

Extended Data Fig. 1: **Two-photon imaging of blood vessels with GFP-labeled endothelium** **(a)** Surface of a pial arteriole (single $xy$-image from a volume stack) showing GFP-labeled endothelium (green) and blood plasma labeled with dextran-TR (orange). **(b)** Same as (a), but imaged through mid-section of the vessel. **(c)** Cross-sections of the arteriole along the dashed cyan lines in (a) and (b). Dashed ellipses indicate lumen diameters estimated from GFP fluorescence in each cross-section. Fluorescence intensity was averaged within the white rectangles to obtain $x$- and $z$-profiles, and diameters were calculated from the distance between intensity peaks. Arrowheads indicate endothelial cell (EC) nuclei protruding into the lumen, altering its cross-sectional shape. The middle row shows an example of a nearly circular cross-section. The fluorescence intensity of both GFP and dextran-Texas Red decreased with depth along the z-axis due to absorption and scattering of the excitation and emitted light by the blood. **(d,e)** Surface of a pial arteriole showing GFP **(d)** and dextran-TR (**e**). Individual ECs can be distinguished by differences in mean GFP intensity and by the presence of nuclei. Arrowheads mark the same intercellular boundary in both images. **(f)** Three images of a single EC acquired at depths separated by 1-μm $z$-steps, showing its nucleus (dashed ellipse) with dark chromatin-rich regions that exclude GFP (arrows).

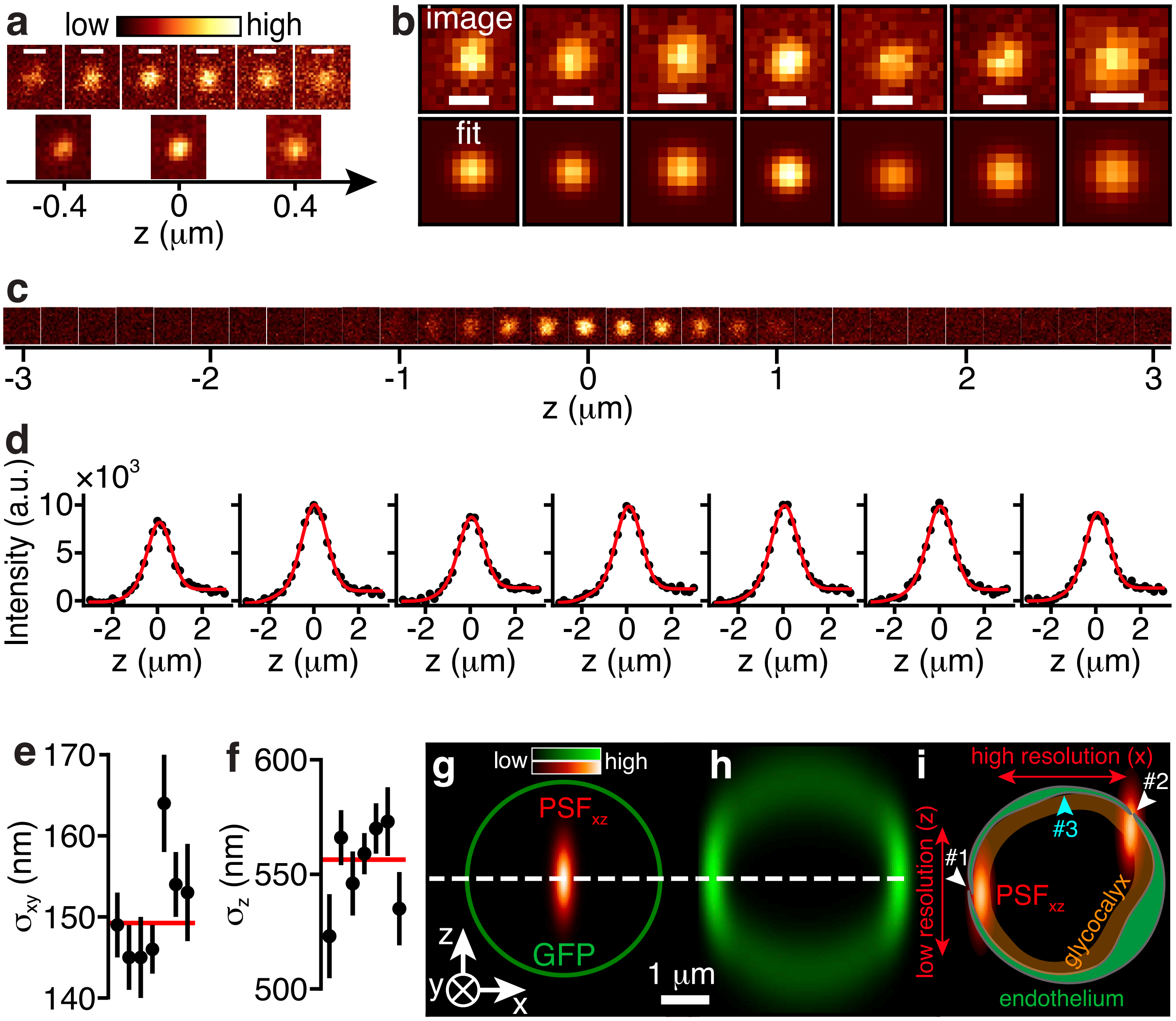

Extended Data Fig. 2: **Point-spread function (PSF) and optical resolution.** **(a)** Top: Images of 100-nm fluorescent beads acquired at different z-depths. Bottom: Processed images obtained by averaging consecutive two frames and 2×2 pixel binning. **(b)** Top row: Averaged images of seven 100-nm beads with the highest mean fluorescence intensity. Bottom row: Fits of Eq. 1. **(c)** Images of a 200-nm bead at different z-depths. **(d)** Analysis of seven image stacks of 200-nm beads (left panel corresponds to bead in (c)). Black dots: summed pixel intensities per z-plane. Red curves: fits of Eq. 2. **(e)** Dots with error bars: Standard deviation of the Gaussian PSF in $xy$-plane, $\sigma_{xy}$, estimated from the seven 100-nm beads. Red line: Weighted average $\sigma_{xy} = 149 \pm 2$ nm (mean ± s.e.m.). The lateral resolution — full width at half maximum (FWHM) — is $351 \pm 4$ nm. **(f)** Dots with error bars: Standard deviation of the Gaussian PSF along $z$-axis, $\sigma_z$, estimated from seven 200-nm beads. Red line: Weighted average $\sigma_z = 556{\pm}5$ (mean ± s.e.m.). The axial resolution (FWHM) is $1310 \pm 11$ nm. **(g)** A circle of uniform fluorescence — e.g. a cross-section of GFP-labeled cylindrical vessel — and the PSF (red). **(h)** Convolution of the circle with the PSF, showing apparent z-axis broadening (compared to x-axis) and bright left and right segments of the circumference, both due to the xz-asymmetry of the PSF. **(i)** A capillary cross-section with two PSFs positioned in the glycocalyx (orange) near contacts sites (white arrowheads #1,2) between two endothelial cells (green), forming capillary wall. Because axial resolution is approximately four times lower than lateral (xy) resolution, detecting local increases in glycocalyx fluorescence at contact sites #1 and #2 is more difficult than at contact site #3 (cyan arrowhead). In addition, variations in fluorescence intensity around the capillary circumference (h) further complicate identification of localized glycocalyx enrichment, in contrast to the top surface segments of large pial arterioles (Fig. 2). Scale bars: (a,b) 500 nm; (c) $1 \times 1$ $\mu$m.

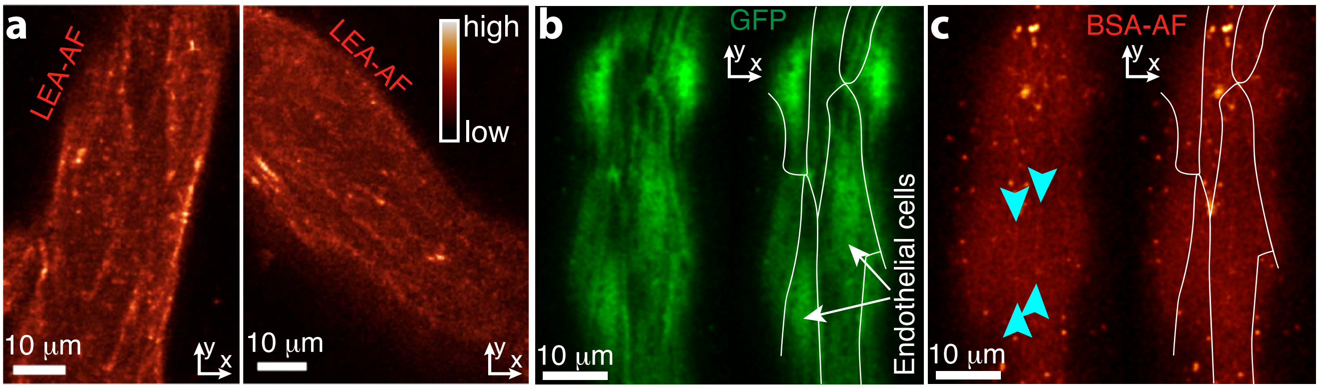

Extended Data Fig. 3: **LEA-AF and BSA-AF labeling of arteriole surfaces.** **(a)** LEA-AF fluorescence at endothelial junctions in two pial arterioles from two different mice. These arterioles are different from that in Fig. 2f. **(b)** GFP fluorescence on the surface of a pial arteriole (left) and the same image with marked ECs perimeters (right). **(c)** BSA-AF fluorescence on the surface of arteriole from (a), showing luminal fluorescence, punctae, and bands of fluorescence near ECs perimeters (arrowheads). The arteriole in (a,b) is different from that in Fig. 2h,i.

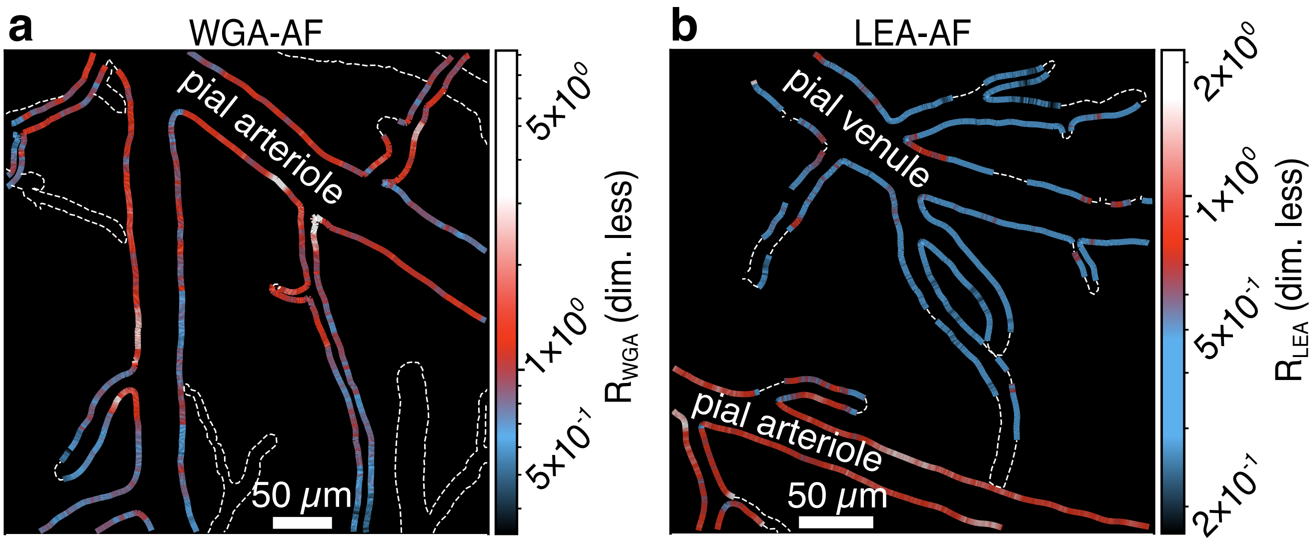

Extended Data Fig. 4: **Glycocalyx maps** for WGA-AF (**a**, $R_{\mathrm{WGA}}$) and LEA-AF (**b**, $R_{\mathrm{LEA}}$), obtained from mice distinct from those in Fig. 3b,d.

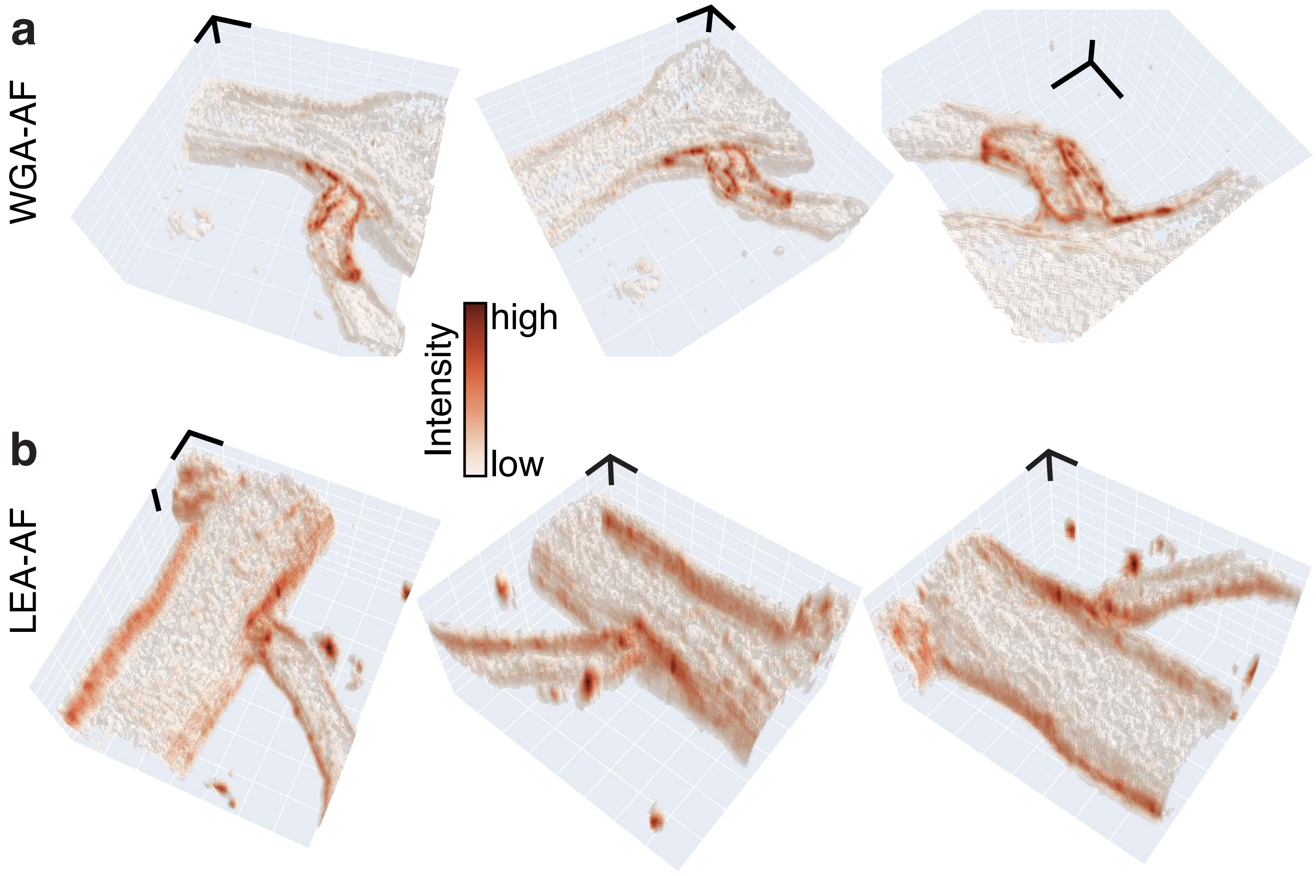

Extended Data Fig. 5: **Volume reconstruction of WGA-AF (a) and LEA-AF (b) labeling at arteriolar branch points**. Each arteriole is shown from three viewing angles. Images are from mice distinct from those in Fig. 4. Scale bars: 10 $\mu$m.

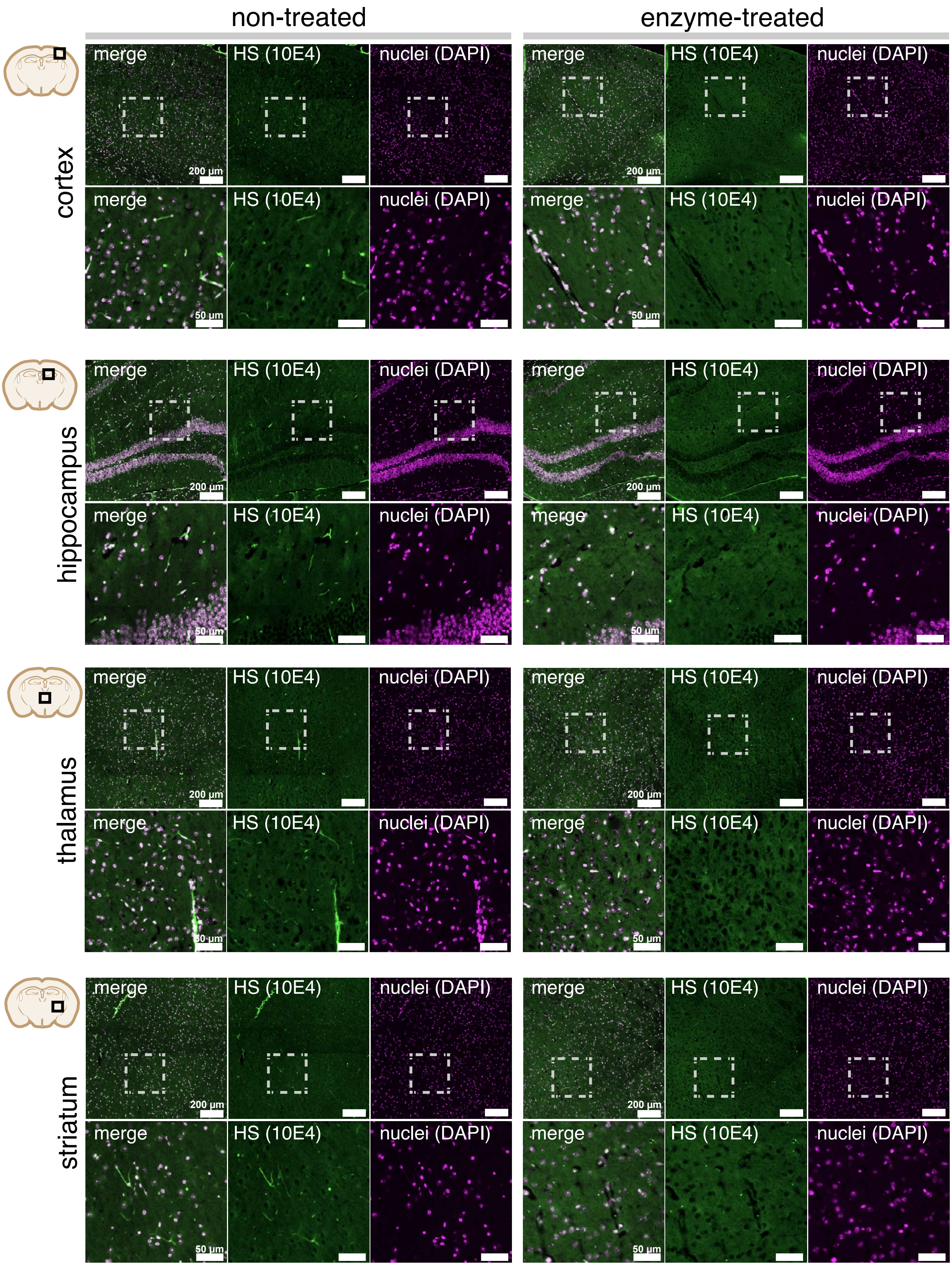

Extended Data Fig. 6: **Immunohistochemical staining of heparan sulfate** with 10E4 antibody in the cortex, hippocampus, thalamus, and striatum of mice treated with saline or enzymes. For each brain region, the top row shows a large field of view, and the bottom row shows a magnified view of the area within the white dashed rectangle. All heparan sulfate images use the same lookup table.

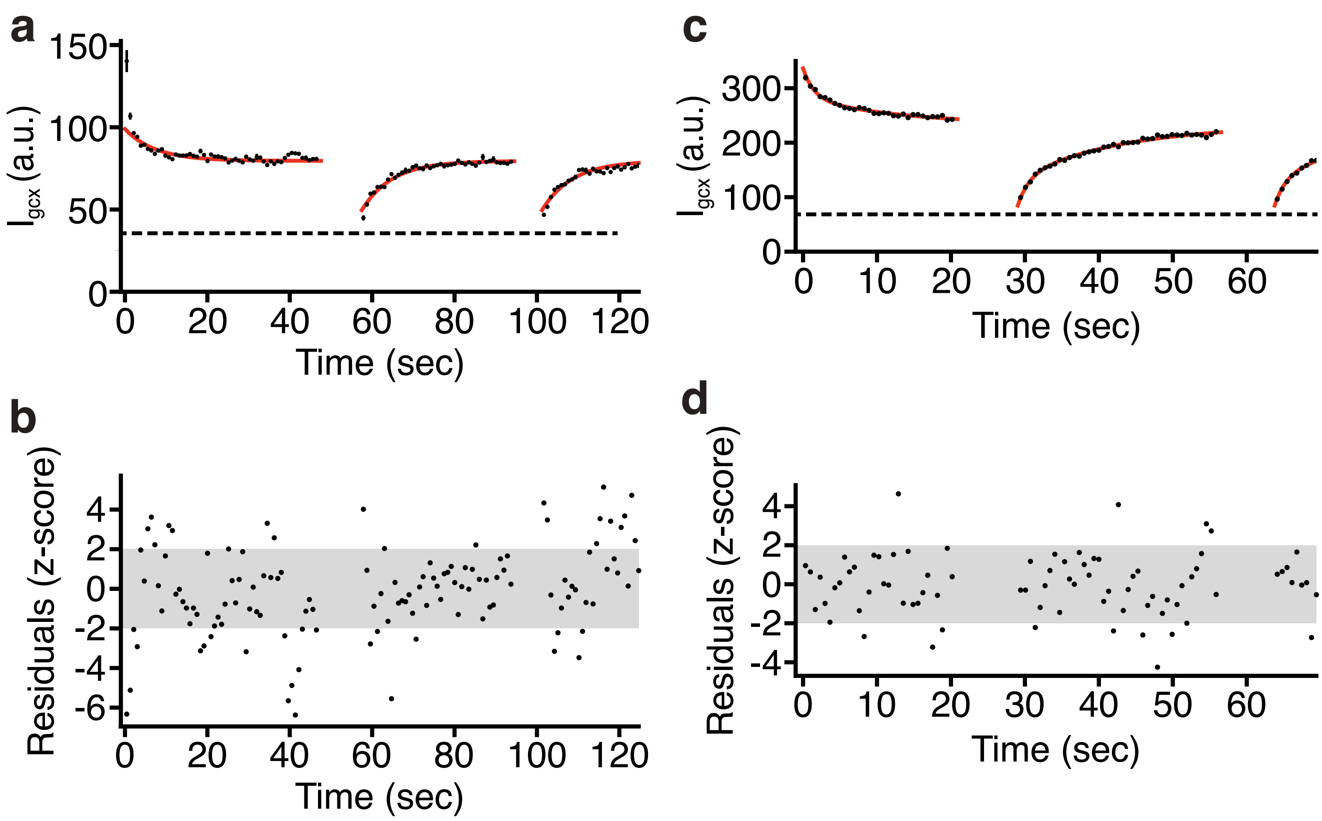

Extended Data Fig. 7: **FRAP analysis with a single binding motif and with a higher WGA-AF dose**. **(a)** Black dots with error bars: WGA-AF fluorescence intensity in the glycocalyx, $I_{\mathrm{gcx}}$ (mean $\pm$ s.e.m. Same data as in Fig. 5e). Red curves: Fits of Eq. (B10), assuming single WGA-AF binding motif (Supplementary Note: Modeling B), to the data. Dashed line: WGA-AF intensity in the plasma. **(b)** Z-scored residuals, i.e. (Fit - Data)/STD, for the fit in (a), where STD are the error bars on the $I_{\mathrm{gcx}}$. For an ideal fit, the residuals should follow the standard normal distribution, with 5% of data points falling outside the gray area. Large deviations from the fitted theory occur at the onset of each phase — Baseline, Recovery #1 and Recovery #2. Because of these discrepancies, we developed a model with two WGA-AF binding motifs (Eq. 4). **(c)** Black dots with error bars: WGA-AF fluorescence intensity in the glycocalyx, $I_{\mathrm{gcx}}$ (mean $\pm$ s.e.m.), measured in an experiment with 2.7 times higher dose (by weight) of WGA-AF. Red curves: Fits of Eq. 4 to the data. Dashed line: WGA-AF intensity in the plasma. Fitted parameters were: $\alpha = 0.416 \pm 0.028$ (dim. less), $I_0^{\mathrm{bsl}} = 268 \pm 8$ a.u., $I_0^{\mathrm{rec}} = 15 \pm 5$ a.u., $k^*_{\mathrm{on1}} = 95 \pm 14$ a.u.$\cdot$s$^{-1}$, $k^*_{\mathrm{on2}} = 13.3 \pm 0.6$ a.u.$\cdot$s$^{-1}$, $r_1 = 0.69 \pm 0.09$ s$^{-1}$, $r_2 = 0.073 \pm 0.003$ s$^{-1}$. Estimated WGA-AF plasma fluorescence intensity was $68 \pm 1$ a.u. **(d)** Z-scored residuals, i.e. (Fit - Data)/STD, for the fit in (a), where STD are the error bars on the $I_{\mathrm{gcx}}$. For an ideal fit, the residuals should follow the standard normal distribution, with 5% of data points falling outside the gray area.

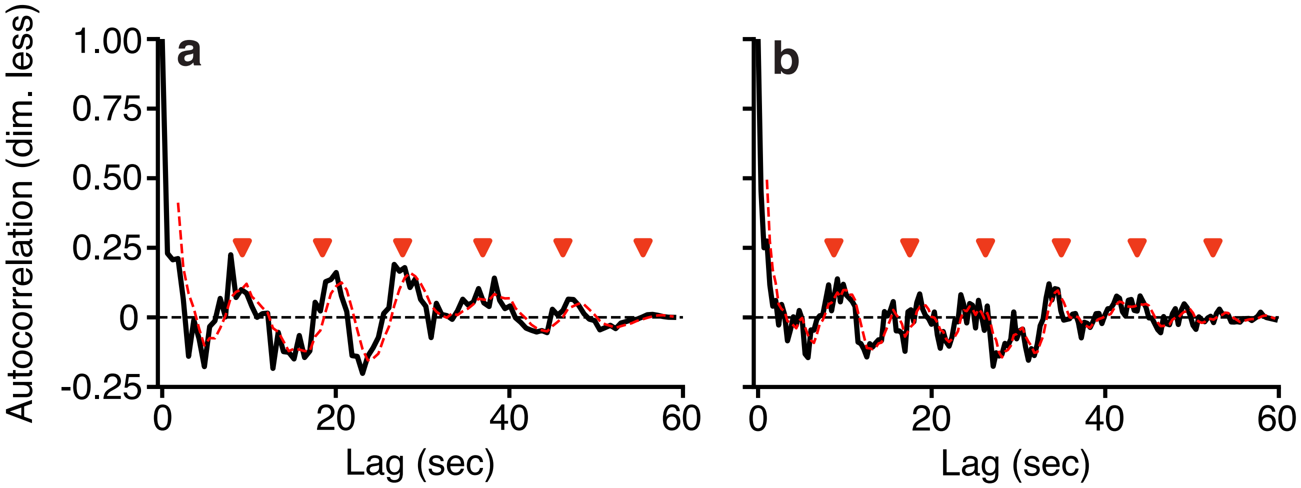

Extended Data Fig. 8: **FRAP traces exhibit oscillations at vasomotion frequencies**. **(a,b)** Black curves: Autocorrelation functions of z-scored residuals (Fig. 5f), obtained from two fits of the FRAP traces in two different mice. Red dashed curves: Smoothed autocorrelation functions (moving average, window size = 2.4 sec). Horizontal dashed line: y = 0. Red triangles: integer multiples of the estimated period, $T_{\mathrm{ac}}$, of the autocorrelation functions: $k \cdot T_{\mathrm{ac}}$ for $k$ from 1 to 6. $T_{\mathrm{ac}}$ was estimated as the average of pairwise differences between time locations of local maxima in the smoothed autocorrelation functions (red dashed curves). We obtained $T_{\mathrm{ac}} = 9.2 \pm 0.5$ s and $T_{\mathrm{ac}} = 8.7 \pm 1.8$ s for (a) and (b), respectively. The two values are consistent with each other and with the typical vasomotion period of $\sim$10 sec (0.1 Hz)[2].

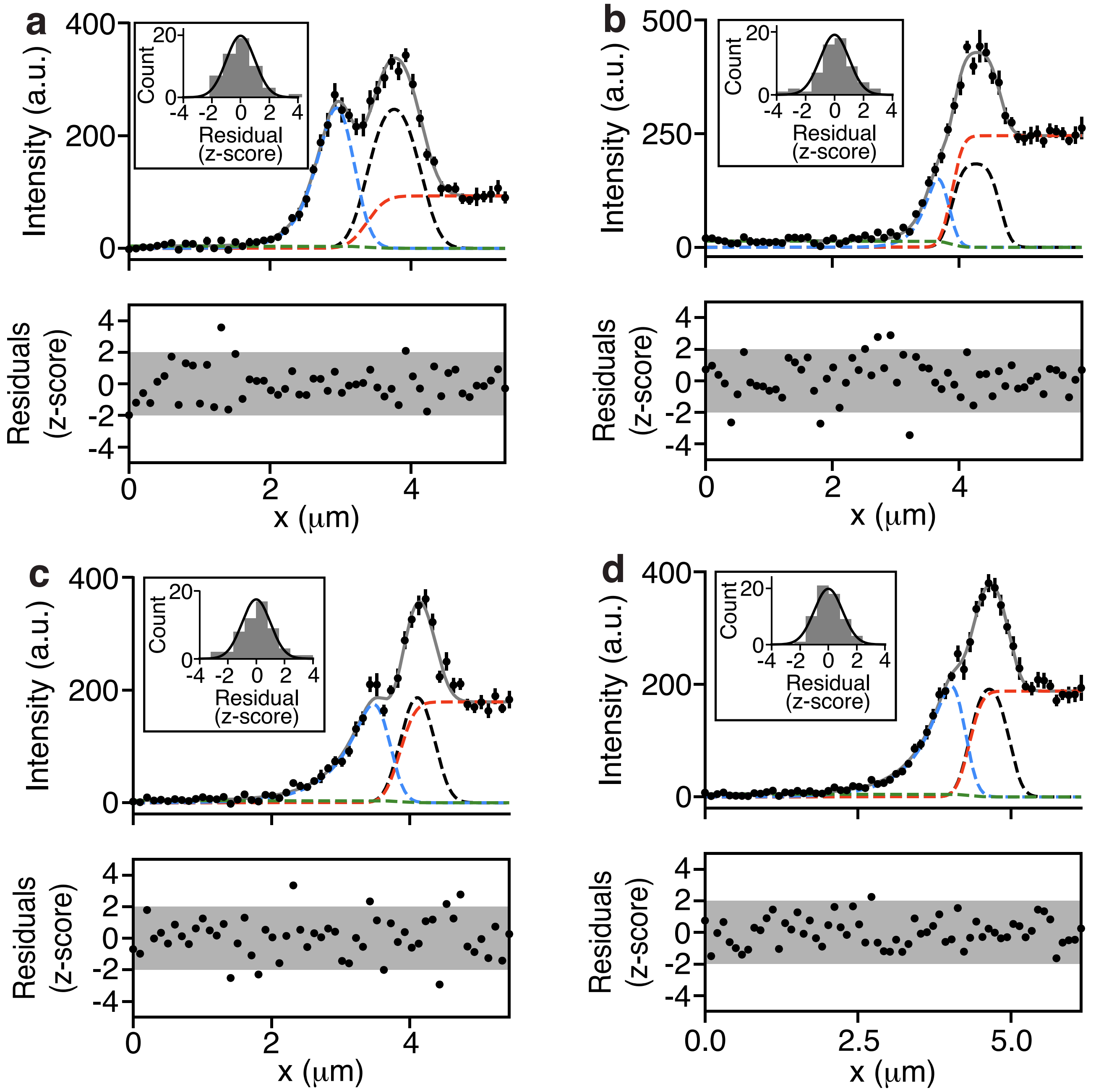

Extended Data Fig. 9: **Estimation of glycocalyx thickness.** (**a-d**) **Top panel**: Fits of Eq. 4 to fluorescence intensity profiles from four different mice (top and bottom rows show example of saline- and enzyme-treated mice). Dots with error bars: Mean ± s.e.m. fluorescence intensities, calculated over columns of pixels (y-axis) in selected images (e.g., Fig. 7a). Gray solid curve: Fit of Eq. 4 to the black data points. Colored dashed curves: Components of the fitted Eq. 4, corresponding to the lumen (red), glycocalyx (black), abluminal region (blue), and background tissue fluorescence (green). The sum of these components yields the gray fitted curve. Inset: Histogram of Z-scored residuals (see below) plotted with the standard normal distribution (black curve). Consistency between the Z-scored residuals and the data implies consistency between the data and the fits. **Bottom panel**: Dots: Z-scored residuals, (data - fit)/s.e.m. estimated from the data in the top panels. Grey shaded zone: [-2, 2] interval along y-axis, which in an ideal fit, should contain, on average, 95% of the measurements.

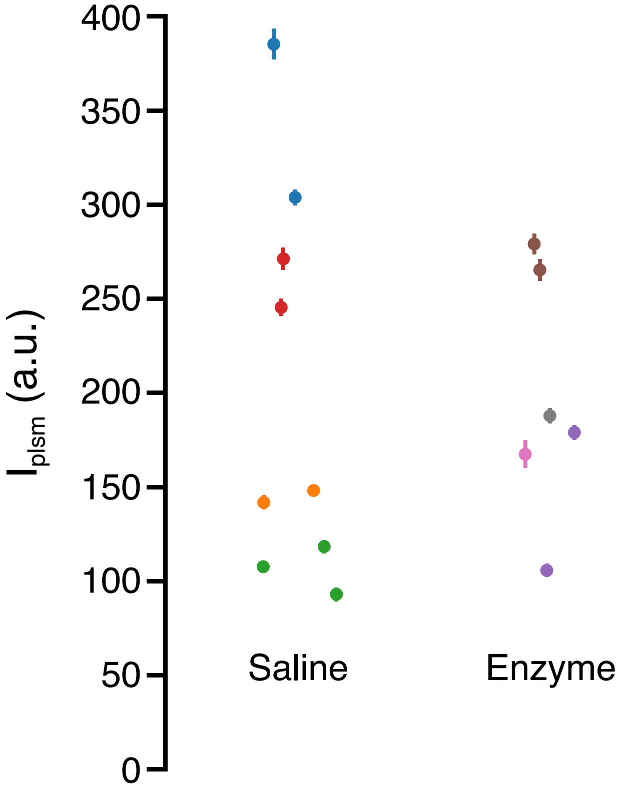

Extended Data Fig. 10: Figure 7g from the manuscript, plotted with error bars on $I_{\mathrm{plsm}}$ values, showing systematic differences between different mice despite the same injected dose of WGA-AF.

#### Supplementary Figures

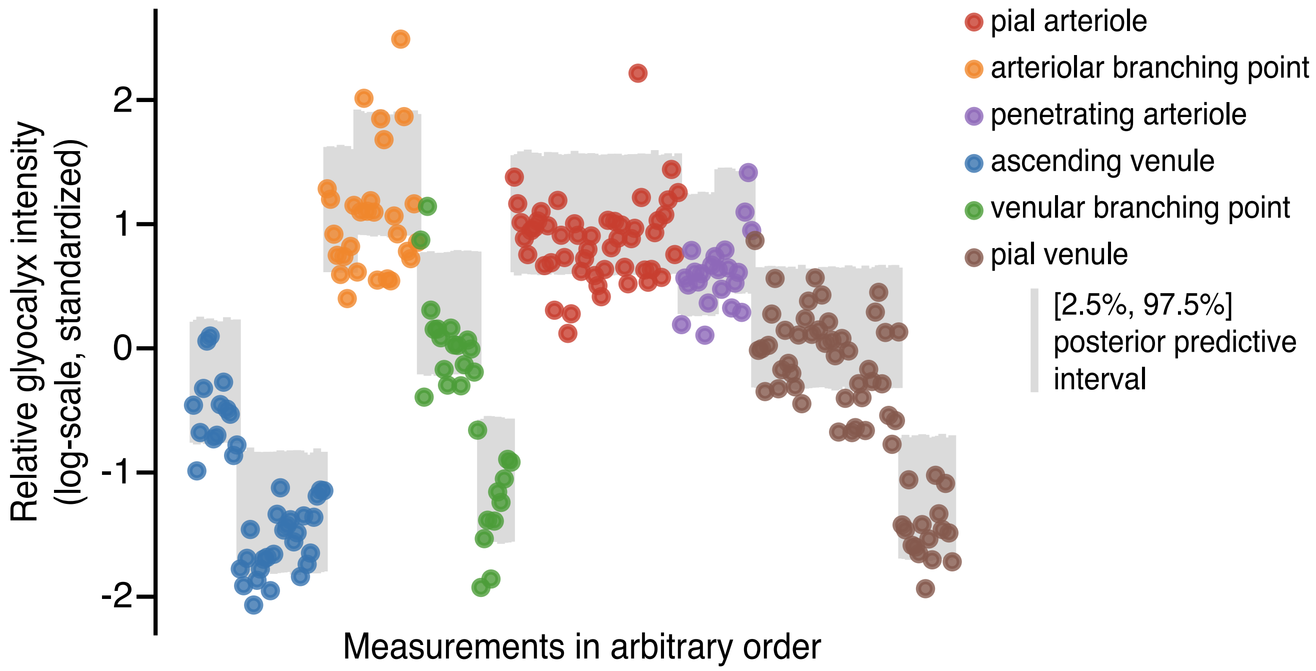

Supplementary Fig. 1: Posterior predictive check for model A (Methods, Eq. D1) of realtive glycocalyx fluorescence intensities shown in glycocalyx maps. For each contour in our dataset (for both WGA-AF and LEA-AF), the plot shows the mean observed measurement (coloured dot) and our model's posterior predictive distribution for this quantity (grey vertical line). Dot colours indicate vessel types. The graph shows general agreement between our model and the observed data, with no particular trend to make better or worse predictions depending on the number of measurements or the vessel type.

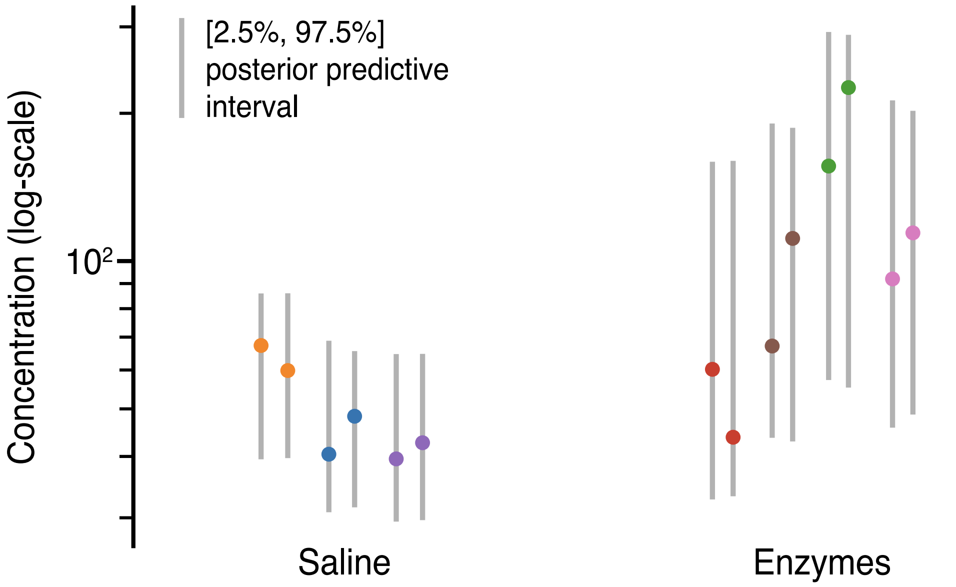

Supplementary Fig. 2: Posterior predictive check for model B (Methods, Eq. D2) of plasma hyaluronan concentration measurements. For each measurement, the coloured dot indicates the observed value and the vertical line summarises our model's posterior predictive distribution. Dot colours indicate mice. Note that the measurements under the enzyme treatment are more dispersed, motivating the use of a distributional model, and that the model was able to capture this heteroskedasticity, as shown by the corresponding wider posterior intervals.

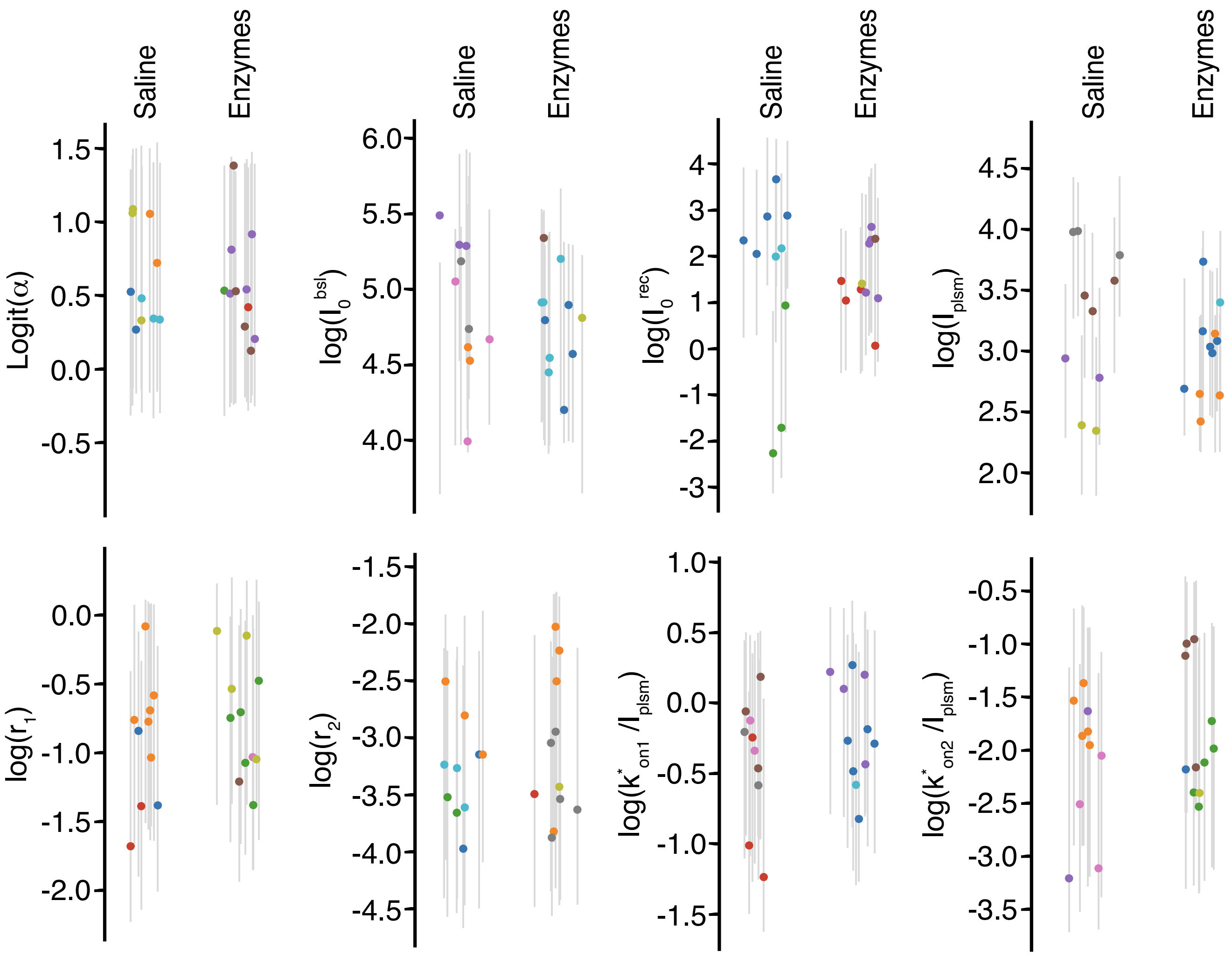

Supplementary Fig. 3: Posterior predictive checks for model C (Methods, Eq. D3) of glycocalyx fluorescence recovery after photobleaching (FRAP) parameters (Fig. 5). For each measurement, the coloured dot indicates the observed value and the vertical line summarises our model's posterior predictive distribution. The graphs show that our models tended to agree with the data, with a slight tendency towards under-sensitivity, as nearly all coloured dots lie well inside their corresponding intervals. This trend suggests that we could have safely used narrower priors, but we did not do so due to the lack of strong information about the target parameters.

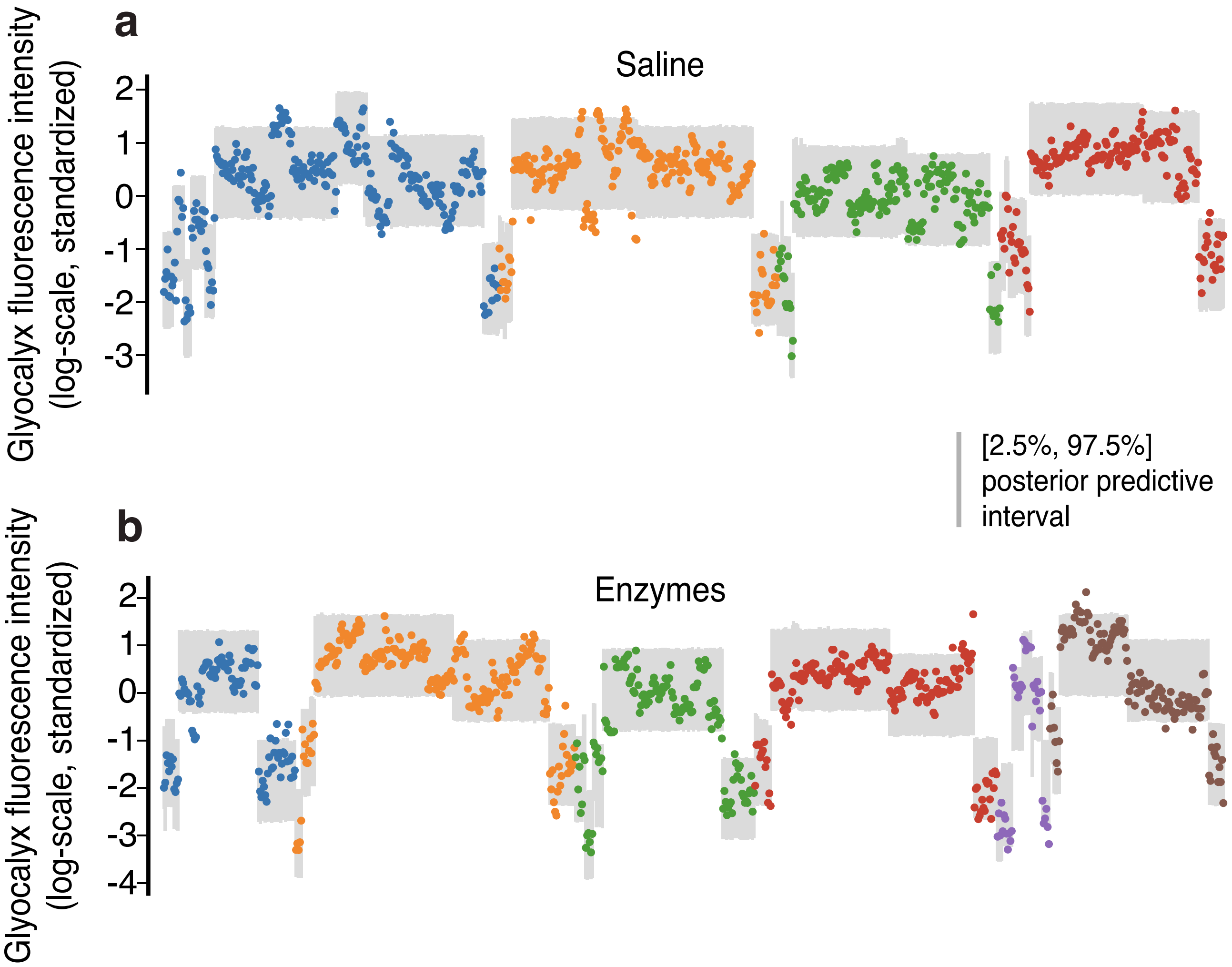

Supplementary Fig. 4: Posterior predictive check for model D (Eq. D4) of glycocalyx fluorescence intensities (Fig. 5cd). For each measurement, the coloured dot indicates the observed value and the vertical line summarises our model's posterior predictive distribution. This graph shows general agreement between the model and observations, with the exception of the mouse with brown dots, some of whose measurements were higher than our model considered plausible. Since there were comparatively few such anomalous measurements we did not update our model to accommodate them.

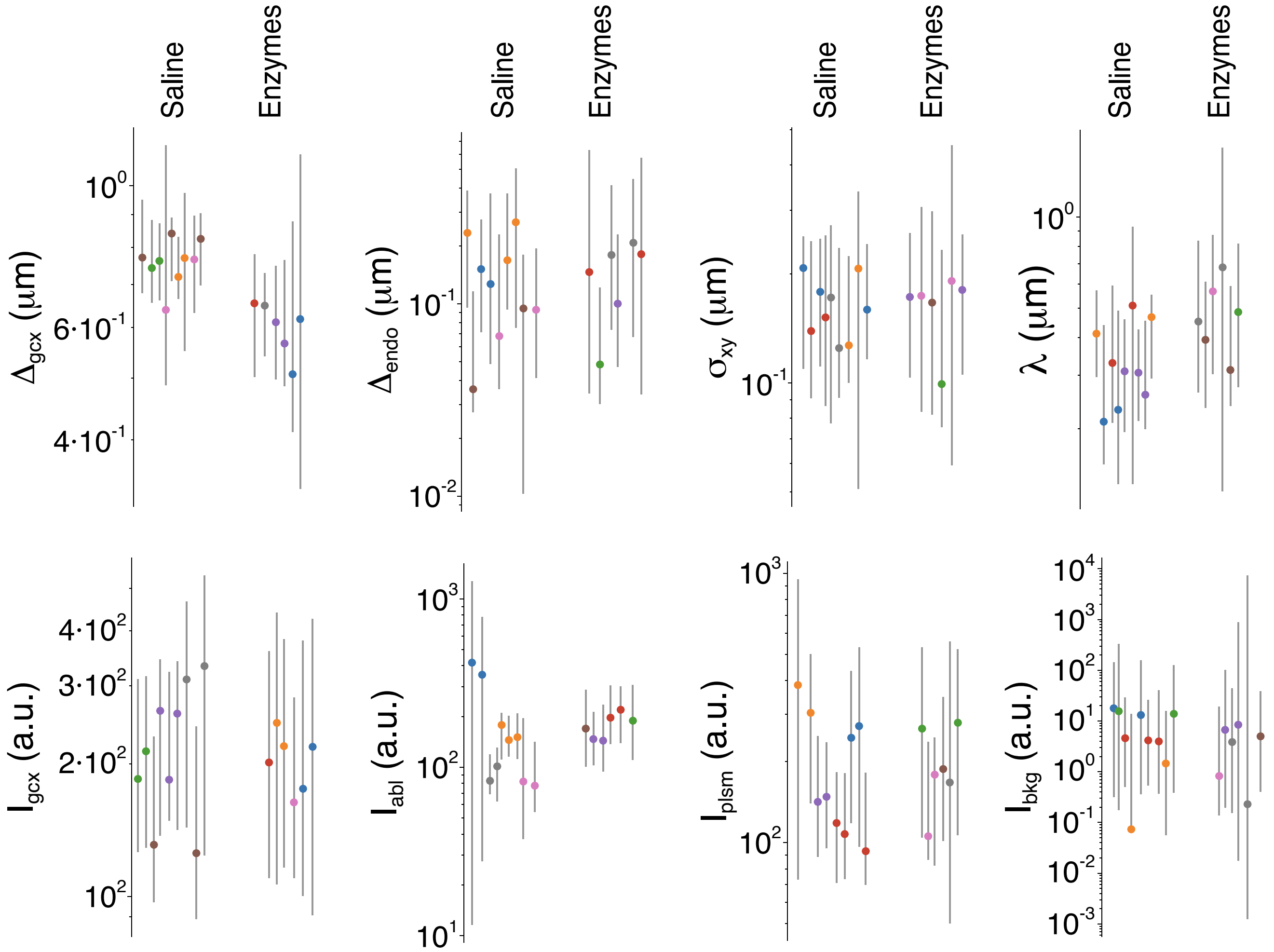

Supplementary Fig. 5: Posterior predictive checks for model E (Eq. D5) of WGA-AF-labeled glycocalyx thickness measurements (Fig. 7). For each measurement, the coloured dot indicates the observed value and the vertical line summarises our model's posterior predictive distribution. The graphs show good overall agreement between the model and data, including capturing distributional information, as shown by the model's appropriately larger and smaller intervals. As with the figure 6 plots, there is a slight tendency towards under-sensitivity to the observations: again we did not use narrower prior distributions as we could not justify this based on the available pre-experimental information.

#### Supplementary Tables

| Parameter | Saline | Enzyme |
|---|---|---|
| $\alpha$ (dim. less) | $0.65 \pm 0.03$ | $0.64 \pm 0.03$ |
| $r_1$ ($s^{-1}$) | $0.44 \pm 0.07$ | $0.50 \pm 0.06$ |
| $r_2$ ($s^{-1}$) | $0.041 \pm 0.06$ | $0.052 \pm 0.010$ |
| $k_1^*/I_{\text{plsm}}$ ($s^{-1}$ a.u.$^{-1}$) | $0.72 \pm 0.09$ | $0.82 \pm 0.09$ |
| $k_2^*/I_{\text{plsm}}$ ($s^{-1}$ a.u.$^{-1}$) | $0.14 \pm 0.02$ | $0.18 \pm 0.03$ |
| $I_0^{\text{bsl}}$ (a.u.) | $144 \pm 19$ | $121 \pm 12$ |
| $I_0^{\text{rec}}$ (a.u.) | $11 \pm 4$ | $6 \pm 1$ |
| $I_{\text{plsm}}$ (a.u.) | $30 \pm 5$ | $21 \pm 2$ |

Supplementary Table. 1: Mean $\pm$ s.e.m. of fitted (Eq. 3) FRAP parameters for saline- and enzyme-treated mice shown in Fig. 5g. $I_{\text{plsm}}$ was estimated independently (not from fitting Eq. 3 to the data) as explained in Fig. 5b.

| Parameter | Saline | Enzyme |
|---|---|---|
| $\Delta_{\text{gcx}}$ (nm) | $760 \pm 20$ | $601 \pm 23$ |
| $\Delta_{\text{endo}}$ (nm) | $138 \pm 25$ | $144 \pm 24$ |
| $\sigma_{xy}$ (nm) | $163 \pm 10$ | $164 \pm 13$ |
| $\lambda$ (nm) | $337 \pm 35$ | $480 \pm 50$ |
| $I_{\text{gcx}}$ (a.u.) | $223 \pm 25$ | $204 \pm 13$ |
| $I_{\text{abl}}$ (a.u.) | $180 \pm 40$ | $204 \pm 13$ |
| $I_{\text{plsm}}$ (a.u.) | $202 \pm 34$ | $197 \pm 26$ |
| $I_{\text{bkg}}$ (a.u.) | $8 \pm 2$ | $4 \pm 1$ |
| $x_{\text{gcx}}$ ($\mu$m) | $3.66 \pm 0.16$ | $3.69 \pm 0.19$ |

Supplementary Table. 2: Mean $\pm$ s.e.m. of fitted (Eq. 4) parameters for saline- and enzyme-treated mice shown in Fig. 7. Note that all calculated values are equally-weighted averages. Because the values of glycocalyx thickness and their error bars, obtained by fitting Eq. 4 to the data, agreed with a single mean (Fig. 7h), we estimated glycocalyx thickness before and after enzymatic treatment as weighted-averages. That gave us glycocalyx thickness $775 \pm 17$ nm and $622 \pm 34$ nm, respectively, which we reported in the paper.

| **Cause** | **Explanation** | **Solution** |
|---|---|---|
| Diffraction limit | An optical image of a point-source of light appears on images as a blurred blob of fluorescence. | Deconvolution, super-localization. |
| Motion blur | Glycocalyx is attached to the vessel walls, which dilate/constrict and change their center position. | Fast imaging, i.e., faster than a fraction of the heartbeat period |
| Labeled RBCs | RBCs, labelled by a dye (e.g. WGA-AF) passing close to the glycocalyx will artificially increase its fluorescence. | Remove RBCs from recorded images (Fig. 5b). |
| Dye in the blood | Dye circulating in blood plasma artificially elevates the fluorescence in the glycocalyx and widens it. | Deconvolution |
| Extravascular dye | Dye fluorescence (e.g. WGA-AF in Fig. 2d) on the abluminal side of the endothelium widens and skews the distribution of fluorescence in the glycocalyx. | Deconvolution and selecting locations with minimal extravascular fluorescence. |
| Curvature of the vessel wall | Widens and skews the fluorescence distribution in the glycocalyx towards the vessel's lumen. | Select large-diameter arterioles or model the effect of the curved wall in capillaries. |
| Vessel surface is not a cylinder | If the vessel wall is not a plane, parallel to the z-axis, the image of the glycocalyx taken in the xy-plane will be artificially blurred. For example, a glycocalyx covering a bulging EC nucleus will appear wider. | Take images in many different locations along a single vessel. Analyse and select locations with the simplest and sharpest fluorescence distribution in the glycocalyx. |

Supplementary Table. 3: Common causes for overestimation of glycocalyx width in *in vivo* experiments.

### Supplementary Note: Modeling

#### A Point-spread function (PSF)

Consider a point source of fluorescence located at $(x, y, z)$, while the laser beam of a two-photon microscope (TPM) is focused at $(x_{\rm psf}, y_{\rm psf}, z_{\rm psf})$. The point-spread function (PSF), denoted $\text{PSF}_{\rm xyz}(x - x_{\rm psf}, y - y_{\rm psf}, z - z_{\rm psf})$, describes the normalized fluorescence intensity at $(x, y, z)$ with the laser focused at $(x_{\rm psf}, y_{\rm psf}, z_{\rm psf})$. In an optically homogeneous, aberration-free, medium, $\text{PSF}_{\rm xyz}$ is typically approximated by a 3D Gaussian function with rotational symmetry about the z-axis [8, 7]:

$$\text{PSF}_{\rm xyz}(x - x_{\rm psf}, y - y_{\rm psf}, z - z_{\rm psf}; \sigma_{xy}, \sigma_z) = G(x - x_{\rm psf}, \sigma_{xy})\, G(y - y_{\rm psf}, \sigma_{xy})\, G(z - z_{\rm psf}, \sigma_z), \tag{A1}$$

where $G$ is a normalized Gaussian centered at zero:

$$G(x; \sigma) = \frac{1}{\sqrt{2\pi\sigma^2}} \exp\left(-\frac{x^2}{2\sigma^2}\right) \tag{A2}$$

Due to light diffraction, when a two-photon microscope (TPM) scans a volume containing a point source of fluorescence located at $(x_0, y_0, z_0)$, the resulting 3D image, $I(x, y, z)$, will not appear as a single bright point but as a distribution of fluorescence surrounding the point-source:

$$I(x, y, z) \propto \text{PSF}_{\rm xyz}(x - x_0, y - y_0, z - z_0; \sigma_{xy}, \sigma_z) = \tag{A3}$$

$$I_{\rm total}\, G(x - x_{\rm psf}, \sigma_{xy})\, G(y - y_{\rm psf}, \sigma_{xy})\, G(z - z_{\rm psf}, \sigma_z) \tag{A4}$$

where $I_{\rm total}$ denotes the total intensity recorded from the point-source. If the point-source is scanned in a single plane, $z = z_{\rm psf}$, the recorded fluorescence intensity in the resulting 2D image is

$$I(x, y) = I_{\rm 2D}\, G(x - x_{\rm psf}, \sigma_{xy})\, G(y - y_{\rm psf}, \sigma_{xy}), \tag{A5}$$

where $I_{\rm 2D} = I_{\rm total}\, G(z_0 - z_{\rm psf}; \sigma_z)$ is the total image intensity, defined as the area under the fluorescence distribution in the image. Thus, $\sigma_{xy}$ can be estimated from a point-source image by fitting it with Eq. A5 using $I_{\rm 2D}$, $\sigma_{xy}$, $x_0$, and $y_0$ as the fitting parameters.

##### PSF: Estimating $\sigma_{xy}$

We imaged a sample containing 100-nm diameter fluorescent beads (#T14792, ThermoFisher TetraSpeck™ Fluorescent Microspheres Size Kit, USA) mounted in a medium. These beads were small enough to appear as point sources (Extended Data Fig. 2b), despite their finite 100-nm diameter, yet large enough to contain sufficient fluorophores for consistently producing high-contrast images (top row in Extended Data Fig. 2a).

Over time, during use and storage, the dye may leak from the beads into the mounting medium. This leakage results in an additional spatially uniform background fluorescence in regions without beads, as the dye equilibrates throughout the medium by diffusion.

To account for this, we fitted the bead images with a model consisting of a constant background term, $I_{\rm bkg}$, and Eq. A5 (with $I_{\rm 2D}$ replaced by $I_b$):

$$I_{\rm bead}(x, y; I_{\rm bkg}, I_b, x_0, y_0, \sigma_{xy}) = I_{\rm bkg} + I_b\, G(x - x_0, \sigma_{xy})\, G(y - y_0, \sigma_{xy}) \tag{A6}$$

$$= I_{\rm bkg} + \frac{I_b}{2\pi\sigma_{xy}^2} \exp\left[-\frac{(x - x_0)^2 + (y - y_0)^2}{2\sigma_{xy}^2}\right] \tag{A7}$$

where $I_b$ is the total bead intensity in the image, and the remaining parameters are defined as in Eq. A5. We fitted Eq. A7 to averaged bead images to reduce noise and enable simpler optimization algorithms (see Methods and Extended Data Fig. 2a).

##### PSF: Estimating $\sigma_z$

In TPM, as in confocal microscopy, $\sigma_z$ is typically several times larger than $\sigma_{xy}$ [14]. Considering this, we could probe the PSF along the $z$-axis using larger beads, 200 nm in diameter, without violating the assumption that the beads act as point sources relative to the PSF's axial extent. The use of 200-nm beads, instead of 100-nm beads, produced brighter images with a higher signal-to-noise ratio due to the increased fluorescence photon yield.

To estimate $\sigma_z$, we recorded volume stacks (Fig. 1c) of images with 200-nm $z$-steps, capturing 200-nm diameter beads as shown in Extended Data Fig. 2(c). We then summed all pixel intensity values in each $xy$-image to obtain the fluorescence intensity profile, $I(z)$, along the $z$-axis:

$$I_\Sigma(z) = \iint_{xy} I(x, y, z)\, dx\, dy = I_{\text{total}}\, G(z - z_0, \sigma_z) \tag{A8}$$

This expression is valid only in the absence of background fluorescence, which is not the case when imaging fluorescent beads. Extended Data Fig. 2(a) (top row) shows that the background fluorescence, $I_{\text{bkg}}$, increases with $z$ as the laser beam moves from the water—used as the immersion medium above the cover glass—into the bead-containing sample. This occurs because the immersion water contains no fluorescent dye, whereas the mounting medium contains dye molecules that have leaked from the beads during storage and imaging (the latter can damage beads and release dye).

Assuming dye diffusion equilibrates its concentration throughout the mounting medium, the free dye concentration away from the beads is spatially uniform in $x$ and $y$ but changes along $z$ as

$$C(z) = C_b\, \mathcal{H}(z - z_b),$$

where $\mathcal{H}$ is the Heaviside step function, $C_b$ is the dye concentration in the mounting medium, and $z_b$ is the $z$-coordinate of the cover glass/mounting-medium interface.

The background fluorescence intensity $I_{\text{bkg}}(z)$ is then given by the convolution of the PSF with the dye concentration distribution:

$$I_{\text{bkg}}(z) \propto (\text{PSF}_{\text{xyz}} * C)(z) = \frac{I_b}{2}\,(1 + \text{erf}(z - z_b)), \tag{A9}$$

where erf denotes the error function, and $I_b$ is the fluorescence intensity of the free dye in the mounting medium. In the presence of background fluorescence, the integrated fluorescence from a bead is

$$I_\Sigma(z; \mu_{\text{drk}}, I_b, I_{\text{total}}, z_b, z_0, \sigma_z) = \mu_{\text{drk}} + \frac{I_b}{2}\,(1 + \text{erf}(z - z_b)) + I_{\text{total}}\, G(z - z_0, \sigma_z), \tag{A10}$$

where $\mu_{\text{drk}}$ is the dark PMT output after bistable-bias subtraction with "Quick-and-noisy" method [6, Section IIC].

#### B Fluorescence recovery after photobleaching (FRAP)

We used the following notation:

| | |
|---|---|
| G | Glycocalyx motif that bind Wheat germ agglutinin (WGA) |
| $W_{\text{AF}}$ | WGA conjugated with Alexa Fluor594 (WGA-AF) |
| $W$ | WGA (i.e. bleached $W_{\text{AF}}$) |
| $GW_{\text{AF}}$ | WGA-AF bound to the glycocalyx (fluorescent) |
| $GW$ | WGA bound to the glycocalyx (non-fluorescent) |
| $[X]$ | Concentration of $X$ |

In our quatitative description of fluorescence recovery in the glycocalyx, we neglected both the radial (blood to glycocalyx) and lateral (unbleached glycocalyx to bleached glycocalyx) diffusion of WGA-AF in the glycocalyx for the following reasons. First, the lowest measured $I_{\text{gcx}}$, immediately after the *Bleaching* was similar to or slightly higher than the intensity of the free WGA-AF in plasma, $I_{\text{plsm}}$ (Fig. 5e). It is unlikely that during the *Bleaching* phase we would never bleach the WGA-AF in the glycocalyx below the $I_{\text{plsm}}$ level, given the spread of $I_0^{\text{rec}}$ values above zero in Fig. 5g. A more plausible explanation is that when we bleach free WGA-AF in the glycocalyx below the plasma level, the radial diffusion of WGA-AF from blood to the glycocalyx will instantaneously equilibrate the WGA-AF concentration in plasma and in the glycocalyx. This is consistent with short diffusion times of BSA molecules (66.5 kDa), ≈0.3 ms, in the lung glycocalyx[9], much shorter than our time resolution, $\Delta t = 0.6$ s (Fig. 5e). While no direct measurements are available for the brain, we estimated diffusion time of ovalbumin (45 kDa) using its diffusion coefficient in rat brain extracellular spaces, $D = 16\ \mu\text{m}^2/\text{sec}$ [10], to estimate the time to diffuse, $\tau$, through a 742-nm thick glycocalyx of brain pial arterioles (Fig. 7c), yielding $\tau = (0.742\ \mu\text{m})^2\ /\ (2{*}16\ \mu\text{m}^2/\text{s}) = 17$ ms, which is 35 times faster than our time resolution.

Second, for a membrane-bound WGA with a lateral diffusion coefficient of $1.5\times10^{-2}\ \mu\text{m}^2/\text{s}$ [4], unbleached WGA-AF would take approximately nine minutes to diffuse four $\mu$m into the bleached region, longer than the entire FRAP recording lasting typically 2–3 min (Fig. 5e). These results reveal a separation of time scales in our FRAP experiments: The radial diffusion equilibrated the concentrations of WGA-AF between blood plasma and glycocalyx so fast we could neglect it and assume that $I_{\text{gcx}}(t)$ was driven by chemical reaction only.

The sequence of chemical reactions occurring during the FRAP measurements is (Fig. 5c):

$$G + W_{\text{AF}} \underset{k_{\text{off}}}{\overset{k_{\text{on}}}{\rightleftarrows}} GW_{\text{AF}} \overset{k_{\text{b}}}{\rightarrow} GW \overset{k_{\text{off}}}{\rightarrow} G, \tag{B1}$$

where the rightmost reaction is effectively irreversible because the second product of the reaction—non-fluorescent WGA (not shown in the scheme)—is rapidly cleared by blood flow.

The corresponding concentration dynamics are described by

$$\begin{aligned} \tfrac{d[G]}{dt} &= 0 \\ \tfrac{d[W_{\text{AF}}]}{dt} &= 0 \\ \tfrac{d[GW_{\text{AF}}]}{dt} &= k_{\text{on}}[W_{\text{AF}}][G] - (k_{\text{off}} + k_{\text{b}})[GW_{\text{AF}}] \\ \tfrac{d[GW]}{dt} &= k_{\text{b}}[GW_{\text{AF}}] - k_{\text{off}}[GW] \end{aligned} \tag{B2}$$

The first equation reflects no net change in the free glycocalyx concentration because it remains in steady state—established during the "Pre-imaging" phase (Fig. 5d), with the total bound WGA, regardless of whether the bound WGA is fluorescent (WGA-AF) or bleached (WGA). Therefore, during the baseline phase, only the concentrations of WGA-AF and WGA vary relative to each other:

$$\frac{d[GW_{\text{AF}}]}{dt} = k_{\text{on}}[W_{\text{AF}}]G_{\text{eq}} - (k_{\text{off}} + k_{\text{b}})[GW_{\text{AF}}], \tag{B3}$$

$$\frac{d[GW]}{dt} = k_{\text{b}}[GW_{\text{AF}}] - k_{\text{off}}[GW]. \tag{B4}$$

We can rewrite the equation for $GW_{\text{AF}}$ in the form:

$$\frac{d[GW_{\text{AF}}]}{dt} = k^*_{\text{on}} - r[GW_{\text{AF}}], \tag{B5}$$

where $\hat{k}_{\text{on}} = k_{\text{on}}W_{\text{AF}}G_{\text{eq}}$ is pseudo zero-order rate constant and $r = k_{\text{off}} + k_{\text{b}}$. Equation B5 has the following solution:

$$[GW_{\text{AF}}](t) = \frac{\hat{k}_{\text{on}}}{r} + \left([GW_{\text{AF}}]_0 - \frac{k^*_{\text{on}}}{r}\right) e^{-rt}, \tag{B6}$$

where $[GW_{\text{AF}}]_0$ is the concentration of $GW_{\text{AF}}$ at zero time.

The FRAP experiment consisted of several phases (Fig. 5de), of which we modeled the *Baseline*, *Recovery 1*, and *Recovery 2* phases. The concentration of $GW_{\text{AF}}$ for these three phases can be expressed as:

$$\begin{aligned} [GW_{\text{AF}}](t) = \big(1 - \mathcal{H}(t - t_1)\big)\left(\frac{\hat{k}_{\text{on}}}{r} + \left([GW_{\text{AF}}]_0^{\text{bsl}} - \frac{\hat{k}_{\text{on}}}{r}\right) e^{-r(t-t_1)}\right) + \\ +\big(\mathcal{H}(t - t_2)(1 - \mathcal{H}(t - t_3))\big)\left(\frac{\hat{k}_{\text{on}}}{r} + \left([GW_{\text{AF}}]_0^{\text{rec}} - \frac{\hat{k}_{\text{on}}}{r}\right) e^{-r(t-t_2)}\right) + \\ +\mathcal{H}(t - t_4)\left(\frac{\hat{k}_{\text{on}}}{r} + \left([GW_{\text{AF}}]_0^{\text{rec}} - \frac{\hat{k}_{\text{on}}}{r}\right) e^{-r(t-t_4)}\right), \end{aligned} \tag{B7}$$

where $t_1$, $t_2$, $t_3$, and $t_4$ denote the start time of the *Baseline*, the start time of *Recovery 1*, the end time of *Recovery 1*, and the start time of *Recovery 2*, respectively. $[GW_{\mathrm{AF}}]_0^{\mathrm{rbsl}}$ and $[GW_{\mathrm{AF}}]_0^{\mathrm{rec}}$ represent the concentrations of $GW_{\mathrm{AF}}$ at the start of the baseline and at the start of both recovery phases, respectively (we assume the same initial concentration after the two consecutive bleaching phases).

Assuming that the fluorescence intensity of $GW_{\mathrm{AF}}$, $I_{\mathrm{gcx}}^{\mathrm{bnd}}$, is proportional to its concentration,

$$I_{\mathrm{gcx}}^{\mathrm{bnd}}(t; I_0^{\mathrm{bsl}}, I_0^{\mathrm{rec}}, k_{\mathrm{on}}^*, r) \propto [GW_{\mathrm{AF}}](t); \tag{B8}$$

$$\begin{aligned} I_{\mathrm{gcx}}^{\mathrm{bnd}}(t; I_0^{\mathrm{bsl}}, I_0^{\mathrm{rec}}, k_{\mathrm{on}}^*, r) = \big(1-\mathcal{H}(t-t_1)\big)\left(\frac{k_{\mathrm{on}}^*}{r} + \left(I_0^{\mathrm{bsl}} - \frac{k_{\mathrm{on}}^*}{r}\right)e^{-r(t-t_1)}\right)+ \\ +\big(\mathcal{H}(t-t_2)(1-\mathcal{H}(t-t_3))\big)\left(\frac{k_{\mathrm{on}}^*}{r} + \left(I_0^{\mathrm{rec}} - \frac{k_{\mathrm{on}}^*}{r}\right)e^{-r(t-t_2)}\right)+ \\ +\mathcal{H}(t-t_4)\left(\frac{k_{\mathrm{on}}^*}{r} + \left(I_0^{\mathrm{rec}} - \frac{k_{\mathrm{on}}^*}{r}\right)e^{-r(t-t_4)}\right), \end{aligned} \tag{B9}$$

where $k_{\mathrm{on}}^* \propto \hat{k}_{\mathrm{on}}$. Note that in the list of parameters in $I_{\mathrm{gcx}}^{\mathrm{bnd}}$ in the equation above, we have listed $t_1$, $t_2$, $t_3$, and $t_4$ separately, as these were not fitted to the data but instead estimated beforehand from the measurements (Fig. 5b).

The total fluorescence intensity, $\hat{I}_{\mathrm{gcx}}$, measured in the glycocalyx is the sum of the fluorescence intensity from free WGA-AF, $I_{\mathrm{plsm}}$, and from glycocalyx-bound WGA-AF, $I_{\mathrm{gcx}}^{\mathrm{bnd}}$:

$$\hat{I}_{\mathrm{gcx}}(t; I_0^{\mathrm{bsl}}, I_0^{\mathrm{rec}}, k_{\mathrm{on}}^*, r, I_{\mathrm{plsm}}) = I_{\mathrm{plsm}} + I_{\mathrm{gcx}}^{\mathrm{bnd}}(t; I_0^{\mathrm{bsl}}, I_0^{\mathrm{rec}}, k_{\mathrm{on}}^*, r) \tag{B10}$$

Note that $I_{\mathrm{plsm}}$ was independently estimated, and only four parameters—$I_0^{\mathrm{bsl}}$, $I_0^{\mathrm{rec}}$, $k_{\mathrm{on}}^*$, and $r$—were fitted to the FRAP data.

To improve the fit to the experimental data, we expanded the model to include two independent glycocalyx motifs capable of binding WGA-AF:

$$\begin{aligned} I_{\mathrm{gcx}}(t; \alpha, I_0^{\mathrm{bsl}}, I_0^{\mathrm{rec}}, k_{\mathrm{on1}}^*, k_{\mathrm{on2}}^*, r_1, r_2, I_{\mathrm{plsm}}) = I_{\mathrm{plsm}}+ \\ \alpha\, I_{\mathrm{gcx}}^{\mathrm{bnd}}(t; I_0^{\mathrm{bsl}}, I_0^{\mathrm{rec}}, k_{\mathrm{on1}}^*, r_1)+ \\ (1-\alpha)\, I_{\mathrm{gcx}}^{\mathrm{bnd}}(t; I_0^{\mathrm{bsl}}, I_0^{\mathrm{rec}}, k_{\mathrm{on2}}^*, r_2), \end{aligned} \tag{B11}$$

where $\alpha$ is the fraction of WGA-AF binding motif #1, characterized by parameters $k_{\mathrm{on1}}^*$ and $r_1$, while $(1-\alpha)$ is the fraction of WGA-AF binding motif #2, characterized by parameters $k_{\mathrm{on2}}^*$ and $r_2$. We fitted Eq. B11 to the FRAP data with $I_{\mathrm{plsm}}$ fixed (estimated independently) and optimized the following parameters: $\alpha$, $I_0^{\mathrm{bsl}}$, $I_0^{\mathrm{rec}}$, $k_{\mathrm{on1}}^*$, $k_{\mathrm{on2}}^*$, $r_1$, and $r_2$.

#### C Glycocalyx thickness

Consider a blood vessel in the microscope's coordinate system $(x, y, z)$, in which an image (Fig. 1ab) is recorded. We made the following assumptions: (i) the vessel's cylindrical axis lies in the focal plane of the objective ($xy$-plane); (ii) the vessel's cylindrical axis is parallel to the $y$-axis; (iii) the vessel's diameter is constant along the $y$-axis.

The first assumption holds for pial arterioles, which lie on the brain surface. The second assumption was satisfied by rotating the microscope's coordinate system so that its $y$-axis aligned (visually, by eye) with the arteriole's cylindrical axis. The third assumption was ensured by selecting regions of arterioles without visible bulging nuclei of endothelial cells.

These three assumptions allow us to exploit symmetry along the $y$-axis and consider only the vessel's cross-section in the $xz$-plane. Furthermore, since the typical pial arteriole radius $R \approx 15$–$20$ $\mu$m is much larger than the PSF's axial extent, $\sigma_z = 0.351$ $\mu$m (Extended Data Fig. 1f), we can neglect the curvature of the vessel wall within the focal volume from which fluorescence is recorded (Extended Data Fig. 1g). In other words, within the focal volume, the vessel wall appears as a plane (rather than a curved cylindrical surface) perpendicular to the recorded $xy$-image plane.

As a result, the model reduces to a single spatial dimension—the $x$-axis in the recorded images. We selected $\approx 1$–$2$ $\mu$m-tall segments from each image, removed rows contaminated by RBCs, and averaged the remaining rows to obtain a fluorescence line profile, $L(x)$, similarly to how we did that in Fig. 5b (midddle).

We define $L(x)$ as the sum of four components: $L_{\text{plsm}}$, $L_{\text{gcx}}$, $L_{\text{abl}}$, and $L_{\text{bkg}}$.

$$L(x) = L_{\text{plsm}}(x) + L_{\text{gcx}}(x) + L_{\text{abl}}(x) + L_{\text{bkg}}(x), \tag{C1}$$

**Component #1: WGA-AF in the lumen.** Let $x_0$ denote the position of the endothelium–lumen boundary. The concentration of free blood WGA-AF, $C_{\text{plsm}}$, is zero outside the vessel ($x < x_0$) and constant throughout the vessel's lumen ($x \geq x_0$):

$$C_{\text{plsm}} \propto \mathcal{H}(x - x_0),$$

where $\mathcal{H}(x)$ is the Heaviside step function.

The corresponding fluorescence of luminal WGA-AF is given by

$$L_{\text{plsm}}(x; I_{\text{plsm}}, x_0, \sigma_{xy}) = I_{\text{plsm}} \left(G(x, \sigma_{xy}) * \mathcal{H}(x - x_0)\right) = I_{\text{plsm}} \frac{1}{2} \left[1 + \operatorname{erf}\left(\frac{x - x_0}{\sqrt{2}\,\sigma_{xy}}\right)\right], \tag{C2}$$

where $*$ denotes convolution and $I_{\text{plsm}}$ is the fluorescence intensity of WGA-AF in the lumen. Here, we use the 1D Gaussian PSF, $G(x, \sigma_{xy})$, instead of the full 3D PSF $\text{PSF}_{\text{xyz}}(x, y, z)$ because the symmetries described above reduce the problem to a single spatial dimension.

**Component #2: WGA-AF in the glycocalyx.** We model the glycocalyx as a homogeneous layer of thickness $\Delta_{\text{gcx}}$, with uniform fluorescence intensity $I_{\text{gcx}}$ for $x \in [x_0,\, x_0 + \Delta_{\text{gcx}}]$ and zero elsewhere. The corresponding fluorescence profile can be expressed as

$$L_{\text{gcx}}(x; I_{\text{gcx}}, x_0, \sigma_{xy}, \Delta_{\text{gcx}}) = I_{\text{gcx}} \left[G(x, \sigma_{xy}) * \left(\mathcal{H}(x - x_0) - \mathcal{H}(x - (x_0 + \Delta_{\text{gcx}}))\right)\right] \tag{C3}$$

$$= \frac{I_{\text{gcx}}}{2} \left[\operatorname{erf}\left(\frac{x - x_0}{\sqrt{2}\,\sigma_{xy}}\right) - \operatorname{erf}\left(\frac{x - (x_0 + \Delta_{\text{gcx}})}{\sqrt{2}\,\sigma_{xy}}\right)\right], \tag{C4}$$

where $*$ denotes convolution.

Here, we assume the partition coefficient (PC) of WGA-AF in the glycocalyx [5] to be exactly one, such that $L_{\text{plsm}}(x > x_0) = I_{\text{plsm}}$. In reality, the partition coefficient for WGA-AF is expected to be less than one, but we could not independently estimate it as in [5], due to binding of WGA-AF to the glycocalyx.

**Component #3: WGA-AF near the abluminal side of the endothelium.** We modeled the endothelium as a layer of thickness $\Delta_{\text{endo}}$, spanning $x \in [x_0 - \Delta_{\text{endo}},\, x_0]$. We assume that no WGA-AF is present within the endothelium itself, as suggested by Fig. 2d, and that the WGA-AF fluorescence in the extravascular space decays exponentially away from the abluminal surface at $x = x_0 - \Delta_{\text{endo}}$. This exponential decay was inferred from the approximately exponential decrease in fluorescence intensity observed in the extravascular region (data not shown).

The fluorescence contribution from this abluminal region can thus be expressed as

$$L_{\text{abl}}(x; I_{\text{abl}}, x_0, \Delta_{\text{endo}}, \lambda, \sigma_{xy}) = I_{\text{abl}} \left[G(x, \sigma_{xy}) * \left(e^{-\frac{x_0 - \Delta_{\text{endo}} - x}{\lambda}} \left(1 - \mathcal{H}(x - x_0)\right)\right)\right] \tag{C5}$$

$$= \frac{I_{\text{abl}}\lambda}{2} \exp\left[\frac{\lambda}{2}\left(2(x - x_0 + \Delta_{\text{endo}}) - \lambda\sigma_{xy}^2\right)\right] \operatorname{erfc}\left(\frac{(x - x_0 + \Delta_{\text{endo}}) - \lambda\sigma_{xy}^2}{\sqrt{2}\,\sigma_{xy}}\right), \tag{C6}$$

where $\lambda$ is the exponential decay length, erfc denotes the complementary error function, and $*$ denotes convolution.

**Component #4: Uniform WGA-AF fluorescence in the extravascular space.** The extravascular space contains unspecific brain autofluorescence and the dark PMT output after bistable-bias subtraction using the "Quick-and-noisy" method [6, Section IIC]. Both contributions have a constant value for $x < x_0$ and are zero inside the vessel. This background fluorescence component can be expressed as

$$L_{\text{bkg}}(x; I_{\text{bkg}}, x_0, \sigma_{xy}) = I_{\text{bkg}} \left[G(x, \sigma_{xy}) * \left(1 - \mathcal{H}(x - x_0)\right)\right] \tag{C7}$$

$$= \frac{I_{\text{bkg}}}{2} \operatorname{erfc}\left(\frac{x - x_0}{\sqrt{2}\,\sigma_{xy}}\right), \tag{C8}$$

where erfc denotes the complementary error function and $*$ is convolution.

**Full fluorescence profile.** Summing the contributions from all four components gives

$$\begin{aligned} L(x; I_{\text{plsm}}, I_{\text{gcx}}, I_{\text{abl}}, I_{\text{bkg}}, \sigma_{xy}, \lambda, x_0, \Delta_{\text{endo}}, \Delta_{\text{gcx}}) = {} & \frac{I_{\text{plsm}}}{2}\left[1 + \operatorname{erf}\left(\frac{x - x_0}{\sqrt{2}\,\sigma_{xy}}\right)\right] \\ & + \frac{I_{\text{gcx}}}{2}\left[\operatorname{erf}\left(\frac{x - x_0}{\sqrt{2}\,\sigma_{xy}}\right) - \operatorname{erf}\left(\frac{x - (x_0 + \Delta_{\text{gcx}})}{\sqrt{2}\,\sigma_{xy}}\right)\right] \\ & + \frac{I_{\text{abl}}\lambda}{2} \exp\left[\frac{\lambda}{2}\left(2(x - x_0 + \Delta_{\text{endo}}) - \lambda\sigma_{xy}^2\right)\right] \\ & \times \operatorname{erfc}\left(\frac{(x - x_0 + \Delta_{\text{endo}}) - \lambda\sigma_{xy}^2}{\sqrt{2}\,\sigma_{xy}}\right) \\ & + \frac{I_{\text{bkg}}}{2} \operatorname{erfc}\left(\frac{x - x_0}{\sqrt{2}\,\sigma_{xy}}\right). \end{aligned} \tag{C9}$$

We fitted Eq. C9 to the fluorescence intensity profiles (Fig. 7b) and optimized the following parameters: $I_{\text{plsm}}$, $I_{\text{gcx}}$, $I_{\text{abl}}$, $I_{\text{bkg}}$, $\sigma_{xy}$, $\lambda$, $x_0$, $\Delta_{\text{endo}}$, $\Delta_{\text{gcx}}$.

### Supplementary Note: Fine-tuning active contours for glycocalyx maps

For generating glycocalyx maps (Online Methods), we used `skimage.segmentation.active_contour` command in Python's Scikit-image library[11]. The final contour shape depends on the algorithm's parameters of which we adjusted $\alpha$, $\beta$, $w_{line}$, $w_{edge}$, max_px_move, and boundary_condition. To find values of these parameters, which result, we used the following steps as a guide:

- Starting with all parameters having their default values, change $w_{line}$ gradually to find its value, at which the contour is attracted to the glycocalyx fluorescence band.
- Keep an eye on the distance between the optimized contour's points. If it is several times smaller or larger than the pixel size, increase $\alpha$.
- If the contour is attracted by the fluorescence inside the vessel lumen ( e.g., labeled red blood cells; $\mathrm{WGA_{plsm}} + \mathrm{WGA_{RBC}}$ in Fig. 1c), decrease $w_{line}$. If this doesn't help, increase $w_{edge}$ gradually to attract the contour to the edge of the vessel. Typically, we set $w_{edge} > 0$ in venules, where the glycocalyx fluorescence band is dim and in some regions is dimmer than the fluorescence in the middle of the vessel lumen.
- If the optimized contour is not smooth enough (e.g., repeats the shape of the initial contour), increase $\beta$.
- If one of the end nodes of the contour is "pulled" to a very bright fluorescence in the glycocalyx band, shorten the initial contour by removing the end point or moving it closer its neighbor point.

#### Supplementary Note: Statistical models

Our modelling strategy involved (i) fitting Bayesian multilevel generalised linear regression models (GLMs), (ii) sampling posteriors via Markov Chain Monte Carlo (MCMC), and (iii) analysing test statistics from the posterior distributions. Model choice depended on measurement type, prior knowledge, and model performance. Following the Bayesian Workflow [3], we began with simple models and iteratively modified components, priors, and hyperparameters to balance descriptive accuracy with computational stability. Final models are detailed below. Instructions for reproducing our analysis are available at https://github.com/teddygroves/glycocalyx/blob/main/README.md.

##### Bayesian posterior sampling

Our results describe posterior distributions of a test statistics (TS — a function of model parameters, e.g., treatment effect differences). We summarized these with means and 2.5–97.5% inter-quantile ranges, representing values consistent with model assumptions. For example, reporting the 1% quantile indicates a 1% model-implied probability of lower values (within small Monte Carlo error). As with any analysis, interpretation depends on the model's validity and accurate posterior sampling.

##### Model assessment and validation

We evaluated models quantitatively with leave-one-out log predictive density[12] and qualitatively with graphical prior and posterior predictive checks. We validated computation using standard Hamiltonian Monte Carlo diagnostics, including $\hat{R}$ [13], checks for divergent transitions [1], EBFI, tree depth, and effective sample size ratios. All models showed $\hat{R} \approx 1$ with no divergences or other signs of failure.

##### Modelling approach

We used linear regression models to represent our measurements after transformation to an unconstrained space. For a transformed measurement $y_u$ with distribution $p$, the expected value was modelled as $E_p(y_u|x, \alpha, \beta) = \eta$, where $\eta$ is a linear predictor based on covariates $x$, intercept parameters $\alpha$ and possibly slope parameters $\beta$. To incorporate structural knowledge (e.g., similarities among mice within treatment or vessel categories), we introduced some hierarchical intercept parameters. To accommodate heteroskedasticity, in some cases we used distributional models, where the measurement error as well as the expected value of each transformed measurement is modelled.

Below we outline the mathematical features of each model used in the final analysis. Full implementations are available in the repository (link above). The equations below use the following notation conventions:

- Superscript, e.g. "$^{global}$" in "$\alpha^{\text{global}}$": label for a parameter.
- Subscript, e.g. "$_k$" in "$\alpha_k^{\text{vessel type}}$": index of a non-scalar variable.
- Tilde, e.g. "$\sim$" in "$\alpha^{\text{global}} \sim N(0, 0.5)$": according to the model, the random variable on the left has the probability distribution on the right.
- $N(\mu, \sigma)$: the normal distribution with expected value $\mu$ and standard deviation $\sigma$.
- $N^+(\sigma)$: the normal distribution centered and truncated below at zero, with standard deviation $\sigma$.
- $ST^+(\nu, \sigma)$: the student-T distribution centered and truncated below at zero, with degrees-of-freedom $\nu$ and standard deviation $\sigma$.

##### Model A

Model A was used for analysis of glycocalyx maps, shown in Fig. 3b,d of the main text and Extended data Fig. 4. We used a linear regression on logarithmic scale, with random intercepts per vessel type and per lectin:vessel type interaction class. Code implementing this model can be found at https://github.com/

teddygroves/glycocalyx/blob/main/src/glycocalyx/model.stan. It's description in tilde notation is as follows:

$$
\begin{aligned}
\ln y &\sim N(\hat{\ln y}, \sigma) \\
\hat{\ln y}_{kv} &= \alpha^{\text{global}} + \alpha_k^{\text{vessel type}} + \alpha_v^{\text{lectin:vessel type}} \\
\sigma &\sim N^+(1) \\
\alpha^{\text{global}} &\sim N(0, 0.5) \\
\alpha^{\text{vessel type}} &\sim N(0, \tau^{\text{vessel type}}) \\
\alpha^{\text{lectin:vessel type}} &\sim N(0, \tau^{\text{lectin:vessel type}}) \\
\tau^{\text{vessel type}} &\sim N(0, 0.5) \\
\tau^{\text{lectin:vessel type}} &\sim N(0, 0.5)
\end{aligned}
\tag{D1}
$$

#### Model B

Model B was used for the analysis of plasma hyaluronan concentration shown in Fig. 5a of the main text. We used a distributional model to account for treatment-dependent heteroskedasticity. For the expected measurement value we used a linear regression on logarithmic scale, with non-random effects for treatment (Enzyme or Saline) and random effects per mouse. For the expected measurement error we used a linear regression on logarithmic scale with a non-random treatment effect. Code implementing this model can be found at https://github.com/teddygroves/glycocalyx/blob/main/scripts/plasma_hyaluronan.py. The description in tilde notation is as follows:

$$
\begin{aligned}
\ln y &\sim N(\hat{\ln y}, \sigma) \\
\hat{\ln y}_{tm} &= \alpha^{y\text{global}} + \alpha_t^{y\text{treatment}} + \alpha_m^{y\text{mouse}} \\
\sigma_t &= \alpha^{\sigma\text{global}} + \alpha_t^{\sigma\text{treatment}} \\
\alpha^{y\text{global}} &\sim N(4.27, 1.73) \\
\alpha^{y\text{treatment}} &\sim N(0, 2.63) \\
\alpha^{y\text{mouse}} &\sim N(0, \tau^{y\text{mouse}}) \\
\alpha^{\sigma\text{global}} &\sim N(0, 1) \\
\alpha^{\sigma\text{treatment}} &\sim N(0, 1) \\
\tau^{\text{ymouse}} &\sim N(0, 0.3)
\end{aligned}
\tag{D2}
$$

Note that the priors for $\alpha^{y\text{global}}$ and $\alpha^{y\text{treatment}}$ are default values chosen by bambi to be weakly informative based on the scale of the measurements.

#### Model C

Model C was used to analyse FRAP data shown in Fig. 5 of the main text. We used separate linear regression models to each set of parameters after transforming each parameter value $y_i$ to an unconstrained value $y_i^u \in \mathbb{R}$. Each model had non-random treatment effects and random effect for mouse and ROI. chosen by bambi. Code implementing this model can be found at https://github.com/teddygroves/glycocalyx/blob/main/scripts/gcx_frap.py. The description in tilde notation is as follows:

$$
\begin{aligned}
y^u &\sim N(\hat{y^u}, \sigma) \\
\hat{y^u}_{tmr} &= \alpha^{\text{global}} + \alpha_t^{\text{treatment}} + \alpha_m^{\text{mouse}} + \alpha_r^{\text{ROI}} \\
\alpha^{\text{global}} &\sim N(m^{\alpha^{\text{global}}}, s^{\alpha^{\text{global}}}) \\
\alpha^{\text{treatment}} &\sim N(0, \sigma^{\alpha^{\text{treatment}}}) \\
\alpha^{\text{mouse}} &\sim N(0, \tau^{\text{mouse}}) \\
\alpha^{\text{roi}} &\sim N(0, \tau^{\text{roi}}) \\
\tau^{\text{mouse}} &\sim N(0, s^{\tau^{\text{mouse}}}) \\
\tau^{\text{roi}} &\sim N(0, s^{\tau^{\text{roi}}})
\end{aligned} \tag{D3}
$$

Note that in this equation the terms $m^{\alpha^{\text{global}}} and s^{\alpha^{\text{global}}}$ etc indicate constant default prior values chosen by bambi based on the scale of the measurements. These values are not shown here as they varied per parameter.

The transformations used were as follows:

| Measured parameter | Natural Range | $y^u$ |
|---|---|---|
| $\alpha$ | $[0, 1]$ | $logit(\alpha)$ |
| $c0$ | $\mathbb{R}^+$ | $\ln c0$ |
| $k1$ | $\mathbb{R}^+$ | $\ln k1$ |
| $k2$ | $\mathbb{R}^+$ | $\ln k2$ |
| $r1$ | $\mathbb{R}^+$ | $\ln r1$ |
| $r2$ | $\mathbb{R}^+$ | $\ln r2$ |
| $I_{\text{plasma}}$ | $\mathbb{R}^+$ | $\ln I_{\text{plasma}}$ |

#### Model D

Model D was used to analyze glycocalyx fluorescence intensities shown in Fig. 7bc of the main text. Linear regression on logarithmic scale with non-random effects for treatment, vessel type and treatment:vessel type interaction, as well as random effects for mouse and vessel. Code implementing this model can be found at https://github.com/teddygroves/glycocalyx/blob/main/scripts/abs_gcx_intensity.py. The description in tilde notation is as follows:

$$
\begin{aligned}
\ln y &\sim N(\hat{\ln} y, \sigma) \\
\hat{\ln} y_{tkmv} &= \alpha^{\text{global}} + \alpha_t^{\text{treatment}} + \alpha_k^{\text{vessel type}} + \alpha_{tk}^{\text{treatment:vessel type}} + \alpha_m^{\text{mouse}} + \alpha_v^{\text{vessel}} \\
\sigma &\sim ST^+(4, 1) \\
\alpha^{\text{global}} &\sim N(0, 2.5) \\
\alpha^{\text{treatment}} &\sim N(0, 0.5) \\
\alpha^{\text{vessel type}} &\sim N(0, 0.5) \\
\alpha^{\text{mouse}} &\sim N(0, \tau^{\text{mouse}}) \\
\alpha^{\text{vessel}} &\sim N(0, \tau^{\text{vessel}}) \\
\tau^{\text{mouse}} &\sim N(0, 0.5) \\
\tau^{\text{vessel}} &\sim N(0, 0.5)
\end{aligned} \tag{D4}
$$

#### Model E

Model D was used to analyze parameters obtained by fitting line-profile of fluorescence shown in Fig. 7gh of the main text. We used a distributional model due to availability of information about the likely accuracy of the measurements in the form of covariates $x$. For the expected measurement value we used a linear regression on log scale with non-random treatment effects and random effects per mouse. For the expected measurement error we used a linear regression on log scale with a non-random, positive-constrained effect for the measurement error predictor. Code implementing this model can be found at https://github.com/teddygroves/glycocalyx/blob/main/scripts/gcx_thickness.py. The description in tilde notation is as follows:

$$
\begin{aligned}
\ln y &\sim N(\hat{\ln y}, \sigma) \\
\hat{\ln y_{tm}} &= \alpha_t^{y\text{treatment}} + \alpha_m^{y\text{mouse}} \\
\sigma_t &= \alpha^{\sigma\text{global}} + \beta^{\sigma} \ln x \\
\alpha^{y\text{treatment}} &\sim N(m^{\alpha^{y\text{treatment}}}, 0.5) \\
\alpha^{y\text{mouse}} &\sim N(0, \tau^{y\text{mouse}}) \\
\alpha^{\sigma\text{global}} &\sim N(0, 1) \\
\beta^{\sigma} &\sim N^{+}(2) \\
\tau^{y\text{mouse}} &\sim N(0, s^{\tau^{y\text{mouse}}})
\end{aligned}
\tag{D5}
$$

Similarly to the FRAP model, in this equation the terms $m^{\alpha^{y\text{treatment}}}$ and $s^{\tau^{y\text{mouse}}}$ indicate default, data-dependent values.

[2] Thomas Broggini, Jacob Duckworth, Xiang Ji, Rui Liu, Xinyue Xia, Philipp Mächler, Iftach Shaked, Leon Paul Munting, Satish Iyengar, Michael Kotlikoff, Susanne J van Veluw, Massimo Vergassola, Gal Mishne, and David Kleinfeld. Long-wavelength traveling waves of vasomotion modulate the perfusion of cortex. *Neuron*, May 2024.

[3] Andrew Gelman, Aki Vehtari, Daniel Simpson, Charles C. Margossian, Bob Carpenter, Yuling Yao, Lauren Kennedy, Jonah Gabry, Paul-Christian Bürkner, and Martin Modrák. Bayesian workflow, 2020.
